# Reconstitution of monoterpene indole alkaloid biosynthesis in genome engineered *Nicotiana benthamiana*

**DOI:** 10.1101/2021.08.12.456143

**Authors:** Quentin M. Dudley, Seohyun Jo, Delia Ayled Serna Guerrero, Monika Chhetry, Mark A. Smedley, Wendy A. Harwood, Nathaniel H. Sherden, Sarah E. O’Connor, Lorenzo Caputi, Nicola J. Patron

## Abstract

Monoterpene indole alkaloids (MIAs) are a diverse class of plant natural products that include a number of medicinally significant compounds. We set out to reconstitute the pathway for strictosidine, a key intermediate of all MIAs, from central metabolism in *Nicotiana benthamiana*. A disadvantage of this host is that its rich background metabolism results in the derivatization of some heterologously produced molecules. We used transcriptomic analysis to identify glycosyltransferases that were upregulated in response to biosynthetic intermediates and produced plant lines with targeted mutations in the genes encoding them. Expression of the early MIA pathway in these lines produced a more favorable product profile. Strictosidine biosynthesis was successfully reconstituted, with the best yields obtained by the co-expression of 14 enzymes, of which a major latex protein-like enzyme (MLPL) from Nepeta (catmint) was critical for improving flux through the iridoid pathway. The removal of endogenous glycosyltransferases did not impact the yields of strictosidine, highlighting that the metabolic flux of the pathway enzymes to a stable biosynthetic intermediate minimizes the need to engineer the endogenous metabolism of the host. The production of strictosidine *in planta* expands the range of MIA products amenable to biological synthesis.

## Introduction

The application of synthetic biology approaches to engineering plant systems has facilitated advances in the control and expression of biosynthetic pathways, enabling plants to serve as an alternative biochemical production chassis^1,2^. A relative of tobacco, *N. benthamiana*^3,4^ has emerged as a favored species for plant-based production of pharmaceutical proteins^5^ and metabolic pathway reconstitution^2^. Successes include gram-scale production of triterpenoids^6^ and milligram scale production of etoposides^7^. However, several studies have reported the accumulation of unintended side products, presumably produced by the off-target activities of endogenous *N. benthamiana* enzymes such as oxidases and glycosylases^8–18^. Although the activity of endogenous enzymes has been exploited in some studies to produce novel molecules^8^ or to compensate for the lack of a known enzyme^19^, derivatization of molecules for the most part is a disadvantage, reducing the potential purity and yield of the target compound.

Monoterpene indole alkaloids (MIAs) are a large group of plant-produced natural products of which over 3000 have been identified^20^. This class of molecules includes many medicinally valuable compounds used to treat addiction, heart conditions, dementia, pain, cancer, malaria, and diabetes. The best characterized MIA-producing plant is *Catharanthus roseus* (Madagascar periwinkle), which makes over 130 MIAs including the bioactive vinblastine and vincristine, which are used as chemotherapies. However, these valuable molecules are present in low concentrations in *C. roseus* (0.0005% dry weight)^21^, which limits their availability. Mass cultivation of *C. roseus* cells is feasible, but a cell line that consistently produces these anticancer molecules has yet to be reported^22^. Although methods for transient expression^23^ and stable genetic transformation^24^ of *C. roseus* plants have been reported, genetically engineering the native plant host to increase yields of these compounds remains technically difficult. Furthermore, the structural complexity of many MIAs means chemical synthesis is often challenging^25,26^. Consequently, alternate routes for production are desirable and the recent discovery of missing steps in the vinblastine pathway^27,28^ makes pathway reconstruction in a heterologous host an increasingly attractive option.

Achieving the production of therapeutically useful amounts of MIAs requires pathway engineering to maximize metabolic flux through the early parts of the pathway. Strictosidine is the last common biosynthetic intermediate from which all 3000+ known MIAs derive. Reconstitution of its ~11 step biosynthetic pathway in microorganisms can require extensive tuning of enzyme expression conditions and strain optimization^29,30^; for example, poor expression of geraniol 8-hydroxylase (G8H) has hampered strictosidine production in yeast^30^. Obtaining useful yields of molecules such as vinblastine, which would require the expression of a further 16+ enzymes beyond strictosidine, is likely to require significant metabolic engineering effort, though yeast has recently been engineered to produce ajmalicine via the genomic integration of 29 expression cassettes, demonstrating the potential for heterologous reconstruction of plant natural product biosynthetic pathways effort^31^.

While yeast remains a promising host for heterologous expression of metabolic pathways, plant-derived proteins frequently require less optimization and engineering for successful expression in a plant host. Expression in *N. benthamiana* was used by Miettinen and coworkers to reconstitute the *C. roseus* iridoid pathway, enabling elucidation of the remaining four missing steps of the pathway^14^. However, they encountered a metabolic bottleneck midway through the 13-step pathway requiring reconstitution in two phases with the latter part requiring the provision of an intermediate substrate (iridotrial) in order to obtain strictosidine^14^. Moreover, the rich endogenous metabolism of *N. benthamiana* acted on the early, hydrophobic intermediates of strictosidine biosynthesis to produce a variety of derivatized, “dead-end” products.

Here we successfully engineer *de novo* production of strictosidine in *N. benthamiana.* We identify endogenous glycosyltransferases that derivatize early pathway intermediates and demonstrate that fewer derivatives accumulate in plant lines with loss-of-function-mutations in the genes encoding these enzymes. We utilize additional genes that enhance the production of nepetalactol and enable high levels of strictosidine to be produced from the central metabolism *of N. benthamiana* without the need for supplementation of any metabolite precursors or intermediates. Overall, this study demonstrates the potential of *N. benthamiana* as a bioproduction chassis for small molecules.

## Methods

### Construction of expression constructs

Binary vectors for *Agrobacterium tumefaciens*-mediated transient expression (agroinfiltration) were either assembled by cloning coding sequences amplified from *C. roseus* cDNA into the pEAQ-HT-DEST1 vector (GenBank GQ497235, **Supplementary Table S1**)^32^ or assembled into expression constructs using the plant modular cloning (MoClo) toolkit^33^. For the latter, coding sequences were either synthesized (Twist Bioscience, San Francisco, CA) removing any native chloroplast transit peptide sequence as well as any instances of BpiI, BsaI, BsmBI and SapI recognition sites by the introduction of synonymous mutation or amplified from *C. roseus* cDNA by PCR with overhangs containing BpiI (BbsI) recognition sites. Amplicons were cloned into pUAP1 (Addgene #63674) as previously described^34^ resulting in level 0 parts (Addgene# 177019 - 177032) flanked by an inverted pair of BsaI recognition sites that produce overhangs compatible with the phytobrick assembly standard^35^ (**Supplementary Table S2)**. These level 0 parts were assembled into level 1 acceptors in a one-step cloning reaction^34^ with level 0 parts encoding CaMV 35s promoter (CaMV35s) and 5’ UTR from tobacco mosaic virus (TMV), and, when required, a synthetic chloroplast transit peptide sequence (**Supplementary Table S3**) to alter subcellular localization.

### Transient expression in *N. benthamiana*

*N. benthamiana* plants were grown in a controlled environment room with 16 hr light, 8 hr hours dark with room at 22 °C, 80% humidity, and ~200 μmol/m^2^/s light intensity. Electrocompetent *A. tumefaciens* GV3101 (MoClo vectors) or LBA4404 (pEAQ vectors) were transformed with the binary plasmid encoding the gene of interest and an individual colony was used to inoculate liquid LB medium containing antibiotics 50 μg/mL rifampicin, 20 μg/mL gentamicin, and 100 μg/mL carbenicillin (MoClo vectors) or 50 μg/mL rifampicin, 100 μg/mL streptomycin, and 50 μg/mL kanamycin (pEAQ vectors). Overnight saturated cultures were centrifuged at 3,400 *x g* for 30 min at room temperature and cells were resuspended in infiltration medium (10 mM 2-(N-morpholino)ethanesulfonic acid (MES) pH 5.7, 10 mM MgCl2, 200 μM 3’,5’-Dimethoxy-4’-hydroxyacetophenone (acetosyringone)) and incubated at room temperature for 2-3 hours with slow shaking. All resuspended cultures were diluted to 0.8 OD_600nm_ (MoClo) or 0.5 OD_600nm_ (pEAQ) and mixed in equal ratios as dictated by the experimental condition. For MoClo vectors, a separate *A. tumefaciens* strain encoding a gene expressing the P19 suppressor of gene silencing from Tomato Bushy Stunt Virus (TBSV) previously shown to increase heterologous expression was included in every infiltration^32^. Healthy plants (29-37 days old) with 3-4 fully expanded true leaves were infiltrated on the abaxial side of the leaf using a 1 mL needleless syringe and grown for five days in an MLR-352-PE plant growth chamber (Panasonic Healthcare Co, Oizumi-Machi, Japan) with 16 hr light, 8 hr hours dark at 22 °C and 120-180 μmol/m^2^/s light intensity. All chemical compounds were purchased from Sigma-Aldrich (St. Louis, MO).

### Metabolite extraction

Five days post-infiltration, 100-300 mg of infiltrated *N. benthamiana* leaf tissue was collected in 1.5 mL microcentrifuge tubes and flash-frozen in liquid nitrogen. Leaf tissue was lyophilized overnight using a VirTis BenchTop SLC freeze dryer (SP Industries, Stone Ridge NY, USA) set to −49 °C and 300 mTorr. Samples were then ground to powder using a 3 mm tungsten carbide bead (Qiagen Cat. No. / ID: 69997) on a TissueLyser II (Qiagen, Hilden, Germany) set to 20 Hz for 20 sec. Lyophilized leaf tissue was extracted with 70% methanol + 0.1% formic acid (1:100, w:v). The solvent contained 10 μM of harpagoside (Extrasynthese, Genay, France) as an internal standard. The extractions were performed at room temperature for 1 hr, with 10 min sonication and 50 min constant shaking. Samples were centrifuged at 17,000 *x g* for 10 min to separate the debris and filtered through 0.2 μm PTFE disk filters before ultra-high performance liquid chromatography-mass spectrometry (UPLC/MS) analysis.

### Metabolite analysis

UPLC/MS analysis was performed on an Impact II qTOF mass spectrometer (Bruker) coupled to an Elute UPLC (Bruker) chromatographic system. Chromatographic separation was carried out on a Phenomenex Kinetex column XB-C18 (100 × 2.10 mm, 2.6 μm particle size) kept at 40 °C and the binary solvent system consisted of solvent A (H2O + 0.1% formic acid) and solvent B (acetonitrile). Flow rate was 600 μL/min. The column was equilibrated with 99% A and 1% B. During the first minute of chromatography, solvent B reached 5%. Then a linear gradient from 5% B to 40% B in 5 min allowed the separation of the compounds of interest. The column was then washed at 100% B for 1.5 min and re-equilibrated to 1% B. Injection volume was 2 μL. Mass spectrometry was performed both in positive and negative ion mode with a scan range *m/z* 100-1000. The mass spectrometer was calibrated using sodium formate adducts. The source settings were the following: capillary voltage 3.5 kV, nebulizer 2.5 Bar, dry gas 11.0 L/min, dry temperature 250 °C. Data analysis was performed using the Bruker Data Analysis software. Quantification of 7-deoxyloganic acid (7-DLA), loganin, loganic acid and strictosidine was based on calibration curves generated using pure compounds. Loganin and loganic acid were purchased from Sigma. 7-deoxyloganic acid and strictosidine were synthesized as previously described^36,37^. The standards were diluted in 70% methanol + 0.1% formic acid to give nine calibrants with concentrations between 40 nM and 10 μM. A linear response was observed for all compounds in this range of concentrations (R^2^>0.993). Putative identification of metabolites was based on the acquisition of high-resolution mass spectrometry data to determine the best fit elemental composition using the Data Analysis software (Bruker).

### Expression analysis of infiltrated plants

Agroinfiltration experiments were performed as described above with four sets of pEAQ vectors: (1) “low geraniol” (GFP, CrGES), (2) “high geraniol” (GFP, CrDXS, CrGGPPS.LSU, CrGES), (3) “nepetalactol” (GFP, CrDXS, CrGGPPS.LSU, CrGES, CrG8H, Cr8HGO, CrISY), and (4) an infiltration control (GFP only) (**Supplementary Table S4**). Leaf tissue from three biological replicates of each condition and a mock infiltrated control were collected five days postinfiltration and flash-frozen in liquid nitrogen. Total RNA was isolated using the RNeasy plant mini kit (Qiagen) with recombinant DNase I (Roche) treatment. Libraries were constructed on a Sciclone® G3 NGSx workstation (PerkinElmer, Waltham, MA) using the TruSeq RNA protocol v2 (Illumina 15026495 Rev.F). RNA quality was assessed using the Quant-iT™ RNA Assay Kit (Life technologies/Invitrogen Q-33140). 1 μg of RNA was purified to extract polyadenylated mRNA using biotin beads, fragmented, and primed with random hexamers to produce first strand cDNA followed by second-strand synthesis to produce ds cDNA. Sample ends were blunted and A-tailed and indexing adapters with corresponding T-overhangs were ligated. The ligated products were size-selected and unligated adapters were removed using XP beads (Beckman Coulter A63880). Samples were amplified with a cocktail of PCR primers to the adapters. The insert size of the libraries was verified using the LabChipGX (PerkinElmer 5067-4626) and DNA High Sensitivity DNA reagents (PerkinElmer CLS760672). Equimolar quantities of each library were pooled, five libraries per pool, and sequenced on one lane of a HiSeq 2500 generating 100 base pair paired-end reads. Data analysis was performed using a web-based Galaxy interface (https://galaxy.earlham.ac.uk)^38^. The *N. benthamiana* draft genome assembly v0.5 http://benthgenome.qut.edu.au/^39^ was expanded to include the genome and plasmids of Agrobacterium C58 (Genbank AE007869.2, AE007870.2, AE007871.2, and AE007872.2) as well as the pEAQ vectors used for infiltration (**Supplementary Table S1**). All short reads from RNA-seq were mapped to the expanded genome using hisat2 v2.1.0 default parameters. Assembled transcripts were generated using Stringtie v1.3.3.1. Transcripts from n=15 experimental samples were consolidated using Stringtie merge v1.3.3. All short reads were again aligned to the expanded genome (this time with the merged transcriptome as a reference) using hisat2 v2.1.0. Differential expression tables were generated using DESeq2 v2.11.40.1. Transcripts from the merged transcriptome were translated using transdecoder and the longest coding sequences annotated using phmmer v3.1v2 with the Swiss-Prot protein database (accessed May 2018) as reference.

### Phylogenetic reconstruction of UDP-glycosyltransferases (UGTs)

*N. benthamiana* transcripts encoding sequences annotated as Family 1 Glycosyltransferases (GT1, Protein family (Pfam) PF0201 or PF3033) were analyzed resulting in 77 sequences >327 amino acids and including an intact Plant Secondary Product Glycosyltransferase (PSPG) box. Protein sequences of 107 UGTs from *Arabidopsis thaliana* were obtained from the *A. thaliana* cytochrome P450, cytochrome b5, P450 reductase, b-glucosidase, and glycosyltransferase site (http://www.p450.kvl.dk) as in described^40^. An additional 9 UGT sequences previously reported to have activity on geraniol or other iridoid substrates from *Actinidia deliciosa* (kiwifruit)^41^, *Camellia sinensis* (tea)^42^, *C. roseus^43^, Gardenia jasminoides* (Cape jasmine)^44^, *Sorghum bicolor^45^, Vitis vinifera* (grape)^46,47^ were also included. Sequences are listed in (**Supplementary Table S5**) along with GenBank accession numbers. The 193 sequences were aligned using MUSCLE 3.8.425 (Edgar 2004) and a phylogenetic tree with 100 bootstraps was generated using RAxML version 8.2.11^48^ within the Geneious program. Phylogenetic trees were visualized using Interactive Tree Of Life (iTOL)^49^.

### Cas9-mediated targeted mutagenesis by Agrobacterium-mediated transformation

Binary vectors expressing Cas9 and single guide RNAs (sgRNAs) for the desired targets were assembled using the plant modular cloning toolkit^33^ as previously described^50^. In brief, primers encoding the desired sgRNA spacer (**Supplementary Table S6**) were used to PCR amplify the sgRNA stem extension scaffold reported by Chen and coworkers^51^. The resulting PCR amplicon was assembled with a level 0 part encoding either an Arabidopsis U6-26 promoter (AtU6-26 Addgene#68261) or a *N. benthamiana* U6 promoter (NbU6-1 Addgene#185623 and NbU6-2 Addgene#185624) (**Supplementary Table S6**). U6 promoters from *N. benthamiana* were identified by sequence homology to Arabidopsis U6-26 and efficacy was confirmed by transient infiltration as previously described^50^. The resulting Level 1 constructs were assembled with synthetic genes conferring resistance to kanamycin and for constitutive expression of Cas9 (**Supplementary Figure S1)**. Efficacy of sgRNAs was confirmed by transient infiltration as previously described^50^. The resulting constructs were transformed into the hypervirulent *A. tumefaciens* strain AGL1 for plant transformation. An individual colony was used to inoculate 10 mL LB medium with antibiotics (50 μg/mL kanamycin and 50 μg/mL rifampicin). Overnight saturated cultures were centrifuged at 3,000 *x g* for 10 min at room temperature and cells were resuspended in 10 mL MS medium with 100 μM acetosyringone and optical density OD_600_ adjusted to 0.6-0.8. *N. benthamiana* was transformed as previously reported^52^ with slight modifications. Young leaves were harvested from 4-week-old, non-flowering plants and surface sterilized. 1-2 cm squares were inoculated with *Agrobacterium* for five minutes at room temperature, blotted dry on sterile filter paper and placed abaxial side down on co-cultivation media pH 5.8 containing MS basal salt^53^, Gamborg’s B5 vitamins^54^, 3% (w/v) sucrose, 0.59 g/L MES hydrate, 6 g/L agarose, 6-benzylaminopurine (BAP, 1.0 mg/L) and naphthaleneacetic acid (NAA, 0.1 mg/L). Explants were co-cultivated for 3 days under white fluorescent light (16 hour light/8 hours dark) at 22±2 ^o^C then transferred to selection medium containing MS basal salts, Gamborg’s B5 vitamins, 3% (w/v) sucrose, 0.59 g/L MES hydrate, 6 g/L Agargel, 6-Benzylaminopurine (BAP, 1.0 mg/L) and naphthaleneacetic acid (NAA, 0.1 mg/L), 100 mg/L kanamycin and 320 mg/L ticarcillin. Explants were sub-cultured onto fresh selection medium at 14-day intervals. Putative transgenic shoots were cultured on rooting media containing MS salts with vitamins (half strength), supplemented with 1 mg/L indole-3-butyric acid, 0.5% sucrose, 0.3% Gelrite, 100 mg/L kanamycin and 320 mg/L ticarcillin. Plantlets were transferred to sterile peat blocks (Jiffy7) in Magenta™ vessels (Sigma), before being transplanted to peat-based compost (90% peat, 10% grit, 4 kg/m^3^ dolomitic limestone, 0.75 kg/m^3^ compound fertilizer (PG Mix™), 1.5 kg/m^3^ controlled-release fertilizer (Osmocote Bloom)) and transferred to a glasshouse.

### Cas9-mediated targeted mutagenesis using RNA viruses and mobile sgRNAs

A TRV2 plasmid vector SPDK3888 (Addgene #149276) was gratefully received from Savithramma Dinesh-Kumar and Dan Voytas. This was modified by adding AarI restriction sites and a lacZ cassette for selection producing pEPQD0KN0750 (Addgene#185627). Spacer sequences for selected targets were incorporated into pEPQD0KN0750 by Golden Gate assembly into AarI sites as previously described^55^ (**Supplementary Table S7**). Constructs were transformed into *A. tumefaciens* GV3101 and an individual colony was used to inoculate LB medium containing 50 μg/mL rifampicin, 20 μg/mL gentamicin, and 50 μg/mL kanamycin and grown overnight shaking at 28 °C. Saturated cultures were centrifuged and resuspended in infiltration medium as described above except that cultures were diluted to 0.3 OD_600nm_ and Agrobacterium strains were mixed in equal ratio with a strain containing pTRV1 (Addgene #148968). Seed of transgenic *N. benthamiana* plants constitutively expressing Cas9 (Cas9 Benthe 193.22 T5 Homozygous) were gratefully received from Dan Voytas and grown for 6 weeks in a greenhouse at 22±2°C. Plants were infiltrated as described above and allowed to grow for 13 weeks before samples of leaf tissue were taken from two different stems (designated A and B).

### Identification of lines with induced mutations

Samples of 40-60 mg leaf tissue were collected from T_0_ plants generated by agrobacterium-mediated stable transformation or from leaves sampled from two stems of plants infiltrated with RNA viruses expressing mobile sgRNAs. Genomic DNA was isolated as previously described^50^. Target loci were amplified using a proof-reading polymerase (Q5® High-Fidelity DNA Polymerase, New England Biolabs, Ipswich, MA) and primers flanking the target sites (**Supplementary Table S8**). Amplicons were sequenced by Sanger sequencing (Eurofins, Luxembourg). Amplicons with multiple peaks that suggested the presence of either genetic chimerism, heterozygous or biallelic mutations, were resolved by cloning amplicons into Zero Blunt™ TOPO™ (Thermo Fisher, Waltham, MA) or pGEM®-T Easy (Promega, Madison, WI) followed by sequencing of plasmids isolated from 10-15 colonies. For plants generated by agrobacterium-mediated stable transformation, the T-DNA copy number of T_0_ plants was estimated by ddPCR as previously described^56^ using primers to *nptII* and the single-copy reference gene Rdr1^57^. T_1_ seeds from single copy plants with homozygous, heterozygous or biallelic mutations at target locations were collected and grown for subsequent analyses. For plants generated using RNA viruses and mobile sgRNAs, seeds (designated as E_1_) were harvested from individual pods at the distal end of stems in which mutations were detected. T_1_ and E_1_ were grown, and the genotype of each target loci was confirmed as above.

## Results

### Metabolites of the early iridoid pathway are derivatized by the endogenous metabolism of *N. benthamiana*

Strictosidine is derived from the terpenoid secologanin. Secologanin belongs to the iridoid class of monoterpenes, which are derived from the oxidation, reductive cyclization and substantial derivatization of geraniol. After formation, secologanin undergoes a condensation and cyclization with tryptamine to form strictosidine. Previous studies aiming at the heterologous expression of low molecular weight terpenoid biosynthesis pathways in *N. benthamiana* have reported the accumulation of derivatized biosynthetic intermediates^8–14^. To investigate derivatization in the early iridoid pathway steps, we transiently expressed the early strictosidine pathway enzymes by co-infiltrating *N. benthamiana* with *A. tumefaciens* strains containing plasmids encoding pathway steps up to *cis-trans* nepetalactol (**Figure 1**). To enhance the pool of geraniol pyrophosphate (GPP) substrate from the plastidial 2-C-methyl-D-erythritol 4-phosphate/1-deoxy-D-xylulose 5-phosphate (MEP/DOXP) pathway, we included a bifunctional geranyl/geranylgeranyl pyrophosphate synthase from *Picea abies* (PaGPPS) and 1-deoxy-D-xylulose 5-phosphate synthase (DXS) from *C. roseus* (CrDXS), both targeted to the plastid. To produce geraniol, these were co-expressed with geraniol synthase from *C. roseus* (CrGES), previously shown to enable the heterologous production of geraniol in plants^58^. We then added the first three dedicated strictosidine pathway steps: geraniol 8-oxidase (CrG8H), 8-hydroxygeraniol oxidoreductase (CrGOR) and iridoid synthase (CrISY) to produce *cis-trans* nepetalactol and perfomed high-resolution mass spectrometry of transiently infected *N. benthamiana* leaves to identify the enzymatic products. As reported by Dong and coworkers^13^, we found that transient expression of CrDXS, PaGPPS and CrGES produces a range of nonvolatile glycosylated and oxidized derivatives of geraniol (**Figure 1** and **Supplementary Table S9**). The addition of later pathway steps, through to *cis-trans* nepetalactol further modified the profile of derivatized products. In general, fewer derivatives accumulated as more pathway steps were added, suggesting that the strong, constitutive expression enabled the pathway enzymes to outcompete endogenous substrates (**Figure 1**).

**Figure 1.**
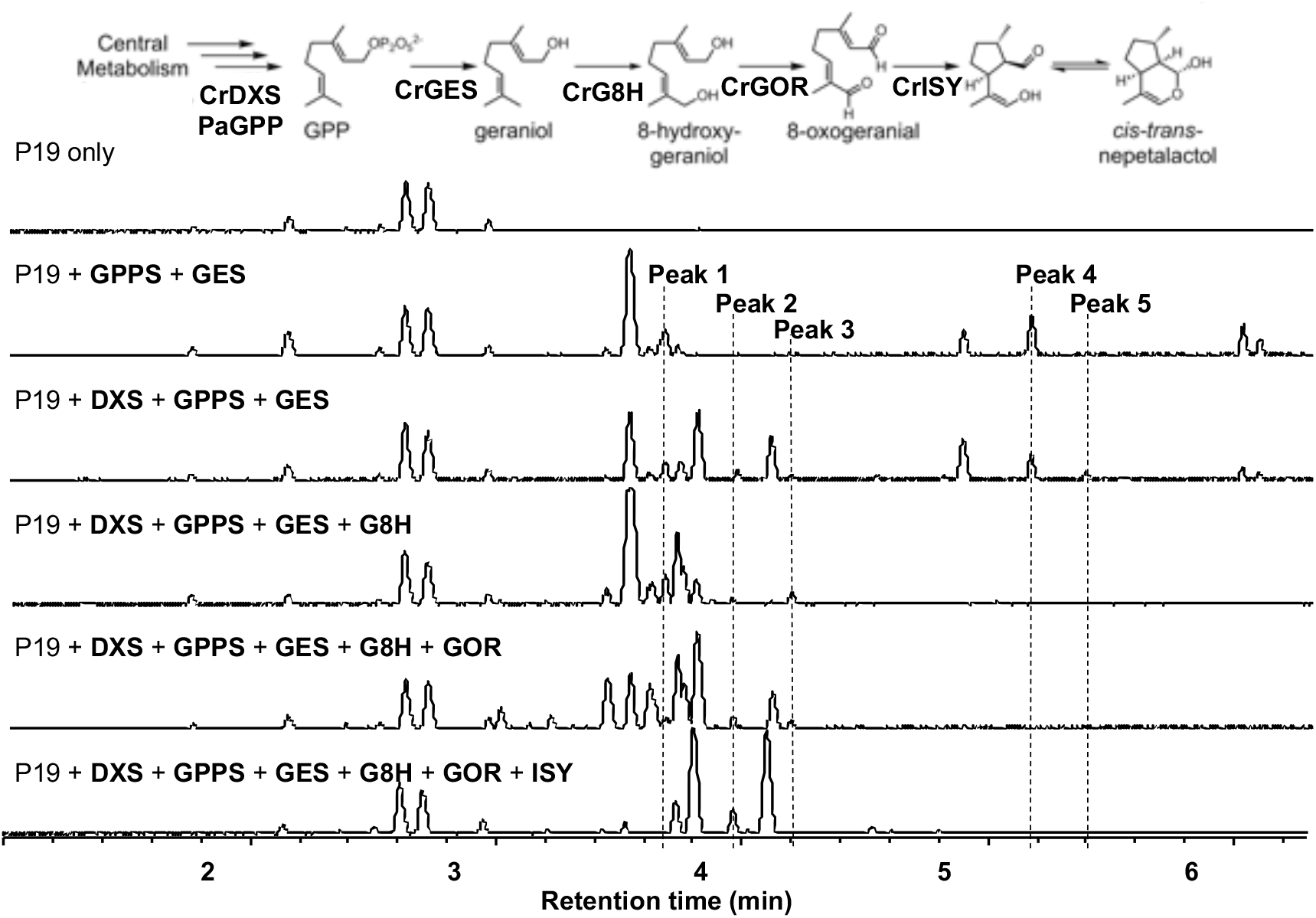
Metabolites of the early iridoid pathway are derivatized by endogenous enzymes from *N. benthamiana.* A common modification was the addition of pentose and hexose sugars e.g., Peak 1, hexosyl hydroxygeraniol [M+HCOOH-H], 3.84 minutes, *m/z* 377.1817; Peak 2, hexosyl hydroxycitronellal [M+HCOOH-H], 4.12 minutes, *m/z* 379.1974; Peak 3, trihexosyl geranic acid [M+HCOOH-H], 4.35 minutes, *m/z* 699.2703); Peak 4, pentosyl hexosyl geraniol [M+HCOOH-H], 5.32 minutes, *m/z* 493.2288; Peak 5, acetyl dihexosyl geraniol [M-H], 5.454 minutes, *m/z* 519.2445; DXS, 1-deoxy-D-xylulose 5-phosphate synthase; GPPS, geranyl diphosphate synthase; GES, geraniol synthase; G8H, geraniol 8-oxidase; GOR, 8-hydroxygeraniol oxidoreductase; ISY, iridoid synthase.

One of the common features of derivatization was the addition of pentose and hexose sugars, which suggests that Family 1 uridine diphosphate-dependent glycosyltransferases (UGTs) are involved in the modification of these foreign metabolites. We hypothesized that many of these UGTs are likely to be involved in the biosynthesis of endogenous secondary metabolites used, for example, in defense. Alternatively, these UGTs may be involved in the detoxification of a variety of metabolites. We therefore considered that introducing loss-of-function mutations into the genes encoding them would be unlikely to have substantial impacts on the development of plants grown in the controlled environments used for agroinfiltration. However, while large numbers of UGTs can be readily identified by searching the genomes of all vascular plants for appropriate homologues, predicting the substrate specificities, and identifying individual genes responsible for modifying specific metabolites is challenging^59^.

### The expression of endogenous UGTs is altered by the early iridoid pathway

To investigate if changes in gene expression correlated with the chemical modifications observed, we conducted transcriptome analysis of leaf samples infiltrated with the iridoid pathway. As a significant transcriptional response to agroinfiltration has previously been observed^60^, we compared changes in expression to both non-infiltrated controls and to control plants infiltrated with a constitutively expressed green fluorescent protein (GFP) (see methods). We observed that the expression of some UGTs increased in response to infiltration, while others increased specifically in response to expression of the pathway to geraniol or to nepetalactol (**Supplementary Table S10**). We also performed a phylogenetic analysis of all UGTs identified in the *N. benthamiana* genome (see methods), investigating their relationship to those found in *Arabidopsis thaliana* and to previously characterized plant UGTs known to be active on geraniol and nepetalactol substrates (**Figure 2** and **Supplementary Figure S2).**

**Figure 2.**
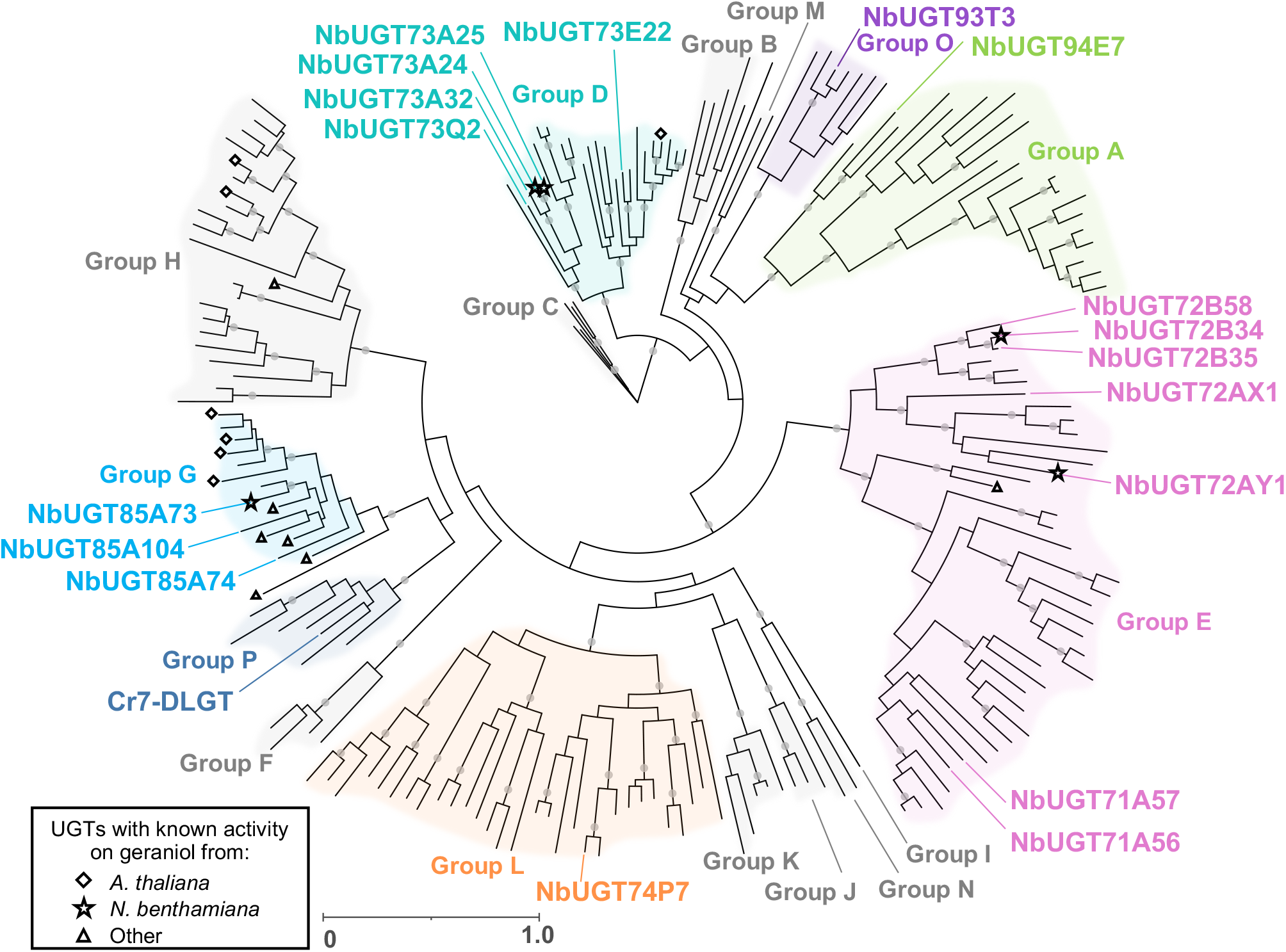
Maximum likelihood (RAxML) phylogenetic comparison of 193 Family 1 UDP-glycosyltransferases (UGTs) from *N. benthamiana* and *A. thaliana* and nine UGT sequences previously shown to be active on geraniol or iridoid substrates. Groups A-P are annotated according to the nomenclature used by Caputi *et al.* 2012^40^. Labeled taxa indicate enzymes in which Cas9-mediated targeted mutations were subsequently introduced. Filled gray circles at nodes indicate bootstrap supports >95. Scale bar represents the number of substitutions per site. A tree with all taxa and bootstrap values is provided in Supplementary Figure S2.

### Production of *N. benthamiana* lines with Cas9-mediated targeted mutations in UGTs

We hypothesized that one way to improve the *N. benthamiana* chassis would be to remove these endogenous UGTs by introducing mutations into the genes encoding them. Using both the expression profiles and the predicted substrate selectivity, we selected genetic targets for Cas9-mediated targeted mutagenesis. We then constructed 8 plasmids targeting a total of 25 UGTs (**Table 1**). We first designed three binary vectors for the stable transformation of wild-type *N. benthamiana* plants (pEPQDPKN0720, pEPQDPKN0724, pEPQDPKN0361). These constructs encoded sgRNAs targeting genes encoding UGTs in Group D, E and G, previously shown to be active on geraniol^61,62^ as well as uncharacterized UGTs from the same groups with the logic that they may be active on the same substrate (**Table 1 and Supplementary Figures S3-5**). As the UGTs targeted by each construct were closely related, some sgRNAs were able to target more than one gene. Consequently, each construct encoded six sgRNAs targeting the first or second exon of either four Group D UGTs, five Group E UGTs or three Group G UGTs (**Table 1 and Supplementary Figures S3-5**). These constructs were delivered to wild type plants and the genotypes of regenerated transgenic (T_0_) plants were determined. We discarded lines in which the genotype was unclear or indicated potential genetic chimerism and selected lines with homozygous, heterozygous or biallelic mutations at target locations. The genotypes were confirmed in T_1_ plants, together with the presence or absence of the T-DNA. All 18 sgRNAs introduced mutations in at least one T_0_ plant. Notably, we were unable to recover any plants with frameshift mutations in both NbUGT72B58 and NbUGT72B35, however, we were able to recover lines with mutations in each of these genes in combination with other Group E UGTs (**Table 1**). We selected six T_1_ plants with mutations in different combinations of 12 individual genes (**Table 1**). All lines were morphologically normal with no obvious changes in growth and development and maturing at the same time as control lines.

**Table 1.**
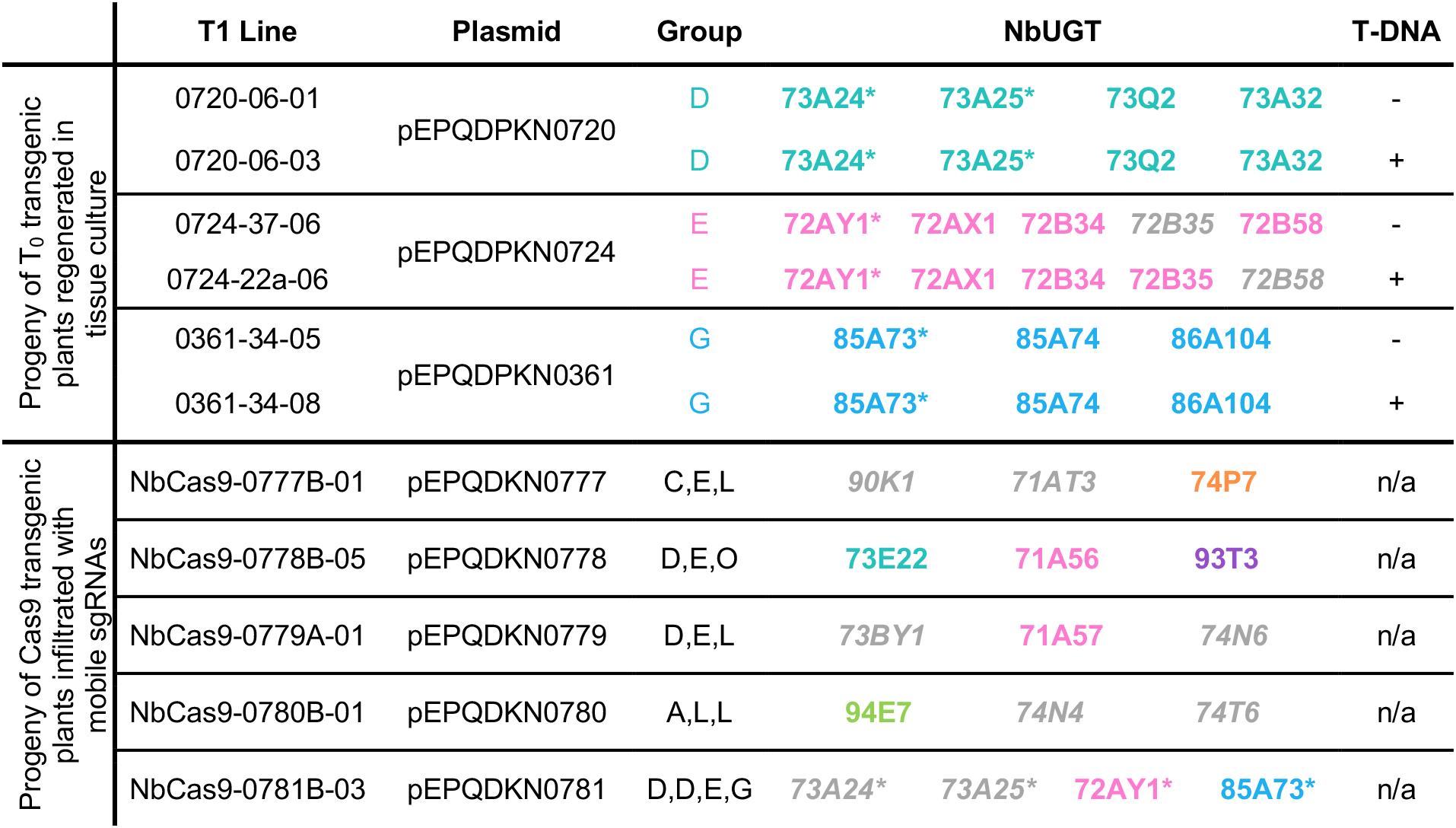
Lines of *N. benthamiana* with Cas9-induced mutations in UGTs. Asterisks indicate that activity on geraniol has previously been reported^62^. NbUGTs in gray italics were wild-type or contained small in-frame deletions unlikely to inactivate the enzyme. T-DNA represents the presence (+) or absence (-) of the T-DNA delivered in this experiment

We also designed and built five TMV-based viral vectors encoding mobile sgRNAs using the methods previously described by Ellison and colleagues^55^. Four vectors contained mobile sgRNAs targeting UGTs that were either strongly upregulated following infiltration (pEPQDKN0777), strongly upregulated in response to expression of the geraniol pathway (pEPQDKN0778) or the nepetalactol pathway (pEPQDKN0779) or had high normalized counts following expression of the nepetalactol pathway (pEPQDKN0780) (**Table 1, Supplementary Figure S6 and Supplementary Table S10**). A final vector encoded three mobile sgRNAs targeting four UGTs known to be active on geraniol^62^ (pEPQDKN0781) (**Table 1** and **Supplementary Figure S6**). These vectors were infiltrated into transgenic plants constitutively expressing Cas9. Progeny seeds (designated as E_1_) were collected from pods that formed on stems in which mutations were detected at the target sites (see methods). The genotypes of E_1_ plants were analyzed, again selecting lines with homozygous, heterozygous or biallelic mutations at target locations. Overall, the mobile sgRNA method was less efficient, with eight out of fifteen mobile sgRNAs introducing mutations at their target. This resulted in the selection of five E_1_ lines, each with mutations in one, two or three target UGTs (**Table 1** and **Supplementary Figure S6**). As above, there were no changes in growth or development.

### Targeted mutagenesis identifies UGTs active on geraniol and prevents accumulation of glycosylated derivatives

To investigate if the introduction of mutations in UGTs would reduce the presence of geraniol derivatives, mutated plants of which the genotypes of all target loci had been confirmed were infiltrated with strains of agrobacterium encoding pathway enzymes to produce either geraniol or *cis-trans* nepetalactol. Samples were analyzed for derivatives by UHPLC/MS as previously discussed and the metabolic profile was compared to parental lines (wild type and Cas9 transgenic lines) and to the progeny of a non-transgenic, non-mutated control line regenerated in tissue culture and grown in identical conditions.

Plants with mutations in the Group A UGT, NbUGT94E7, did not accumulate trihexosyl geranic acid (*m/z* 699.2713, [M+HCOOH-H]) following expression of pathway enzymes to produce either geraniol or nepetalactol (**Figure 3 and Supplementary Figures S7 and S8**). The presence of acetyl dihexosyl geraniol (*m/z* 519.2445, [M-H]) was also eliminated in samples in which pathway enzymes to produce nepetalactol were expressed. In addition, three lines with mutations in the Group G NbUGT85A73 in combination with either NbUGT72AY1 (Group E) or NbUGT85A74 and NbUGT85A104 (Group G) accumulated less or no hexosyl hydroxygeraniol (*m/z* 377.1817, [M+HCOOH-H]), hexosyl hydroxycitronellal (*m/z* 379.1974, [M+HCOOH-H]), pentosyl hexosyl geraniol (*m/z* 493.2288, [M+HCOOH-H]) or acetyl dihexosyl geraniol (*m/z* 519.2445, [M-H]) (**Figure 3 and Supplementary Figures S7 and S8**). Interestingly, these peaks were only absent in samples in which pathway enzymes to produce nepetalactol were expressed.

**Figure 3.**
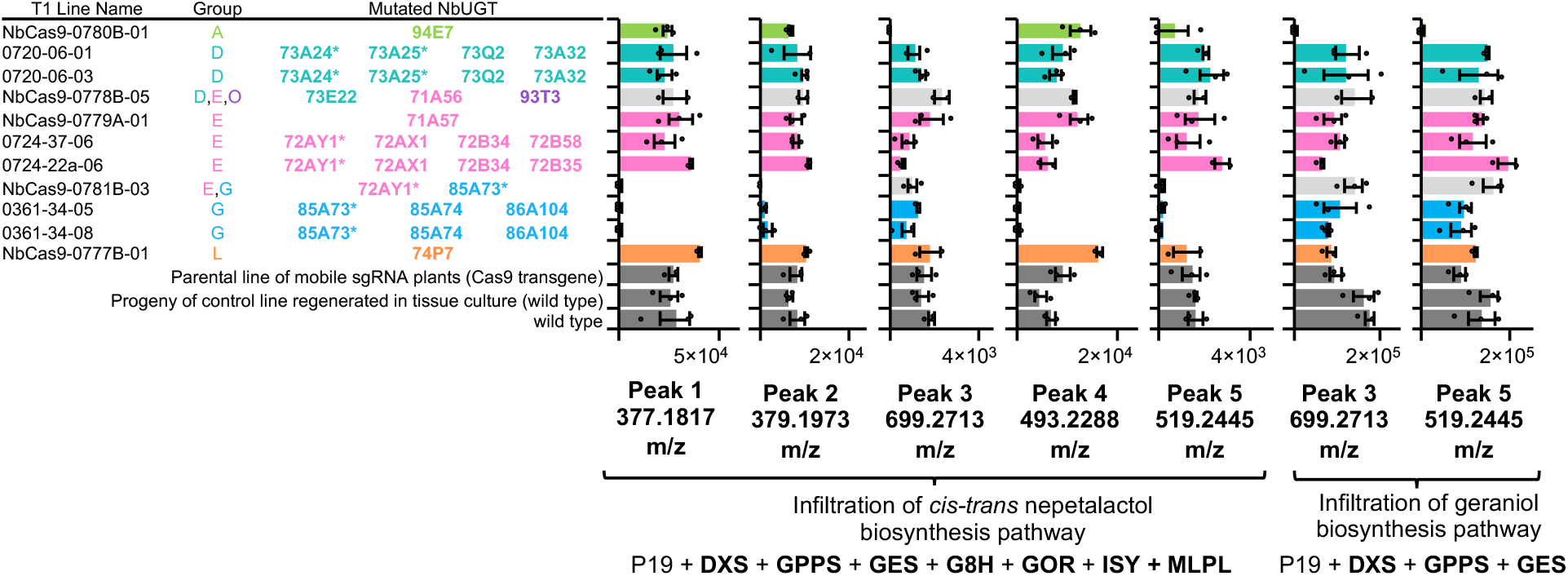
Mutation of NbUGT94E7 (Group A) or NbUGT85A73 (Group G) eliminates the accumulation of specific derivative compounds. The assigned identities of peaks 1-5 are provided in Figure 1. Asterisks indicate that activity on geraniol has previously been reported.

### MLPL1 from *Nepeta mussinii* is essential for 7-DLA production in *N. benthamiana*

We then focussed on extending the iridoid pathway with the goal of reconstituting the pathway through to the central strictosidine intermediate. A previously reported attempt encountered a metabolic bottleneck requiring the infiltration of iridotrial in order to obtain strictosidine^i4^. The biosynthetic pathway for production of *cis-trans* nepetalactone in the genus *Nepeta* shares the secoiridoid pathway from *C. roseus* up to the stereoselective reduction of 8-oxogeranial to an enol intermediate by the NADPH-dependent iridoid synthase (ISY)^63^. It has recently been shown that in *Nepeta* ISY works in combination with either nepetalactol-related short-chain dehydrogenase-reductases (NEPS) or a major latex protein-like enzyme (MLPL) to control the stereoselectivity of the ring closure^64,65^. We therefore hypothesized that addition of the MLPL from *Nepeta mussinii* (a.k.a *Nepeta racemosa*), which is specific for the stereochemistry found in strictosidine, would enhance flux through the pathway in *N. benthamiana.* Recent efforts at reconstitution in yeast^31^ and cell-free^66^ systems have also indicated that this enzyme substantially enhances yield.

Infiltration of all pathway enzymes to produce 7-DLA without MLPL did not produce a peak for 7-DLA (expected *m/z* 359.1349) or acylated 7-DLA (expected *m/z* 401.1447), consistent with previous reports^14^ (**Figure 4**). However, the inclusion of MLPL produced a clear peak of *m/z* 359.1342 which matches the retention time of the 7-DLA standard. Exclusion of 7-deoxyloganetic acid glucosyl transferase (7-DLGT) produces a peak with the same *m/z* but this does not match the retention time of 7-DLA. It is possible that endogenous UGTs are able to use 7-deoxyloganetic acid to produce a glucose ester. As observed with the early pathway, strong, constitutive expression of pathway genes (in this case, 7-DLGT) may outcompete native enzymes resulting in the absence of this putative glucose ester peak in the spectra of the full 7-DLA pathway.

**Figure 4.**
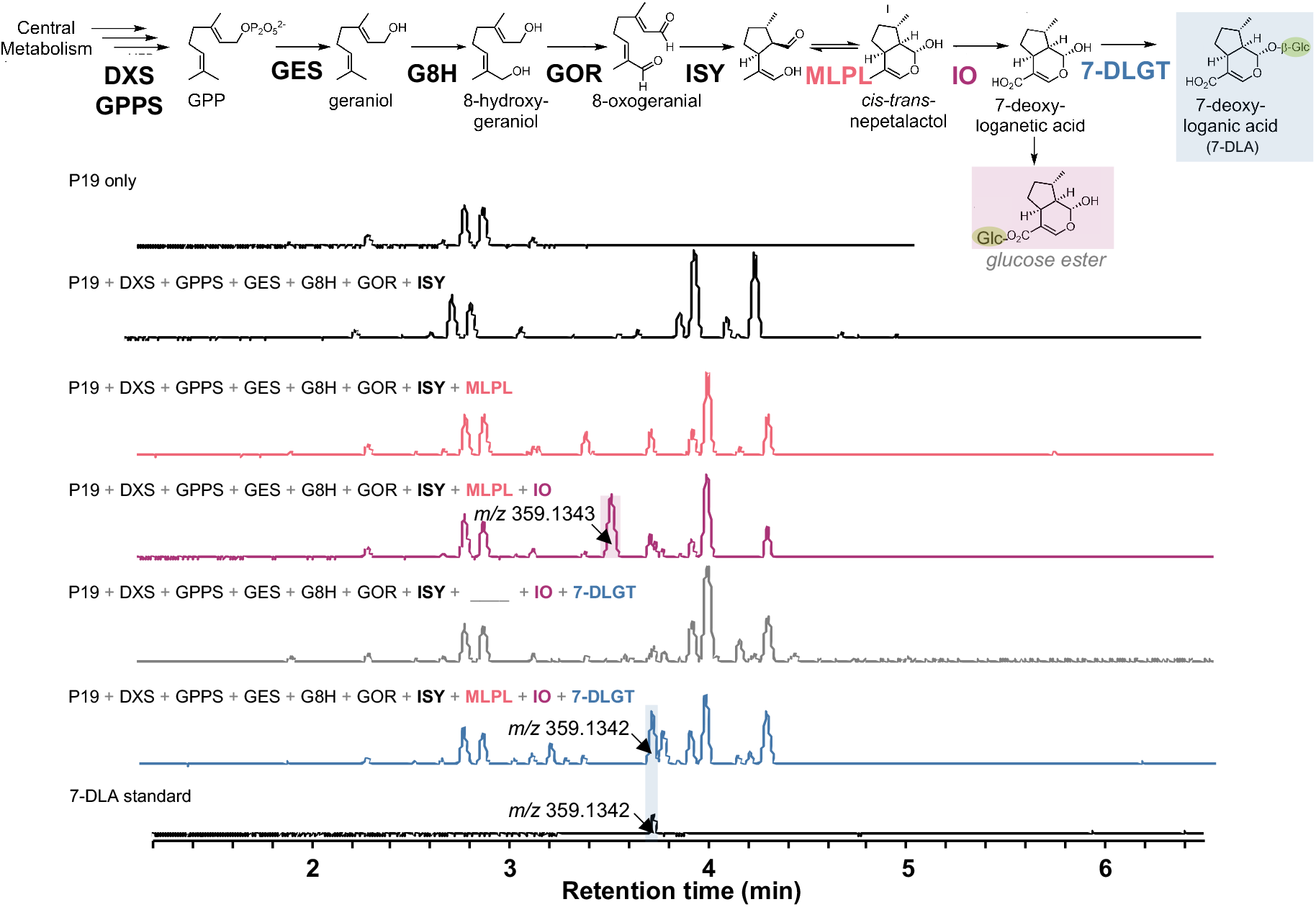
MLPL1 from *Nepeta mussinii* enables 7-deoxyloganic acid production in *N. benthamiana.* Transient expression of all pathway enzymes plus MLPL produces a peak of 359.1342 *m/z* which matches the mass and retention time of a 7-deoxyloganic acid (7-DLA) standard. The reconstituted pathway without DLGT also produces a 359.1343 *m/z* peak but at a different retention time, likely due to endogenous glycosyltransferases *of N. benthamiana* producing the glucose ester of 7-deoxyloganetic acid. DXS, 1-deoxy-D-xylulose 5-phosphate synthase; GPPS, geranyl diphosphate synthase; GES, geraniol synthase; G8H, geraniol 8-oxidase; GOR, 8-hydroxygeraniol oxidoreductase; ISY, iridoid synthase; MLPL, major latex protein–like; IO, iridoid oxidase; 7-DLGT, 7-deoxyloganetic acid glucosyltransferase

### Reconstitution of the complete strictosidine pathway

Building on the experimental conditions that successfully produced 7-DLA, we sequentially added the remaining five pathway enzymes and measured the pathway intermediates after the sequential addition of each gene (**Figure 5A, Supplementary Figure S9-S13**). Co-infiltration of strains expressing enzymes for the entire pathway produces a distinct peak at 531 *m/z* matching the strictosidine standard (**Figure 5C, Supplementary Figure S12**). This pathway configuration produces 4.29 ± 2.00 μM of strictosidine which correlates to 0.23 ± 0.11 mg strictosidine / g dry weight leaf tissue (0.023% DW). The only major biosynthetic intermediate that was observed to accumulate was loganin, suggesting secologanin synthase is a bottleneck step. This contrasts with previous data suggesting that loganic acid O-methyltransferase (LAMT), which has a low substrate affinity (K_m_ = 12.5–14.8 mM)^67,68^, might be a rate-limiting step of the late stages of the secoiridoid pathway. In addition to strictosidine, transient expression of the full pathway also produces a smaller amount of a compound (**Figure 5C, Supplementary Figure S12**) with a mass shift of 86 Da from strictosidine, suggesting that this may be a malonylated derivative of strictosidine produced by endogenous *N. benthamiana* acyltransferases. Transient expression of the strictosidine pathway without GPPS and MLPL (**Figure 5B**) confirms the beneficial effect of these enzymes on strictosidine yield. Addition of MLPL increased strictosidine production >60 fold while supplementation of GPPS improved yield ~5 fold. DXS supplementation did not change the amount of strictosidine produced (**Figure 1**).

**Figure 5.**
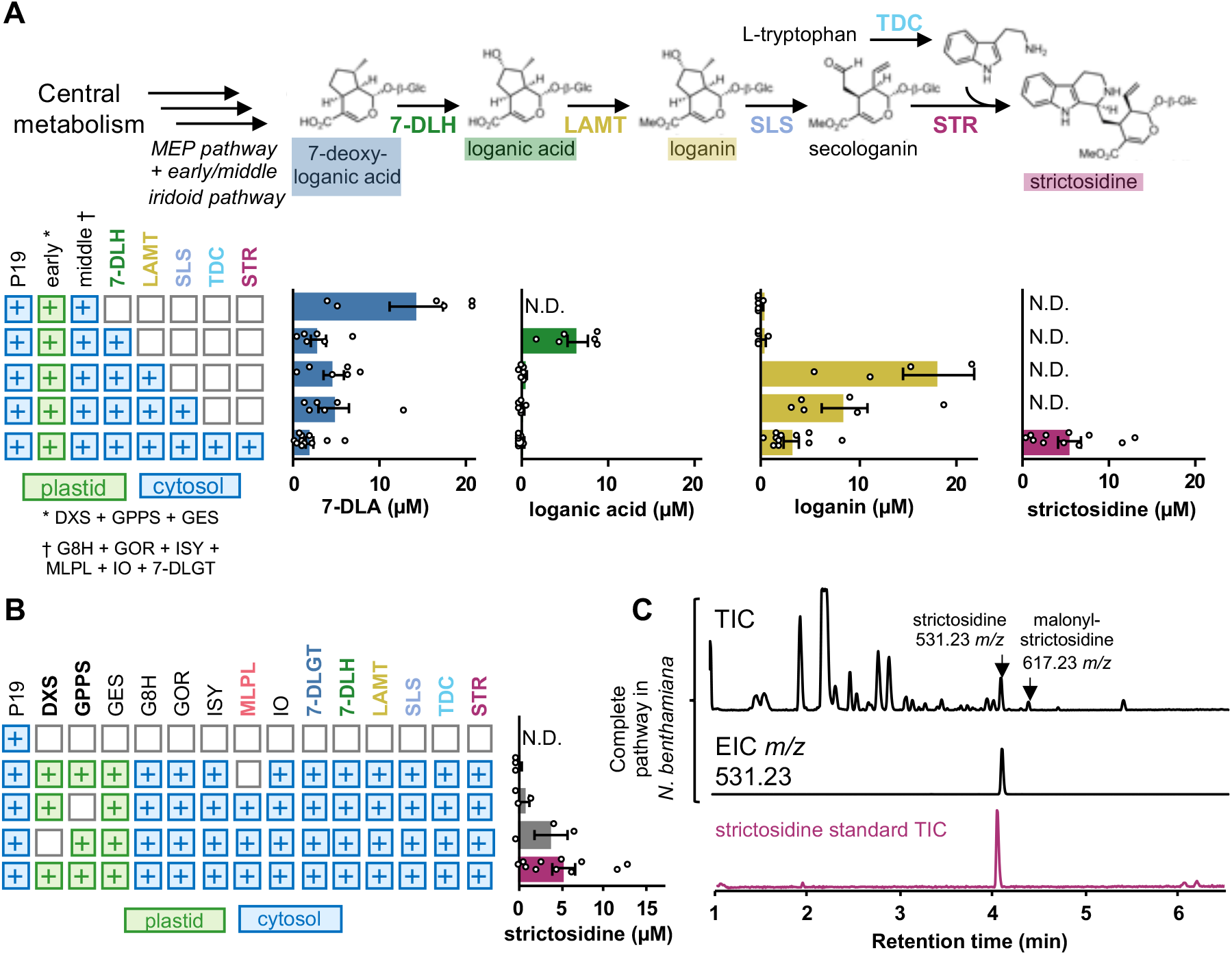
Reconstitution of strictosidine biosynthesis in *N. benthamiana.* (A) Quantification of intermediates and final product strictosidine by UPLC/MS analysis. (B) Absence of MLPL reduces strictosidine production >60 fold while supplementation of GPPS improves yield ~5 fold. (C) the total ion chromatogram of leaf tissue infiltrated with the entire pathway to strictosidine (including DXS, GPPS, and MLPL). The peak at 4.09 min retention time in the total ion chromatogram (TIC) and extracted ion chromatogram (EIC) at 531.2336 *m/z* matches a strictosidine standard. N.D., not detected; 7-DLH, 7-deoxyloganic acid hydroxylase; LAMT, loganic acid O-methyltransferase; SLS, secologanin synthase; TDC, tryptophan decarboxylase; STR, strictosidine synthase. Values and error bars represent the mean and the standard error of n=3 or n=6 biological replicates (independent leaf samples).

We also infiltrated the full pathway into the previously identified lines with mutations in UGT94E7 (Group A) or NbUGT85A73 (Group G), each of which had shown the accumulation of fewer early iridoid derivatives. However, we did not observe any significant changes in the yield of strictosidine or strictosidine by-products (**Supplementary Figure S14)**.

### Mimicking *C. roseus* subcellular compartmentalization maximizes strictosidine production

To compare the effect of chloroplast and cytosolic enzyme localization on yields of 7-DLA and strictosidine production, we added (or removed in the case of GPPS/GES) a transit peptide to each enzyme (**Figure 6**). To enhance flux of isoprenoid precursors in the cytosol, we aimed to alleviate the rate limiting step of the mevalonate pathway by co-infiltrating a truncated 3-hydroxy-3-methylglutaryl-coenzyme A reductase (tHMGR) from oat, previously shown to improve titres of the triterpenoid β-amyrin in *N. benthamiana*^6^. When all enzymes were localized to the cytosol, flux through the secoiridoid pathway was minimal (~90-fold reduction for 7-DLA) while localization of all pathway enzymes to the chloroplast resulted in 5-fold less 7-DLA (**Figure 6A**). The best yields of 7-DLA (**Figure 6A**) and strictosidine (**Figure 6B**) were obtained with chloroplast localization of the early pathway and cytosolic localization of subsequent steps, which mimicked the native localization pattern in *C. roseus*. Production of 7-DLA in the chloroplast is possibly limited by the availability of partner P450 reductases for G8H and iridoid oxidase (IO) or small molecule substrates such as UDP-glucose for 7-DLGT, however, all possible divisions of pathway enzymes between the cytosol and chloroplast still produce 7-DLA indicating that pathway intermediates can cross the chloroplast membrane.

**Figure 6.**
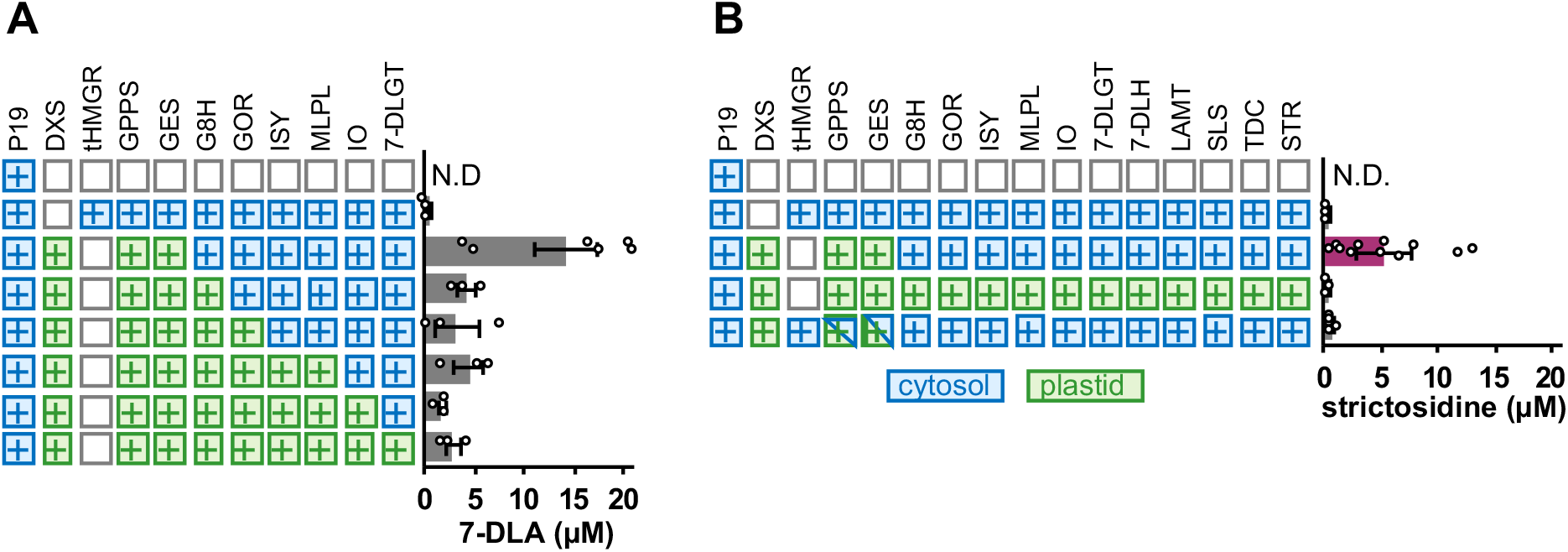
Relocation of 7-DLA (A) and strictosidine (B) biosynthesis genes to the cytosol (blue) or the chloroplast (green) decreases product yield. tHMGR, truncated 3-hydroxy-3-methylglutaryl-coenzyme A reductase. Values and error bars represent the mean and the standard error of n=3 or n=6 biological replicates (independent leaf samples).

## Discussion

In this work, we set out to improve the production of biosynthetic intermediates and products of the iridoid and MIA pathway in *N. benthamiana.* Previous studies have reported that, when *N. benthamiana* is used as a host for heterologous expression of terpenoid natural products, a variety of terpenoid derivatives accumulate, potentially limiting the use of this species as a chassis for the production of some molecule types^8–14^. As found in previous studies, we observed here that early iridoid biosynthetic intermediates were frequently modified by the addition of pentose and hexose sugars suggesting the involvement of *N. benthamiana* Family 1 UGTs (**Figure 1** and **Supplementary Table S9**). Further, while it is already known that the gene expression is significantly affected by agroinfiltration^60^, comparative transcriptomics revealed that the expression of some endogenous UGTs are further modulated by the expression of monoterpene or iridoid metabolic pathways (**Supplementary Table S10)**. This indicates that plants not only respond to the infiltration of *Agrobacterium*, but also to the presence of foreign enzymes and/or metabolites.

Although *N. benthamiana* does not accumulate many monoterpenes, these endogenous UGTs may be involved in the biosynthesis of larger decorated metabolites and are thus able to act promiscuously on the early iridoid intermediates. Alternatively, they may be part of a xenobiotic detoxification mechanism to protect against bioactives produced by pathogens. We reasoned that these enzymes were unlikely to be essential and focused efforts to improve the *N. benthamiana* chassis by identifying and eliminating the activity of these enzymes. We used Cas9-based molecular tools to introduce loss-of-function mutations into 20 UGT target genes (**Table 1** and **Supplementary Figures S3-S6**). We employed two approaches: the production of transgenic plants constitutively expressing Cas9 together with multiple sgRNAs^50^ and the recently described use of a viral vector to transiently express mobile sgRNAs in plants carrying a Cas9 transgene^55^. While we were able to recover lines with mutations in most or all target lines using the former method, the mobile sgRNA method, though less efficient, was significantly less laborious and could more easily be scaled to investigate many target genes.

Relative to control lines, plants with mutations in the Group A UGT, NbUGT94E7, no longer accumulated the peak putatively assigned as trihexosyl geranic acid (*m/z* 699.2713, [M+HCOOH-H]) (**Figure 3 and Supplementary Figures S7 and S8**). This UGT also lacks the GSS motif proposed to be important for defining the loop structure of monoglucosyltransferases^62,69^. We additionally identified that, relative to control lines, four peaks were significantly diminished or absent in plants with mutations in the Group G UGT, NbUGT85A73 (**Figure 3 and Supplementary Figures S7 and 8**). This UGT was previously reported as being active on geraniol^62^. Surprisingly, these peaks were only eliminated in plants infiltrated with all pathway enzymes required to produce nepetalactol, and still remained when pathway genes for expression of geraniol were present. It is likely that hexosyl hydroxygeraniol (*m/z* 377.1817, [M+HCOOH-H]) and hexosyl hydroxycitronellal (*m/z* 379.1974, [M+HCOOH-H]) are produced by glycosylation of 8-hydroxygeraniol and therefore require the expression of geraniol 8-oxidase. However, it is unclear why the accumulation of pentosyl hexosyl geraniol (*m/z* 493.2288, [M+HCOOH-H]) and acetyl dihexosyl geraniol (*m/z* 519.22425, [M-H]) were reduced only when the extended pathway was infiltrated.

Our approach provides proof-of-concept for the application of gene editing approaches to improve *N. benthamiana* as a bioproduction chassis for small, terpenoid type molecules by reducing the accumulation of derivatives. However, we observed that mutations of individual UGTs each reduced the accumulation of minor peaks. To achieve significant impact, the production of lines with mutations in multiple genes may be required to dramatically impact the yield of metabolites of interest. We also found that as the pathway is extended by the coexpression of additional enzymes that fewer derivative peaks are observed (**Figures 1, 4 and 5**). This suggests that the pathway enzymes have high affinity for the substrates and, particularly when expressed from strong promoters, are very likely able to outcompete endogenous enzymes for substrates. Strictosidine is a polar, glycosylated molecule that is likely sequestered in the vacuole for storage and is less prone to derivatization than its hydrophobic iridoid precursors. Indeed, when the pathway for strictosidine was expressed in lines that produced fewer derivatives or early pathway intermediates, there was no significant change in yield compared to expression in wild type *N. benthamiana* (**Supplementary Figure S14)**.

We were able to demonstrate the production of high quantities of strictosidine (0.23 ± 0.11 mg/g DW) from central metabolism in a photosynthetic organism. We also enhanced the production of strictosidine by improving the key cyclization step required for formation of nepetalactol. The plant genus *Nepeta* L., colloquially known as catnip or catmint, uses the early part of the secoiridoid pathway to produce nepetalactones. Recent work revealed that production of *cistrans* nepetalactol is assisted by MLPL in *Nepeta*^65^. The heterologous expression of MLPL from *N. mussinii* to assist with cyclization of the reactive enol product of ISY overcame bottlenecks in the secoiridoid pathway (**Figure 4**) and was critical for enabling heterologous production of strictosidine in *N. benthamiana* (**Figure 5**). MLPL similarly enhances secoiridoid metabolic engineering in microorganisms such as yeast^31^. Additionally, a cell-free *in vitro* one-pot enzyme cascade included MLPL with nine other pathway enzymes, accessory proteins, and cofactor regeneration enzymes to produce ~1 g/L nepetalactone^66^.

The MIA pathway in *C. roseus* is highly compartmentalized across subcellular compartments and cell types. The first committed step is geraniol synthesis (GES), which is localized to the chloroplast of internal phloem-associated parenchyma (IPAP) cells^70^. To increase substrate availability for GES, we co-expressed GPPS from *P. abies* ^71^ which was also utilized by Miettinen and coworkers^14^. Expression of chloroplast-targeted PaGPPS improved yield ~5 fold (**Figure 5**) consistent with previously reported effects on the production of geraniol^13^. Of note, PaGPPS is nearly identical (**Supplementary Figure S15**) to a GPPS from *Picea glauca* shown *in vitro* to produce higher levels of the monoterpene limonene relative to six other GPPS sequences commonly used in terpene metabolic engineering^72^. We also co-expressed DXS from *C. roseus.* Interestingly, CrDXS had relatively little effect on the yield of strictosidine in contrast to previously reported effects of DXS on the production of diterpenoids^10,73^.

Recent efforts to engineer metabolic pathways have found benefits in altering the compartmentalization of pathway enzymes^13,74^. For example, the pathway to produce the cyanogenic glucoside dhurrin was relocated to the chloroplast of *Nicotiana tabacum* where ferredoxin, reduced via the photosynthetic electron transport chain, can serve as an efficient electron donor to the two cytochromeP450s (CYPs) within the pathway^75^. Additionally, the localization of enzymes encoding the late steps of the artemisinin pathway to the chloroplast in *N. tabacum* produced higher levels of artemisinin (800 μg/g DW)^76^ and artemisinic acid (~1200 μg/g DW)^77^ compared to localization within the cytosol (6.8 μg/g DW artemisinin)^78^. This increase is possibly due to the isolation of metabolites from the cytosol where they may both impact viability and be exposed to unwanted derivatization by endogenous glycosyltransferases^11,12,79^. Production of halogenated indican^80^ and vanillin^81^ in *N. benthamiana* also benefited from chloroplast localization. In contrast, a recent report found that production of diterpenoids (typically synthesized in the chloroplast) was dramatically enhanced by co-opting the cytosolic mevalonate pathway to produce GGPP rather than the chloroplast MEP pathway^82^.

In this study, we found that the optimal configuration for reconstructing the pathway to strictosidine within *N. benthamiana* leaves is to match the *C. roseus* localization pattern that utilizes the chloroplast MEP pathway for isoprenoid precursors to produce geraniol and then localizing the remaining pathway enzymes in the cytosol (**Figure 4**). We hypothesize that monoterpene production in the cytosol is limited since GPP (10 carbons) produced by GPPS is also the substrate for the *N. benthamiana* farnesylpyrophosphate synthase (*FPPS*), which produces farnesylpyrophosphate (FPP) (15 carbons). Supporting this hypothesis is data suggesting that all four copies of NbFPPS are upregulated to produce sesquiterpenoid phytosterols and phytoalexins in response to *Phytophthora infestans^83^. A. tumefaciens* also elicits widespread transcriptional remodeling when infiltrated into *N. benthamiana*^60^. This competition between GES and FPPS might also explain the higher levels of geraniol and geraniol derivatives in the plastid reported previously^13^. Future efforts to conditionally inactivate NbFPPS during heterologous production might enable a metabolic engineering strategy that could take advantage of both the plastid and cytosolic route to geraniol production.

In *C. roseus*, geraniol diffuses or is transported from the plastid into the cytosol to react with G8H, which is tethered to the exterior of the ER membrane. The next steps (G8H to deoxyloganic acid hydroxylase (7-DLH)) are active in the cytosol of IPAP cells with two CYPs (G8H and IO) also anchored to the ER membrane^14^. Loganic acid is then transported by NPF2.4/5/6^84^ to epidermal cells where four more enzymes (LAMT to strictosidine synthase (STR)) produce strictosidine ^85,86^. Tryptamine and secologanin are imported into the vacuole, where strictosidine is synthesized and accumulated with export of strictosidine mediated by the transporter NPF2.9^87^. Thus, four pathway CYPs (G8H, IO, 7-DLH and secologanin synthase (SLS)) are likely interfacing with endogenous *N. benthamiana* CYP reductases for electron transfer of NADPH to the CYPs. It is possible that the necessary CYP reductases are less abundant in the chloroplast and thus limit the accumulation of strictosidine in this compartment. It is also possible that the CYP450s are improperly membrane anchored or that higher stromal pH of the chloroplast (~8.0)^88^ inhibits enzyme activity compared to the cytosol (pH ~7.0). We also considered the lack of an additional cofactor for LAMT, S-adenosylmethionine (SAM) to explain the low levels of strictosidine production (relative to 7-DLA) in the chloroplast. However, this small molecule is known to be actively transported into the plastid^89^ and is an essential substrate for ChlM (Mg-protoporphyrin IX methyltransferase) involved in chlorophyll biosynthesis within the chloroplast.

The production of strictosidine *in planta* opens up new avenues to produce a wealth of MIA products using biological synthesis. Although the endogenous metabolism of this species is detrimental to the accumulation of monoterpenes, it enables accumulation of complex molecules. Particularly as new biosynthesis pathways for additional MIAs are discovered (e.g. the anti-addictive compound ibogaine^90^, the antimalarial quinine^91^), the possibility of coupling this work in a plug-and-play manner with downstream biosynthesis modules is an exciting prospect for natural product synthesis.

## Supplementary data

**Supplementary Figure S1.** Design and hierarchical assembly of binary constructs used for Cas9-mediated mutagenesis of *Nicotiana benthamiana.*

**Supplementary Figure S2.** Maximum likelihood (RAxML) phylogenetic comparison of 193 Family 1 UDP-glycosyltransferases (UGTs).

**Supplementary Figure S3.** Cas9-mediated mutagenesis of three Group G UGTs.

**Supplementary Figure S4.** Cas9-mediated mutagenesis of four Group D UGTs.

**Supplementary Figure S5.** Cas9-mediated mutagenesis of five Group E UGTs.

**Supplementary Figure S6.** Supplementary Figure S6. Cas9-mediated mutagenesis of 16 UGTs using mobile sgRNAs

**Supplementary Figure S7.** Abundance of derivative peaks in lines of *N. benthamiana* infiltrated with the geraniol biosynthesis pathway

**Supplementary Figure S8** Abundance of derivative peaks in lines of *N. benthamiana* infiltrated with the nepetalactol biosynthesis pathway.

**Supplementary Figure S9**. Detection of 7-deoxyloganic acid following transient expression of pathway genes in *N. benthamiana.*

**Supplementary Figure S10**. Detection of loganic acid following transient expression of pathway genes in *N. benthamiana.*

**Supplementary Figure S11**. Detection of loganin following transient expression of pathway genes in *N. benthamiana.*

**Supplementary Figure S12.** Detection of strictosidine following transient expression of pathway genes in *N. benthamiana.*

**Supplementary Figure S13.** Detection of putative malonyl-strictosidine following transient expression of pathway genes in *N. benthamiana.*

**Supplementary Figure S14.** Quantification of strictosidine following transient expression of pathway genes in mutated and wild-type *N. benthamiana* by UPLC/MS analysis.

**Supplementary Figure S15.** Amino acid sequence alignment of GPPS enzymes.

**Supplementary Table S1.** Coding sequences cloned into pEAQ-HT-DEST1.

**Supplementary Table S2.** Coding sequences cloned into pUAP1.

**Supplementary Table S3.** Level 1 expression constructs.

**Supplementary Table S4.** Ratios of *A. tumefaciens* strains containing pEAQ plasmid vectors infiltrated into *N. benthamiana.*

**Supplementary Table S5.** Family 1 UDP-glycosyltransferases (UGTs).

**Supplementary Table S6.** Primers used for amplification of sgRNA scaffolds.

**Supplementary Table S7.** Primers used for construction of mobile single guide RNA plasmid vectors.

**Supplementary Table S8.** Primers used for genotyping plants with Cas9-induced mutations.

**Supplementary Table S9**. LC/MS analysis and putative identification of iridoid pathway derivatives produced by pathway expression in *N. benthamiana.*

**Supplementary Table S10.** Expression levels and protein features of *N. benthamiana* UDP-glycosyltransferases (UGTs) selected for mutagenesis.

## Author Contributions

Q.M.D., S.E.O., L.C, and N.J.P. conceived the study. Q.M.D performed DNA assembly and transient expression with assistance from S.J., N.H.S. constructed and performed infiltration experiments with pEAQ vectors. Q.M.D. performed transcriptomic analysis, assembled all vectors for mutagenesis, delivered RNA viruses for mutagenesis, and performed genotyping. M.C. and M.A.S performed all plant tissue culture with supervision from W.A.H. D.A.S.G and L.C. performed metabolite extraction and analysis. Q.M.D., S.E.O., N.J.P., and L.C. wrote the manuscript. S.E.O. and N.J.P obtained funding and provided supervision.

## Acknowledgements

We thank Benjamin Lichman for guidance on working with MLPL. We also thank Nicola Soranzo for help with RNAseq analysis within Galaxy. pL0-AstHMGR was a generous gift from Anne Osbourn. Plasmids pNJB069 (pTRV1), pEE393, and pEE515 along with seeds of *N. benthamiana* expressing SpCas9 were a generous gift from Dan Voytas. Library preparation and sequencing was delivered via the BBSRC National Capability in Genomics and Single Cell Analysis (BBS/E/T/000PR9816) at the Earlham Institute by members of the Genomics Pipelines Group. We also thank Lesley Phillips, Catherine Taylor, and the JIC Horticultural services for help with plant husbandry.

## Data Availability

Plasmids are available from Addgene. Transcriptome data will be made available before publication (submission is in progress).

