## Supplementary data for "Reconstitution of monoterpene indole alkaloid biosynthesis in genome engineered *Nicotiana benthamiana*"

**Supplementary Table S1** Coding sequences cloned into pEAQ-HT-DEST1

| **Enzyme** | | **Organism** | **Key Reference** | **Accession #** |
|---|---|---|---|---|
| CrDXS | 1-deoxy-D-xylulose 5-phosphate synthase | *Catharanthus roseus* (Madagascar periwinkle) | this study | DQ848672 |
| CrGPPS | geranyl diphosphate synthase large subunit | *Catharanthus roseus* (Madagascar periwinkle) | Rai et al Molecular Plant 6(5):1531-49 (2013) | JX417183 |
| CrGES | geraniol synthase | *Catharanthus roseus* (Madagascar periwinkle) | Simkin et al Phytochemistry 85:36-43 (2013) | JN882024 |
| CrG8H | geraniol 8-oxidase | *Catharanthus roseus* (Madagascar periwinkle) | Collu et al FEBS Lett 508(2):215-220 (2001) | AJ251269 |
| CrGOR | 8-hydroxygeraniol reductase | *Catharanthus roseus* (Madagascar periwinkle) | Miettinen et al Nat Comm 5:3606 (2014) and Krithika et al Sci Rep 5 8258 (2015) | KF302069 |
| CrISY | iridoid synthase | *Catharanthus roseus* (Madagascar periwinkle) | Geu-Flores et al Nature 492(7427):138-142 (2012) | JX974564 |
| GFP | green fluorescent protein | *Aequorea victoria* (crystal jelly) | Chalfie et al Science. 263, 802–805 (1994) | AF183395.1 |

**Supplementary Table S2** Coding sequences cloned into pUAP1 (Addgene #63674) resulting in Level 0 standard parts. Asterisk indicates that the sequence encoding the native transit peptide was removed.

| Enzyme | | Organism | Key reference(s) | Accession # | Level 0 plasmid name (AATG-GCTT) | Addgene # |
|---|---|---|---|---|---|---|
| CrDXS* | 1-deoxy-D-xylulose 5-phosphate synthase | *Catharanthus roseus* (Madagascar periwinkle) | *this study* | DQ848672 | pEPQD0CM0065 | 177019 |
| AstHMGR | truncated 3-hydroxy-3-methylglutaryl-coenzyme A reductase | *Avena strigosa* (oat) | Reed *et al. Metabolic Engineering* 42: 185–93 (2017) | KY284573 | pL0-AstHMGR** | n/a - a gift from Anne Osbourn |
| PaGPPS* | geranyl pyrophosphate synthase; geranyl diphosphate synthase | *Picea abies* (Norway spruce) | Schmidt *et al. Plant Physiology* 152(2): 639–55 (2010) | GQ369788 | pEPQD0CM0818 | 177020 |
| CrGES* | geraniol synthase | *Catharanthus roseus* (Madagascar periwinkle) | Simkin *et al. Phytochemistry* 85:36-43 (2013) | JN882024 | pEPQD0CM0063 | 177021 |
| CrG8H | geraniol 8-oxidase; geraniol-10-hydroxylase; CYP76B6 | *Catharanthus roseus* (Madagascar periwinkle) | Collu *et al. FEBS Lett* 508(2):215-220 (2001) | AJ251269 | pEPQD0CM0058 | 177022 |
| CrGOR | 8-hydroxygeraniol oxidoreductase; 10-hydroxygeraniol oxidoreductase; alcohol dehydrogenase 10 | *Catharanthus roseus* (Madagascar periwinkle) | Miettinen *et al. Nat Comm* 5:3606 (2014) and Krithika *et al. Sci Rep* 5 8258 (2015) | KF302069 | pEPQD0CM0059 | 177023 |
| CrISY | iridoid synthase | *Catharanthus roseus* (Madagascar periwinkle) | Geu-Flores *et al. Nature* 492(7427):138-142 (2012) | JX974564 | pEPQD0CM0060 | 177024 |
| NmMLPL | major latex protein–like | *Nepeta mussinii (aka Nepeta racemosa)* | Lichman *et al. Sci Adv* 6(20) eaba0721 (2020) and Lichman *et al. Nat Chem Bio* 15 71-79 (2019) | MT108267.1 | pEPQD0CM0068 | 177025 |
| CrIO | iridoid oxidase; CYP76A26 | *Catharanthus roseus* (Madagascar periwinkle) | Miettinen *et al. Nat Comm* 5:3606 (2014) and Salim *et al. Phytochemistry* 101:23-31 (2014) | KF302066 | pEPQD0CM0061 | 177026 |
| Cr7-DLGT | 7-deoxyloganetic acid glucosyl transferase; UGT709C2 | *Catharanthus roseus* (Madagascar periwinkle) | Miettinen *et al. Nat Comm* 5:3606 (2014) and Asada *et al. Plant Cell* 25(10):4123-4134 (2013) | KF302068 | pEPQD0CM0062 | 177027 |
| Cr7-DLH | 7-deoxyloganic acid hydroxylase; CYP72A224 | *Catharanthus roseus* (Madagascar periwinkle) | Miettinen *et al. Nat Comm* 5:3606 (2014) and Salim *et al. Plant J* 76(5):754:765 (2013) | KF302067 | pEPQD0CM0762 | 177028 |
| CrLAMT | loganic acid O-methyltransferase | *Catharanthus roseus* (Madagascar periwinkle) | Murata *et al. Plant Cell* 20(3):524-542 (2008) | EU057974 | pEPQD0CM0763 | 177029 |
| CrSLS | secologanin synthase; CYP72C | *Catharanthus roseus* (Madagascar periwinkle) | Irmler *et al. Plant J* 24(6):797-804 (2000) | KF309242.1 or KF415117.1 | pEPQD0CM0764 | 177030 |
| CrTDC | tryptophan decarboxylase | *Catharanthus roseus* (Madagascar periwinkle) | de Luca *et al. PNAS* 86(8):2582-2586 (1989) | M25151 | pEPQD0CM0765 | 177031 |
| CrSTR | strictosidine synthase | *Catharanthus roseus* (Madagascar periwinkle) | Pasquali *et al. Plant Mol Biol* 18(6):1121-1131 (1992) | X61932 | pEPQD0CM0766 | 177032 |

**Supplementary Table S3** Level 1 expression constructs assembled from Level 0 parts using the Plant MoClo Toolkit (Addgene Kit #1000000044)

| | (Promoter+5'UTR) *or* (Promoter)+(5'UTR+cTP) | | CDS | 3'UTR + Terminator | Level 1 Acceptor |
|---|---|---|---|---|---|
| **overhangs created by BsaI:** | **GGAG-TACT** | **TACT-AATG** | **AATG-GCTT** | **GCTT-CGCT** | **CGCT-GGAG** |
| pEPQD1CB0104 (P6_35SshortTMV-P19-35Sterm) | pICH51277 | | pICH44022 | pUAP41414 | pICH47781 |
| pEPQD1CB0817 (P1_35SshortTMV-AstHMGR-35Sterm) | pICH51277 | | pL0-AstHMGR | pUAP41414 | pICH47732 |
| pEPQD1CB0107 (P1 _35SshortTMV-cTP_CrDXS2-35Sterm) | pICH41388 | pICH78133 | pEPQD0CM0065 | pUAP41414 | pICH47732 |
| pEPQD1CB0108 (P2_35SshortTMV-PaGPPS1-35Sterm) | pICH51277 | | pEPQD0CM0818 | pUAP41414 | pICH47742 |
| pEPQD1CB0109 (P2_35SshortTMV-cTP_PaGPPS1-35Sterm) | pICH41388 | pICH78133 | pEPQD0CM0818 | pUAP41414 | pICH47742 |
| pEPQD1CB0110 (P3_35SshortTMV-CrGES-35Sterm) | pICH51277 | | pEPQD0CM0063 | pUAP41414 | pICH47751 |
| pEPQD1CB0112 (P3_35SshortTMV-cTP_CrGES-35Sterm) | pICH41388 | pICH78133 | pEPQD0CM0063 | pUAP41414 | pICH47751 |
| pEPQD1CB0113 (P4_35SshortTMV-CrG8H-35Sterm) | pICH51277 | | pEPQD0CM0058 | pUAP41414 | pICH47761 |
| pEPQD1CB0114 (P4_35SshortTMV-cTP_CrG8H-35Sterm) | pICH41388 | pICH78133 | pEPQD0CM0058 | pUAP41414 | pICH47761 |
| pEPQD1CB0115 (P5_35SshortTMV-Cr8HGO/GOR-35Sterm) | pICH51277 | | pEPQD0CM0059 | pUAP41414 | pICH47772 |
| pEPQD1CB0116 (P5_35SshortTMV-cTP_Cr8HGO/GOR-35Sterm) | pICH41388 | pICH78133 | pEPQD0CM0059 | pUAP41414 | pICH47772 |
| pEPQD1CB0117 (P6_35SshortTMV-CrISY-35Sterm) | pICH51277 | | pEPQD0CM0060 | pUAP41414 | pICH47781 |
| pEPQD1CB0118 (P6_35SshortTMV-cTP_CrISY-35Sterm) | pICH41388 | pICH78133 | pEPQD0CM0060 | pUAP41414 | pICH47781 |
| pEPQD1CB0119 (P1 _35SshortTMV-NmMLP-35Sterm) | pICH51277 | | pEPQD0CM0068 | pUAP41414 | pICH47732 |
| pEPQD1CB0120 (P1 _35SshortTMV-cTP_NmMLP-35Sterm) | pICH41388 | pICH78133 | pEPQD0CM0068 | pUAP41414 | pICH47732 |
| pEPQD1CB0121 (P2_35SshortTMV-CrIO-35Sterm) | pICH51277 | | pEPQD0CM0061 | pUAP41414 | pICH47742 |
| pEPQD1CB0122 (P2_35SshortTMV-cTP_CrIO-35Sterm) | pICH41388 | pICH78133 | pEPQD0CM0061 | pUAP41414 | pICH47742 |
| pEPQD1CB0123 (P3_35SshortTMV-CrDLGT-35Sterm) | pICH51277 | | pEPQD0CM0062 | pUAP41414 | pICH47751 |
| pEPQD1CB0124 (P3_35SshortTMV-cTP_CrDLGT-35Sterm) | pICH41388 | pICH78133 | pEPQD0CM0062 | pUAP41414 | pICH47751 |
| pEPQD1CB0767 (P4_35SshortTMV-Cr7DLH-35Sterm) | pICH51277 | | pEPQD0CM0762 | pUAP41414 | pICH47761 |
| pEPQD1CB0768 (P4_35SshortTMV-cTP_Cr7DLH-35Sterm) | pICH41388 | pICH78133 | pEPQD0CM0762 | pUAP41414 | pICH47761 |
| pEPQD1CB0769 (P5_35SshortTMV- CrLAMT-35Sterm) | pICH51277 | | pEPQD0CM0763 | pUAP41414 | pICH47772 |
| pEPQD1CB0770 (P5_35SshortTMV-cTP_ CrLAMT-35Sterm) | pICH41388 | pICH78133 | pEPQD0CM0763 | pUAP41414 | pICH47772 |
| pEPQD1CB0771 (P6_35SshortTMV- CrSLS-35Sterm) | pICH51277 | | pEPQD0CM0764 | pUAP41414 | pICH47781 |
| pEPQD1CB0772 (P6_35SshortTMV-cTP_ CrSLS-35Sterm) | pICH41388 | pICH78133 | pEPQD0CM0764 | pUAP41414 | pICH47781 |
| pEPQD1CB0773 (P1_35SshortTMV- CrTDC-35Sterm) | pICH51277 | | pEPQD0CM0765 | pUAP41414 | pICH47732 |
| pEPQD1CB0774 (P1_35SshortTMV-cTP_ CrTDC-35Sterm) | pICH41388 | pICH78133 | pEPQD0CM0765 | pUAP41414 | pICH47732 |
| pEPQD1CB0775 (P2_35SshortTMV- CrSTR-35Sterm) | pICH51277 | | pEPQD0CM0766 | pUAP41414 | pICH47742 |
| pEPQD1CB0776 (P2_35SshortTMV-cTP_ CrSTR-35Sterm) | pICH41388 | pICH78133 | pEPQD0CM0766 | pUAP41414 | pICH47742 |

**Parts from Plant MoClo Parts Kit (Addgene Kit #1000000047)**
pICH51277 (CaMV 35S short promoter + TMV omega 5'UTR) Addgene#50268
pICH41388 (CaMV 35S short promoter) Addgene#50253
pICH78133 (TMV omega 5'UTR+chloroplast transit peptide RbcS) Addgene#50292
pUAP41414 (CMV 3'UTR+terminator) Addgene#50337
pICH44022 (L0, P19 CDS) Addgene#50330
pICH47732 (L1 P1 acceptor forward) Addgene#48000
pICH47742 (L1 P2 acceptor forward) Addgene#48001
pICH47751 (L1 P3 acceptor forward) Addgene#48002
pICH47761 (L1 P4 acceptor forward) Addgene#48003
pICH47772 (L1 P5 acceptor forward) Addgene#48004
pICH47781 (L1 P6 acceptor forward) Addgene#48005

***Abbreviations***
CaMV = Cauliflower mosaic virus
TMV = Tobacco Mosaic Virus
P19 suppressor of gene silencing (Tomato Bushy Stunt Virus)
cTP = chloroplast transit peptide

**Supplementary Table S4.** Ratios of *A. tumefaciens* strains containing pEAQ plasmid vectors infiltrated into *N. benthamiana*

| | **Low geraniol** | **High geraniol** | **Nepetalactol** | **Infiltration control** |
|---|---|---|---|---|
| **GFP** | 6 units | 1 unit | 1 unit | 1 unit |
| **CrDXS** | | 1 unit | 1 unit | |
| **CrGGPPS.LSU** | | 1 unit | 1 unit | |
| **CrGES** | 1 unit | 1 unit | 1 unit | |
| **CrG8H** | | | 1 unit | |
| **Cr8HGO/GOR** | | | 1 unit | |
| **CrISY** | | | 1 unit | |

**Supplementary Table S5.** Family 1 UDP-glycosyltransferases (UGTs)

| Enzyme | Group code | Organism | Genbank ID | Notes |
|---|---|---|---|---|
| **AdGT4** | **Group G** | *Actinidia deliciosa (kiwifruit)* | AIL51400 | Activity on geraniol reported by Yauk *et al. Plant J.* 80:317–30 (2014) |
| **AtUGT79B1** | **Group A** | *Arabidopsis thaliana* | AB018115, MJP23, 2800-4206 | |
| **AtUGT79B100** | **Group A** | *Arabidopsis thaliana* | AC006193, 78745-80088 | |
| **AtUGT79B111** | **Group A** | *Arabidopsis thaliana* | AC006193, 80861-82219 | |
| **AtUGT79B2** | **Group A** | *Arabidopsis thaliana* | AL161571, 70597-71964 | |
| **AtUGT79B3** | **Group A** | *Arabidopsis thaliana* | AL035602, T29A15, 13430-14791 | |
| **AtUGT79B4** | **Group A** | *Arabidopsis thaliana* | AP000606, MTO24, 63467-64813 | |
| **AtUGT79B5** | **Group A** | *Arabidopsis thaliana* | AC012561, F11F12, 44850-46196 | |
| **AtUGT79B6** | **Group A** | *Arabidopsis thaliana* | AB007644, K19P17, 59557-60918 | |
| **AtUGT79B7** | **Group A** | *Arabidopsis thaliana* | AC006567, T15G18, 32112-33440 | |
| **AtUGT79B8** | **Group A** | *Arabidopsis thaliana* | AC004786, 55469-56797 | |
| **AtUGT79B9** | **Group A** | *Arabidopsis thaliana* | AB007644, K19P17, 55445-56788 | |
| **AtUGT91A1** | **Group A** | *Arabidopsis thaliana* | AC006340, 3826-5238 | |
| **AtUGT91B1** | **Group A** | *Arabidopsis thaliana* | AB026639, 14713-16113 | |
| **AtUGT91C1** | **Group A** | *Arabidopsis thaliana* | AB025613, 26973-28355 | |
| AtUGT89A2 | Group B | *Arabidopsis thaliana* | AL162751, 85361-86758 | |
| AtUGT89B1 | Group B | *Arabidopsis thaliana* | AC016662, F2P9, 86614-88035 | |
| AtUGT89C1 | Group B | *Arabidopsis thaliana* | AC024174, 19642-20949 | |
| AtUGT90A1 | Group C | *Arabidopsis thaliana* | AC005167, F12A24, 24146-26230 | |
| AtUGT90A2 | Group C | *Arabidopsis thaliana* | AC005489, F14N23, 95228-96717 | |
| AtUGT90A4 | Group C | *Arabidopsis thaliana* | AL391149, T9L3, 73006-74878 | |
| **AtUGT73B1** | **Group D** | *Arabidopsis thaliana* | AL021961, F28A23, 51775-53366 | |
| **AtUGT73B2** | **Group D** | *Arabidopsis thaliana* | AL021961, F28A23, 48984-50524 | |
| **AtUGT73B3** | **Group D** | *Arabidopsis thaliana* | AL021961, F28A23, 176825-178270 | |
| **AtUGT73B4** | **Group D** | *Arabidopsis thaliana* | AC006248, F26H6, 11828-13404 | |
| **AtUGT73B5** | **Group D** | *Arabidopsis thaliana* | AC006248, F26H6, 8823-10458 | |
| **AtUGT73C1** | **Group D** | *Arabidopsis thaliana* | AC006282, F13K3, 61903-63378 | Low activity on geraniol (10-40%), reported by Caputi *et al. Chem Eur J* 14(22):6656–62 (2008) |
| **AtUGT73C2** | **Group D** | *Arabidopsis thaliana* | AC006282, F13K3, 64414-65904 | |
| **AtUGT73C3** | **Group D** | *Arabidopsis thaliana* | AC006282, F13K3, 68990-70480 | Very low activity on geraniol (1-10%), reported by Caputi *et al. Chem Eur J* 14(22):6656–62 (2008) |
| **AtUGT73C4** | **Group D** | *Arabidopsis thaliana* | AC006282, F13K3, 66599-68089 | |
| **AtUGT73C5** | **Group D** | *Arabidopsis thaliana* | AC006282, F13K3, 74865-76352 | High activity on geraniol (>40%) reported by Caputi *et al. Chem Eur J* 14(22):6656–62 (2008) |
| **AtUGT73C6** | **Group D** | *Arabidopsis thaliana* | AC006282, F13K3, 71711-73198 | Low activity on geraniol (10-40%), reported by Caputi *et al. Chem Eur J* 14(22):6656–62 (2008) |
| **AtUGT73C7** | **Group D** | *Arabidopsis thaliana* | AL132958, T4D2, 26553-28025 | |
| **AtUGT73D1** | **Group D** | *Arabidopsis thaliana* | AL132958, T4D2, 21804-23327 | |
| **AtUGT71B1** | **Group E** | *Arabidopsis thaliana* | AB025634, MSD21, 21447-22865 | |

| Enzyme | Group code | Organism | Genbank ID | Notes |
|---|---|---|---|---|
| AtUGT71B2 | Group E | *Arabidopsis thaliana* | AB025634, MSD21, 23981-25438 | |
| AtUGT71B5 | Group E | *Arabidopsis thaliana* | AL161541, 37302-38738 | |
| AtUGT71B6 | Group E | *Arabidopsis thaliana* | AB025634, MSD21, 31933-33372 | |
| AtUGT71B7 | Group E | *Arabidopsis thaliana* | AB025634, MSD21, 33809-35296 | |
| AtUGT71B8 | Group E | *Arabidopsis thaliana* | AB025634, MSD21, 37125-38567 | |
| AtUGT71C1 | Group E | *Arabidopsis thaliana* | AC005496, T27A16, 51968-53413 | |
| AtUGT71C2 | Group E | *Arabidopsis thaliana* | AC005496, T27A16, 48813-50237 | Very low activity on geraniol (1-10%), reported by Caputi *et al. Chem Eur J* 14(22):6656–62 (2008) |
| AtUGT71C3 | Group E | *Arabidopsis thaliana* | AC067971, F10K1.3, 12382-13812 | |
| AtUGT71C4 | Group E | *Arabidopsis thaliana* | AC067971, F10K1.4, 14158-15597 | |
| AtUGT71C5 | Group E | *Arabidopsis thaliana* | AC067971, F10K1, 16229-17671 | |
| AtUGT71D1 | Group E | *Arabidopsis thaliana* | AC005496, T27A16, 45718-47121 | |
| AtUGT71D2 | Group E | *Arabidopsis thaliana* | AC005496, T27A16, 40783-42186 | |
| AtUGT72B1 | Group E | *Arabidopsis thaliana* | AL161491, 75572-77014 | |
| AtUGT72B2 | Group E | *Arabidopsis thaliana* | AC023628, F6F3.19, 89227-90669 | |
| AtUGT72B3 | Group E | *Arabidopsis thaliana* | AC023628, F6F3.22, 95474-96919 | |
| AtUGT72C1 | Group E | *Arabidopsis thaliana* | AL161590, 17779-19152 | |
| AtUGT72D1 | Group E | *Arabidopsis thaliana* | AC006135, F24H14, 59343-60755 | |
| AtUGT72E1 | Group E | *Arabidopsis thaliana* | AL132979, T3A5, 67179-68615 | |
| AtUGT72E2 | Group E | *Arabidopsis thaliana* | AB018119, MSN2, 30560-32005 | |
| AtUGT72E3 | Group E | *Arabidopsis thaliana* | AF077407, F9D12, 79017-80462 | |
| AtUGT88A1 | Group E | *Arabidopsis thaliana* | AP000373, 56926-58404 | Low activity on geraniol (10-40%), reported by Caputi *et al. Chem Eur J* 14(22):6656–62 (2008) |
| AtUGT78D1 | Group F | *Arabidopsis thaliana* | AC009917, F26G16, 65767-67224 | |
| AtUGT78D2 | Group F | *Arabidopsis thaliana* | AL391141, F2K13, 64922-66486 | |
| AtUGT78D3 | Group F | *Arabidopsis thaliana* | AL391141, F2K13, 60292-61817 | |
| AtUGT85A1 | Group G | *Arabidopsis thaliana* | AC006551, F12K8, 101428-104184 | High activity on geraniol (>40%) reported by Caputi *et al. Chem Eur J* 14(22):6656–62 (2008) |
| AtUGT85A2 | Group G | *Arabidopsis thaliana* | AC068562, 3508-5967 | High activity on geraniol (>40%) reported by Caputi *et al. Chem Eur J* 14(22):6656–62 (2008) |
| AtUGT85A3 | Group G | *Arabidopsis thaliana* | AC006551, F12K8, 105703-107198; AC068562, T16E15, | |
| AtUGT85A4 | Group G | *Arabidopsis thaliana* | AC013430, 74211-75743 | High activity on geraniol (>40%) reported by Caputi *et al. Chem Eur J* 14(22):6656–62 (2008) |
| AtUGT85A5 | Group G | *Arabidopsis thaliana* | AC068562, T16E15.2, 1156-2919 | Very low activity on geraniol (1-10%), reported by Caputi *et al. Chem Eur J* 14(22):6656–62 (2008) |
| AtUGT85A7 | Group G | *Arabidopsis thaliana* | AC068562, T16E15.5, 8945-10571 | High activity on geraniol (>40%) reported by Caputi *et al. Chem Eur J* 14(22):6656–62 (2008) |
| AtUGT76B1 | Group H | *Arabidopsis thaliana* | AC073395, F11B9, 93894-95312 | |
| AtUGT76C1 | Group H | *Arabidopsis thaliana* | AB017060, K18J17 | |

| **Enzyme** | **Group code** | **Organism** | **Genbank ID** | **Notes** |
|---|---|---|---|---|
| AtUGT76C2 | Group H | *Arabidopsis thaliana* | AB005237, MJJ3, 86268-87835; AB017060, K18J17, 1-2 | |
| AtUGT76C3 | Group H | *Arabidopsis thaliana* | AB017060, K18J17, 7401-9269 | |
| AtUGT76C4 | Group H | *Arabidopsis thaliana* | AB017060, K18J17, 2536-4403 | |
| AtUGT76C5 | Group H | *Arabidopsis thaliana* | AB017060, K18J17, 5455-6900 | |
| AtUGT76D1 | Group H | *Arabidopsis thaliana* | AC002505, T9J22, 49628-51237 | Low activity on geraniol (10-40%), reported by Caputi *et al. Chem Eur J* 14(22):6656–62 (2008) |
| AtUGT76E1 | Group H | *Arabidopsis thaliana* | AB025604, F2O15, 69708-71158 | |
| AtUGT76E111 | Group H | *Arabidopsis thaliana* | AL133314, F12A12, 91897-93329 | Very low activity on geraniol (1-10%), reported by Caputi *et al. Chem Eur J* 14(22):6656–62 (2008) |
| AtUGT76E122 | Group H | *Arabidopsis thaliana* | AL133314, F12A12, 88508-89949 | High activity on geraniol (>40%) reported by Caputi *et al. Chem Eur J* 14(22):6656–62 (2008) |
| AtUGT76E2 | Group H | *Arabidopsis thaliana* | AB025604, F2O15, 72621-74054 | High activity on geraniol (>40%) reported by Caputi *et al. Chem Eur J* 14(22):6656–62 (2008) |
| AtUGT76E3 | Group H | *Arabidopsis thaliana* | AL096859, T6H20, 92466-93881 | |
| AtUGT76E4 | Group H | *Arabidopsis thaliana* | AL133314, F12A12, 96862-98299 | |
| AtUGT76E5 | Group H | *Arabidopsis thaliana* | AL096859, T6H20, 81966-83384 | |
| AtUGT76E6 | Group H | *Arabidopsis thaliana* | AL133314, F12A12, 94420-95845 | |
| AtUGT76E7 | Group H | *Arabidopsis thaliana* | AB028606, F16F17, 34184-35615 | |
| AtUGT76E9 | Group H | *Arabidopsis thaliana* | AB028606, F16F17, 7449-9225 | |
| AtUGT76F1 | Group H | *Arabidopsis thaliana* | AL161667, F1I16, 61935-63899 | |
| AtUGT76F2 | Group H | *Arabidopsis thaliana* | AL161667, F1I16, 59290-61366 | |
| AtUGT83A1 | Group I | *Arabidopsis thaliana* | AC011664, F1C9, 38853-40497 | |
| AtUGT87A1 | Group J | *Arabidopsis thaliana* | AC004165, T27E13, 50431-51895 | |
| AtUGT87A2 | Group J | *Arabidopsis thaliana* | AC004165, 47973-49464 | |
| AtUGT86A1 | Group K | *Arabidopsis thaliana* | AC006922, T1J8, 88046-89708 | |
| AtUGT86A2 | Group K | *Arabidopsis thaliana* | AC005851, 28144-30597 | |
| **AtUGT74B1** | **Group L** | *Arabidopsis thaliana* | AC002396, F3I6, 4859-6322 | |
| **AtUGT74C1** | **Group L** | *Arabidopsis thaliana* | AC006533, T9H9, 34299-36197 | |
| **AtUGT74D1** | **Group L** | *Arabidopsis thaliana* | AC006533, T9H9, 13342-15897 | |
| **AtUGT74E1** | **Group L** | *Arabidopsis thaliana* | AC007153, F3F20, 82737-84239 | |
| **AtUGT74E2** | **Group L** | *Arabidopsis thaliana* | AC007153, F3F20, 84720-86163 | |
| **AtUGT74F1** | **Group L** | *Arabidopsis thaliana* | AC002333, F18O19, 82534-84019 | |
| **AtUGT74F2** | **Group L** | *Arabidopsis thaliana* | AC002333, F18O19, 77132-78568 | |
| **AtUGT75B1** | **Group L** | *Arabidopsis thaliana* | AC007153, F3F20, 27198-28607 | |
| **AtUGT75B2** | **Group L** | *Arabidopsis thaliana* | AC007153, F3F20, 18020-19387 | |
| **AtUGT75C1** | **Group L** | *Arabidopsis thaliana* | AL161538, 19874-21244 | |
| **AtUGT75D1** | **Group L** | *Arabidopsis thaliana* | AL161542, 3576-5000 | |
| **AtUGT84A1** | **Group L** | *Arabidopsis thaliana* | AL161541, 167121-168572 | |

| Enzyme | Group code | Organism | Genbank ID | Notes |
|---|---|---|---|---|
| AtUGT84A2 | Group L | *Arabidopsis thaliana* | AB019232, MIL23, 33583-35070 | |
| AtUGT84A3 | Group L | *Arabidopsis thaliana* | AL161541, 170982-172421 | |
| AtUGT84A4 | Group L | *Arabidopsis thaliana* | AL161541, 175213-176640 | |
| AtUGT84B1 | Group L | *Arabidopsis thaliana* | AC002391, F21P24, 58701-60071 | |
| AtUGT84B2 | Group L | *Arabidopsis thaliana* | AC002391, F21P24, 56464-57816 | |
| AtUGT92A1 | Group M | *Arabidopsis thaliana* | AL353013, T24H18, 21475-22941 | |
| AtUGT82A1 | Group N | *Arabidopsis thaliana* | AP002046, MMP21, 6263-8510 | |
| **CsUGT85K11** | **Group G** | *Camellia sinensis (tea)* | BAO51834 | Activity on geraniol reported by Ohgami *et al. Plant Physiology* 168(2):464–77 (2015) |
| **CrUGT709C2 (7-DLGT)** | **Group P** | *Catharanthus roseus (Madagascar periwinkle)* | BAO01109.1 | Iridoid activity reported by Asada *et al. Plant Cell* 25(10):4123–34 (2013) |
| **GjUGT85A24** | **Group G** | *Gardenia jasminoides (Cape jasmine)* | BAK55737.1 | Iridoid activity reported by Nagatoshi *et al. J Biol Chem* 286(37):32866–74 (2011) |
| **NbUGT79A16** | **Group A** | *Nicotiana benthamiana* | MT945391 | |
| **NbUGT91A20** | **Group A** | *Nicotiana benthamiana* | MT945398 | |
| **NbUGT91R10** | **Group A** | *Nicotiana benthamiana* | MT945403 | |
| **NbUGT91S4** | **Group A** | *Nicotiana benthamiana* | MT945363 | |
| **NbUGT94AQ1** | **Group A** | *Nicotiana benthamiana* | MT945339 | |
| **NbUGT94AR1** | **Group A** | *Nicotiana benthamiana* | MT945389 | |
| **NbUGT94E7** | **Group A** | *Nicotiana benthamiana* | MT945325 | |
| **NbUGT94U2** | **Group A** | *Nicotiana benthamiana* | MT945381 | |
| NbUGT89D20 | Group B | *Nicotiana benthamiana* | MT945364 | |
| NbUGT89V1 | Group B | *Nicotiana benthamiana* | MT945356 | |
| NbUGT90A22 | Group C | *Nicotiana benthamiana* | MT945366 | |
| NbUGT90K1 | Group C | *Nicotiana benthamiana* | MT945337 | |
| **NbUGT73A24** | **Group D** | *Nicotiana benthamiana* | MT945326 | High activity on geraniol (81%) reported by Sun *et al. Plant J* 100:20–37 (2019) |
| **NbUGT73A25** | **Group D** | *Nicotiana benthamiana* | MT945327 | High activity on geraniol (47%) reported by Sun *et al. Plant J* 100:20–37 (2019) |
| **NbUGT73A32** | **Group D** | *Nicotiana benthamiana* | MT945371 | |
| **NbUGT73AB10** | **Group D** | *Nicotiana benthamiana* | MT945365 | |
| **NbUGT73BY1** | **Group D** | *Nicotiana benthamiana* | MT945354 | |
| **NbUGT73BZ1** | **Group D** | *Nicotiana benthamiana* | MT945359 | |
| **NbUGT73E20** | **Group D** | *Nicotiana benthamiana* | MT945362 | |
| **NbUGT73E21** | **Group D** | *Nicotiana benthamiana* | MT945336 | |
| **NbUGT73E22** | **Group D** | *Nicotiana benthamiana* | MT945368 | |
| **NbUGT73E23** | **Group D** | *Nicotiana benthamiana* | MT945372 | |
| **NbUGT73Q2** | **Group D** | *Nicotiana benthamiana* | MT945333 | |
| **NbUGT71A56** | **Group E** | *Nicotiana benthamiana* | MT945345 | |
| **NbUGT71A57** | **Group E** | *Nicotiana benthamiana* | MT945369 | |
| **NbUGT71AJ1** | **Group E** | *Nicotiana benthamiana* | MT945347 | Very low activity on geraniol (1%) reported by Sun *et al. Plant J* 100:20–37 (2019) |
| **NbUGT71AT2** | **Group E** | *Nicotiana benthamiana* | MT945342 | |

| Enzyme | Group code | Organism | Genbank ID | Notes |
|---|---|---|---|---|
| **NbUGT71AT3** | **Group E** | *Nicotiana benthamiana* | MT945324 | |
| **NbUGT71AT4** | **Group E** | *Nicotiana benthamiana* | MT945343 | |
| **NbUGT71AT5** | **Group E** | *Nicotiana benthamiana* | MT945386 | |
| **NbUGT71AU1** | **Group E** | *Nicotiana benthamiana* | MT945367 | |
| **NbUGT71AV1** | **Group E** | *Nicotiana benthamiana* | MT945376 | |
| **NbUGT71X3** | **Group E** | *Nicotiana benthamiana* | MT945350 | |
| **NbUGT71X4** | **Group E** | *Nicotiana benthamiana* | MT945358 | |
| **NbUGT72AX1** | **Group E** | *Nicotiana benthamiana* | MT945344 | Very low activity on geraniol (2%) reported by Sun *et al. Plant J* 100:20–37 (2019) |
| **NbUGT72AY1** | **Group E** | *Nicotiana benthamiana* | MT945401 | Low activity on geraniol (23%) reported by Sun *et al. Plant J* 100:20–37 (2019) |
| **NbUGT72B34*G322V** | **Group E** | *Nicotiana benthamiana* | MT945340 | Very low activity on geraniol (11%) reported by Sun *et al. Plant J* 100:20–37 (2019) |
| **NbUGT72B35** | **Group E** | *Nicotiana benthamiana* | MT945341 | Very low activity on geraniol (1%) reported by Sun *et al. Plant J* 100:20–37 (2019) |
| **NbUGT72B58** | **Group E** | *Nicotiana benthamiana* | MT945379 | |
| NbUGT78C2 | Group F | *Nicotiana benthamiana* | MT945402 | |
| **NbUGT85A104** | **Group G** | *Nicotiana benthamiana* | MT945331 | |
| **NbUGT85A73** | **Group G** | *Nicotiana benthamiana* | MT945328 | High activity on geraniol (100%) reported by Sun *et al. Plant J* 100:20–37 (2019) |
| **NbUGT85A74** | **Group G** | *Nicotiana benthamiana* | MT945370 | Reported as inactive by Sun *et al. Plant J* 100:20–37 (2019) |
| NbUGT76A4 | Group H | *Nicotiana benthamiana* | MT945330 | |
| NbUGT76A5 | Group H | *Nicotiana benthamiana* | MT945393 | |
| NbUGT87AB1 | Group J | *Nicotiana benthamiana* | MT945378 | |
| NbUGT86A25 | Group K | *Nicotiana benthamiana* | MT945374 | |
| NbUGT86A26 | Group K | *Nicotiana benthamiana* | MT945380 | |
| NbUGT86A27 | Group K | *Nicotiana benthamiana* | MT945388 | |
| **NbUGT74B13** | **Group L** | *Nicotiana benthamiana* | MT945360 | |
| **NbUGT74N4** | **Group L** | *Nicotiana benthamiana* | MT945323 | |
| **NbUGT74N5** | **Group L** | *Nicotiana benthamiana* | MT945334 | |
| **NbUGT74N6** | **Group L** | *Nicotiana benthamiana* | MT945361 | |
| **NbUGT74P7** | **Group L** | *Nicotiana benthamiana* | MT945332 | |
| **NbUGT74P8** | **Group L** | *Nicotiana benthamiana* | MT945355 | |
| **NbUGT74P9** | **Group L** | *Nicotiana benthamiana* | MT945405 | |
| **NbUGT74T6** | **Group L** | *Nicotiana benthamiana* | MT945322 | |
| **NbUGT75A4** | **Group L** | *Nicotiana benthamiana* | MT945348 | |
| **NbUGT75A5** | **Group L** | *Nicotiana benthamiana* | MT945351 | |
| **NbUGT75S2** | **Group L** | *Nicotiana benthamiana* | MT945353 | |
| **NbUGT84A75** | **Group L** | *Nicotiana benthamiana* | MT945384 | |
| **NbUGT84A76** | **Group L** | *Nicotiana benthamiana* | MT945395 | |
| NbUGT92G9 | Group M | *Nicotiana benthamiana* | MT945373 | |
| NbUGT82E1 | Group N | *Nicotiana benthamiana* | MT945399 | |
| **NbUGT93S1** | **Group O** | *Nicotiana benthamiana* | MT945346 | |
| **NbUGT93S2** | **Group O** | *Nicotiana benthamiana* | MT945394 | |

| Enzyme | Group code | Organism | Genbank ID | Notes |
|---|---|---|---|---|
| **NbUGT93T1** | **Group O** | *Nicotiana benthamiana* | MT945352 | |
| **NbUGT93T2** | **Group O** | *Nicotiana benthamiana* | MT945357 | |
| **NbUGT93T3** | **Group O** | *Nicotiana benthamiana* | MT945382 | |
| **NbUGT93U1** | **Group O** | *Nicotiana benthamiana* | MT945375 | |
| **NbUGT709B3** | **Group P** | *Nicotiana benthamiana* | MT945392 | |
| **NbUGT709J6** | **Group P** | *Nicotiana benthamiana* | MT945329 | |
| **NbUGT709L7** | **Group P** | *Nicotiana benthamiana* | MT945349 | |
| **NbUGT709Q1** | **Group P** | *Nicotiana benthamiana* | MT945377 | Reported as inactive by Sun *et al. Plant J* 100:20–37 (2019) |
| **NbUGT709Q2** | **Group P** | *Nicotiana benthamiana* | MT945387 | |
| **NbUGT709U2** | **Group P** | *Nicotiana benthamiana* | MT945407 | |
| NbUGT95F1 | undefined | *Nicotiana benthamiana* | MT945404 | |
| SbUGT95B1 | undefined | *Sorghum bicolor* | AAF17077.1 | Activity on geraniol reported by Jones *et al. J Biol Chem* 274(50):35483–91 (1999) |
| **VvGT7** | **Group E** | *Vitus vinifera (grape)* | XP_002276546 | Activity on geraniol reported by Bönisch *et al. Plant Physiology* 165(2):561–81 (2014) |
| **VvGT14** | **Group G** | *Vitus vinifera (grape)* | ENA: CCB45585.1 | Activity on geraniol reported by Bönisch *et al. Plant Physiology* 166(1):23–39 (2014) |
| **VvGT16** | **Group G** | *Vitus vinifera (grape)* | ENA: CCB58004.1 | Activity on geraniol reported by Bönisch *et al. Plant Physiology* 166(1):23–39 (2014) |
| VvGT15 | Group H | *Vitus vinifera (grape)* | ENA: CCB43518.1 | Activity on geraniol reported by Bönisch *et al. Plant Physiology* 166(1):23–39 (2014) |

**Supplementary Table S6.** Primers used for amplification of sgRNA scaffolds from pEPOR1CB0022 Addgene#117537 which contains the sgRNA stem extension scaffold sequence first reported by Chen *et al. Cell* 155(7):1479–91 (2013). Details of the U6-promoter and L1 acceptor that the resulting PCR amplicon was assembled with are also provided.

| Construct Name | sgRNA # | target UGT | L1 acceptor | U6 Promoter | F primer | R primer |
|---|---|---|---|---|---|---|
| pEPQDPKN0361 | 2 | NbUGT85104 | Position 2 pICH47742 Addgene#48001 | pICSL90002 (U6-26, A thaliana) Addgene#68261 | tgtGGTCTCtattgTGATATGAAAGCCTCGTATAgtttaagagctatgctggaaac | tGGTCTCtagcgaaaaaaagcaccgact (for all amplifications) |
| | 3 | NbUGT85A73 | Position 3 pICH47751 Addgene#48002 | pEPQD0CM0032 (pUAP-NbU6-1) Addgene#185623 | tgtGGTCTCtttcgACCCGTCTCGTGCATCGTGTgtttaagagctatgctggaaac | |
| | 4 | NbUGT85A74 | Position 4 pICH47761 Addgene#48003 | pEPQD0CM0033 (pUAP-NbU6-2) Addgene#185624 | tgtGGTCTCtctcgAGAGGACCTTATTCTCTTAAgtttaagagctatgctggaaac | |
| | 6 | NbUGT85A73 | Position 6 pICH47781 Addgene#48005 | pICSL90002 (U6-26, A thaliana) Addgene#68261 | tgtGGTCTCtattgCGTGGCCCTGATTCTCTCAAgtttaagagctatgctggaaac | |
| | 7 | NbUGT85104 | Position 7 pICH47791 Addgene#48006 | pEPQD0CM0032 (pUAP-NbU6-1) Addgene#185623 | tgtGGTCTCtttcgTGTGTCGCGTCATCGTTAGAgtttaagagctatgctggaaac | |
| | 1 | NbUGT85A74 | Position 1 pICH47732 Addgene#48000 | pICSL90002 (U6-26, A thaliana) Addgene#68261 | tgtGGTCTCtattGAGAAATGGGTTCTGTTGAAgtttaagagctatgctggaaac | |
| pEPQDPKN0720 | 2 | NbUGT73Q2 | Position 2 pICH47742 Addgene#48001 | pICSL90002 (U6-26, A thaliana) Addgene#68261 | tgtGGTCTCtattGTTGACGTTGCAGCCAAGCTgtttaagagctatgctggaaac | |
| | 3 | NbUGT73A24 and NbUGT73A25 and *(NbUGT73A32 - 2 bp mismatch)* | Position 3 pICH47751 Addgene#48002 | pEPQD0CM0032 (pUAP-NbU6-1) Addgene#185623 | tgtGGTCTCtttcGTGCCATGGAAAACTATTCTgtttaagagctatgctggaaac | |
| | 4 | NbUGT73A24 and NbUGT73A32 *(NbUGT73A25 - 2 bp mismatch)* | Position 4 pICH47761 Addgene#48003 | pEPQD0CM0033 (pUAP-NbU6-2) Addgene#185624 | tgtGGTCTCtctcGCTGCAGAATCAGTAGTCCAgtttaagagctatgctggaaac | |
| | 6 | NbUGT73A24 and NbUGT73A25 and *(NbUGT73A32 - 1 bp mismatch)* | Position 6 pICH47781 Addgene#48005 | pICSL90002 (U6-26, A thaliana) Addgene#68261 | tgtGGTCTCtattgTTAATTTCGTGAGGCAAATTgtttaagagctatgctggaaac | |
| | 7 | NbUGT73Q2 | Position 7 pICH47791 Addgene#48006 | pEPQD0CM0032 (pUAP-NbU6-1) Addgene#185623 | tgtGGTCTCtttcGAGAATTTAGGTGCGTTGAGgtttaagagctatgctggaaac | |
| | 1 | NbUGT73A24 and NbUGT73A25 | Position 1 pICH47732 Addgene#48000 | pICSL90002 (U6-26, A thaliana) Addgene#68261 | tgtGGTCTCtattgAGCTTCGCCATGTCTAGTGTgtttaagagctatgctggaaac | |
| pEPQDPKN0724 | 2 | NbUGT72B34 and NbUGT72B35 and *(NbUGT72B58 - 1 bp mismatch)* | Position 2 pICH47742 Addgene#48001 | pICSL90002 (U6-26, A thaliana) Addgene#68261 | tgtGGTCTCtattgTGAGCTTGTGGGGCCCAATTgtttaagagctatgctggaaac | |
| | 3 | NbUGT72AY1 | Position 3 pICH47751 Addgene#48002 | pEPQD0CM0032 (pUAP-NbU6-1) Addgene#185623 | tgtGGTCTCtttcGAAAACCACTCTCACCAATGgtttaagagctatgctggaaac | |
| | 4 | NbUGT72AX1 | Position 4 pICH47761 Addgene#48003 | pEPQD0CM0033 (pUAP-NbU6-2) Addgene#185624 | tgtGGTCTCtctcGGATGATGTGTCCCATGCCAgtttaagagctatgctggaaac | |
| | 6 | NbUGT72AY1 | Position 6 pICH47781 Addgene#48005 | pICSL90002 (U6-26, A thaliana) Addgene#68261 | tgtGGTCTCtattGGTTGCCTAAGACTAGAACTgtttaagagctatgctggaaac | |
| | 7 | NbUGT72B34 and NbUGT72B35 | Position 7 pICH47791 Addgene#48006 | pEPQD0CM0032 (pUAP-NbU6-1) Addgene#185623 | tgtGGTCTCtttcGTTTCAATTTTAACATCAAGgtttaagagctatgctggaaac | |
| | 1 | NbUGT72B58 | Position 1 pICH47732 Addgene#48000 | pICSL90002 (U6-26, A thaliana) Addgene#68261 | tgtGGTCTCtattgTTAGAGATGGGACCGTTAGTgtttaagagctatgctggaaac | |

**Supplementary Table S7.** Primers used for construction of mobile single guide RNA plasmid vectors. pEPQDKN0761 (Addgene #185630) containing an sgRNA fused to truncated flowering locus T was used as a PCR template.

| Construct | Target UGT | Position in final construct | F Primer | R Primer |
|---|---|---|---|---|
| pEPQDKN0777 | NbUGT71AT3 | 1 | aCGTCTCgcaggcacctgcaacgAAACTTTTGAACACCATAGTCTAAgttttagagctag | tCGTCTCccgagtcacctgctagtGCACgctcaacacgtacccggccgcgattggccataagtaacctttagagt |
| | NbUGT74P7 | 2 | aCGTCTCgcaggcacctgcaacgGTGCGATTTATGTGGCCTTGTGCGgttttagagctag | tCGTCTCccgagtcacctgctagtCCTGgctcaacacgtacccggccgcgattggccataagtaacctttagagt |
| | NbUGT90K1 | 3 | aCGTCTCgcaggcacctgcaacgCAGGGGTGGCGCAAAAGGGCCGCGgttttagagctag | tCGTCTCccgagcacctgctagtCACTttggccataagtaaccttt |
| pEPQDKN0778 | NbUGT71A56 | 1 | aCGTCTCgcaggcacctgcaacgAAACTCACCTTGTACCCACTGTGGgttttagagctag | tCGTCTCccgagtcacctgctagtGCACgctcaacacgtacccggccgcgattggccataagtaacctttagagt |
| | NbUGT73E22 | 2 | aCGTCTCgcaggcacctgcaacgGTGCATCGCGCAGTGTCAATCATAgttttagagctag | tCGTCTCccgagtcacctgctagtCCTGgctcaacacgtacccggccgcgattggccataagtaacctttagagt |
| | NbUGT93T3 | 3 | aCGTCTCgcaggcacctgcaacgCAGGGTAAACAGGAAGACCATAAGgttttagagctag | tCGTCTCccgagcacctgctagtCACTttggccataagtaaccttt |
| pEPQDKN0779 | NbUGT73BY1 | 1 | aCGTCTCgcaggcacctgcaacgAAACCCACACCTCTCAATGCCCAAgttttagagctag | tCGTCTCccgagtcacctgctagtGCACgctcaacacgtacccggccgcgattggccataagtaacctttagagt |
| | NbUGT74N6 | 2 | aCGTCTCgcaggcacctgcaacgGTGCCTACCAAGCGGAAAGCTTCGgttttagagctag | tCGTCTCccgagtcacctgctagtCCTGgctcaacacgtacccggccgcgattggccataagtaacctttagagt |
| | NbUGT71A57 | 3 | aCGTCTCgcaggcacctgcaacgCAGGCTTATGACCGGCAATAAACGgttttagagctag | tCGTCTCccgagcacctgctagtCACTttggccataagtaaccttt |
| pEPQDKN0780 | NbUGT74T6 | 1 | aCGTCTCgcaggcacctgcaacgAAACGTATCGATACCATTTCCGATgttttagagctag | tCGTCTCccgagtcacctgctagtGCACgctcaacacgtacccggccgcgattggccataagtaacctttagagt |
| | NbUGT74N4 | 2 | aCGTCTCgcaggcacctgcaacgGTGCCATAAACTATGCAATTCACAgttttagagctag | tCGTCTCccgagtcacctgctagtCCTGgctcaacacgtacccggccgcgattggccataagtaacctttagagt |
| | NbUGT94E7 | 3 | aCGTCTCgcaggcacctgcaacgCAGGCTAAGGGCGTGATTGAGATGgttttagagctag | tCGTCTCccgagcacctgctagtCACTttggccataagtaaccttt |
| pEPQDKN0781 | NbUGT85A73 | 1 | aCGTCTCgcaggcacctgcaacgAAACCGTGGCCCTGATTCTCTCAAgttttagagctag | tCGTCTCccgagtcacctgctagtGCACgctcaacacgtacccggccgcgattggccataagtaacctttagagt |
| | NbUGT73A24 and NbUGT73A25 | 2 | aCGTCTCgcaggcacctgcaacgGTGCAGCTTCGCCATGTCTAGTGTgttttagagctag | tCGTCTCccgagtcacctgctagtCCTGgctcaacacgtacccggccgcgattggccataagtaacctttagagt |
| | NbUGT72AY1 | 3 | aCGTCTCgcaggcacctgcaacgCAGGGGTTGCCTAAGACTAGAACTgttttagagctag | tCGTCTCccgagcacctgctagtCACTttggccataagtaaccttt |

**Supplementary Table S8.** Primers used for genotyping plants with Cas9-induced mutations. (A) Primers used for PCR amplification and sequencing of UGT target genes (B) Primers used for ddPCR amplification to determine T-DNA copy number. Asterisk indicates phosphorothioate bond.

**A**

| UGT | Group | Sequencing primer (forward) | Sequencing primer (reverse) | Sequencing primer (alternate rev.) |
|---|---|---|---|---|
| NbUGT85A73 | G | caactcaaatcaactgtgaattttcc | gggagcagcctatatatgtcgg | |
| NbUGT85A74 | G | aattgaaaatattgtctatcgaaagagg | aatacccatttgttatatcacttgtatctg | |
| NbUGT85A104 | G | ccaaaactgcctttgaacaatc | cgtacctttcactggacaataacc | |
| NbUGT73A24 | D | tctattttctgccactaaagcagg | tccactgaatcgaaccaaca*c | ctcttccccagatcgctc*g |
| NbUGT73A25 | D | acacagctttacttttctttctgctac | gttctcaaaacctcagtcactaactt*c | tctcttccccagattgctc*a |
| NbUGT73Q2 | D | ctactgcagttttttctttctcatcag | gaactcctatctttaaaatctctacgagc | |
| NbUGT73A32 | D | aataagaactgaaactacatctagcagtagc | atataagtccatatatcttacatgccattg | tgactgactcttcatccgatag*g |
| NbUGT72B34*G322V | E | cctccctcctaatcttattgctat*c | cagaagtagacacatacagtttggc | gaatctgaatcgggtcaggtaag |
| NbUGT72B35 | E | cgcaaataaatcagcgtaattttactag | gcacaattactaccagcttagaacaac | tgaaccgggtcggttacc |
| NbUGT72AX1 | E | taattaaacaaatcgcacaacagtatc | atccaggagtttgaataggttc*g | |
| NbUGT72B58 (first exon) | E | aagaaaagtcttgtttagcaagcag | ggtccagaagttcctttgctt*c | |
| NbUGT72B58 (second exon) | E | ccctctcaaatacacttggatcag | acaagaacaaaacccatgtgctag | |
| NbUGT72AY1 | E | gtagtttgtgaaattgtttatggcttc | cgaattgtaacattacctcgtcattc | |
| NbUGT71AT3 | E | gatgaatgaactaattttcattcctttag | ggaacaatctcagaatccattaaagc | |
| NbUGT74P7 | L | ttagtacataaaagatgacctatggtgc | aaggcaacgaaacaggaacag | |
| NbUGT90K1 | C | actgctatatacataagaccgcgttc | gaaaacgtaaaaggcacgtctg | |
| NbUGT71A56 | E | caaaacatccactgcaacatagtc | cataacttgggattccaaactcg | |
| NbUGT73E22 | D | gtatcaaacaatggctgttgtca*g | gaatgttaaatttttgggcaaca*c | |
| NbUGT93T3 | O | cttctatgggtagctgtgagaatttac | ttaagcaattcttcctcaagttg*g | |
| NbUGT73BY1 | D | caaatccctttaataaaaatccatgc | tgaactcaatcttgtgtggca*c | |
| NbUGT74N6 | L | ctcctcattcataattggccatag | aattgattcaccaacatttcaagt*g | |
| NbUGT71A57 | E | gcaaagaagaaaatggaagatatcaag | acgacttcctatcaagaaatgtgtg | |
| NbUGT74T6 | L | attttagctcttccttatccaagcc | tttaacaccaactcaaaatacgctg | |
| NbUGT74N4 | L | gcttgatcttgccatatccagt*g | ggtacatctgaactctcaattgtacttg | |
| NbUGT94E7 | A | catggatacacaagtaatagaatgtggt | tctttttctttagctgattttgcc | |

**B**

| gene | ddPCR primer (forward) | ddPCR primer (reverse) |
|---|---|---|
| *nptII* | cctgccgagaaagtatccat | tcttcgtccagatcatcctg |
| *N. benthamiana Rdr1* (reference gene) | gttacgccatccatgtgttg | cagagttcaatttgccagca |

### Supplementary Table S9.

| Peak number | RT (min) | m/z | Putative annotation | Ion Formula | M-H | M-H2O-H | M+HCOOH-H | P19 + GPPS + GES | P19 + DXS + GPPS + GES | P19 + DXS + GPPS + GES + G8H | P19 + DXS + GPPS + GES + G8H + GOR | P19 + DXS + GPPS + GES + G8H + GOR + ISY | P19 + DXS + GPPS + GES + G8H + GOR + ISY + MLPL | m/z |
|---|---|---|---|---|---|---|---|---|---|---|---|---|---|---|
| - | 3.18 | 567.1930 | Dihexosyl hydroxycarboxy geranial (+HCOOH-H) | C23H35O16 | C23H36O16 | C23H38O17 | C22H34O14 | | | | y | | y | 567.1930 |
| - | 3.38 | 569.2085 | Dihexosyl hydroxycarboxy geraniol (+HCOOH-H) | C23H37O16 | C23H38O16 | C23H40O17 | C22H36O14 | | | | y | y | y | 569.2085 |
| - | 3.40 | 391.1610 | Hexosyl carboxygeranial (+HCOOH-H) | C17H27O10 | C17H28O10 | C17H30O11 | C16H26O8 | | | | | | y | 391.1610 |
| - | 3.60 | 359.1348 | Pentosyl dihydroxy geranial (+HCOOH-H) or hexosyl carboxygeranic acid | C16H23O9 | C16H24O9 | C16H26O10 | C15H22O7 | y | y | y | y | y | | 359.1348 |
| - | 3.69 | 391.1609 | Hexosyl dihydroxy geranial (+HCOOH-H) | C17H27O10 | C17H28O10 | C17H30O11 | C16H26O8 | y | y | y | y | y | y | 391.1609 |
| - | 3.76 | 359.1348 | Pentosyl dihydroxy geranial (+HCOOH-H) | C16H23O9 | C16H24O9 | C16H26O10 | C15H22O7 | y | y | y | y | | | 359.1348 |
| 1 | 3.84 | 377.1817 | Hexosyl hydroxygeraniol (+HCOOH-H) | C17H29O9 | C17H30O9 | C17H32O10 | C16H28O7 | y | y | y | y | | | 377.1817 |
| - | 3.97 | 393.1765 | Hexosyl dihydroxygeraniol (+HCOOH-H) | C17H29O10 | C17H30O10 | C17H32O11 | C16H28O8 | y | y | y | y | | | 393.1765 |
| - | 3.90 | 361.1503 | Pentosyl hydroxygeranic acid (+HCOOH-H) | C16H25O9 | C16H26O9 | C16H28O10 | C15H24O7 | y | y | y | y | y | y | 361.1503 |
| - | 3.97 | 393.1765 | Hexosyl carboxygeraniol (+HCOOH-H) | C17H29O10 | C17H30O10 | C17H32O11 | C16H28O8 | | y | y | y | y | y | 393.1765 |
| - | 4.00 | 347.1712 | Pentosyl hydroxygeraniol (+HCOOH-H) | C16H27O8 | C16H28O8 | C16H30O9 | C15H26O6 | | | y | | | | 347.1712 |
| - | 4.03 | 595.2235 | Malonyl hexosyl pentosyl hydroxygeraniol (+HCOOH-H) | C25H39O16 | C25H40O16 | C25H42O17 | C24H38O14 | | | y | | | | 595.2235 |
| 2 | 4.12 | 379.1974 | Hexosyl hydroxycitronellal (+HCOOH-H) | C17H31O9 | C17H32O9 | C17H34O10 | C16H30O7 | | | y | y | | | 379.1974 |
| - | 4.27 | 395.1922 | Hexosyl dihydroxycitronellol (+HCOOH-H) | C17H31O10 | C17H32O10 | C17H34O11 | C16H30O8 | | y | y | y | y | y | 395.1922 |
| 3 | 4.35 | 699.2713 | Trihexosyl geranic acid (+HCOOH-H) | C29H47O19 | C29H48O19 | C29H50O20 | C28H46O17 | | y | y | y | | | 699.2713 |
| - | 5.04 | 537.2187 | Dihexosyl geranic acid (+HCOOH-H) | C23H37O14 | C23H38O14 | C23H40O15 | C22H36O12 | y | y | | | | | 537.2187 |
| 4 | 5.32 | 493.2288 | Pentosyl hexosyl geraniol (+HCOOH-H) | C22H37O12 | C22H38O12 | C22H40O13 | C21H36O10 | y | y | | | | | 493.2288 |
| 5 | 5.54 | 519.2445 | Acetyl dihexosyl geraniol | C24H39O12 | C24H40O12 | C24H42O13 | C23H38O10 | y | y | | | | | 519.2445 |
| - | 6.17 | 803.3701 | Malonyl hexosyl geraniol (dimer) | C38H59O18 | C38H60O18 | C38H62O19 | C37H58O16 | y | y | | | | | 803.3701 |
| - | 6.24 | 831.3286 | Malonyl hexosyl geranic acid (dimer) | C38H55O20 | C38H56O20 | C38H58O21 | C37H54O18 | y | y | | | | | 831.3286 |

**Supplementary Table S10** Expression levels and protein features of *N. benthamiana* UDP-glycosyltransferases (UGTs) selected for mutagenesis. Relative activity on geraniol is reported from Sun et al. 2019 *Plant Journal*. 100:20–37

| UGT | Group | Geraniol activity (relative) | Length (amio acids) | His | Asp | PSPG | GSS | GFP/ Untrans-formed | Low geraniol /GFP | High geraniol /GFP | Nepeta-lactol /GFP | untrans-formed | GFP | Low geraniol | High geraniol | Nepeta-lactol |
|---|---|---|---|---|---|---|---|---|---|---|---|---|---|---|---|---|
| | | | | | | | | fold change | | | | normalised counts | | | | |
| NbUGT90K1 | Group C | | 468 | yes | yes | yes | yes | 5.180 | 0.109 | -0.378 | -0.202 | 234 | 9698 | 8983 | 6890 | 7796 |
| NbUGT71AT3 | Group E | | 476 | yes | yes | yes | yes | 8.660 | 0.084 | 0.100 | 0.792 | 28 | 15346 | 14462 | 15514 | 25270 |
| NbUGT74P7 | Group L | | 454 | yes | yes | yes | yes | 4.827 | -0.034 | 0.292 | 0.224 | 292 | 8520 | 8724 | 10707 | 10209 |
| NbUGT73E22 | Group D | | 488 | yes | yes | yes | yes | -1.839 | 0.388 | 1.862 | 2.044 | 602 | 202 | 151 | 613 | 702 |
| NbUGT71A56 | Group E | | 399 | yes | yes | yes | no | -1.267 | 0.347 | 1.330 | 2.046 | 2331 | 1069 | 792 | 2458 | 4448 |
| NbUGT93T3 | Group O | | 473 | yes | yes | yes | yes | -1.998 | 0.127 | 1.135 | -0.070 | 1668 | 451 | 413 | 913 | 393 |
| NbUGT73BY1 | Group D | | 493 | yes | yes | yes | yes | 0.444 | 0.081 | 0.430 | 2.494 | 174 | 270 | 252 | 360 | 1912 |
| NbUGT71A57 | Group E | | 484 | yes | yes | yes | yes | -0.485 | -0.292 | 0.185 | 3.285 | 59 | 29 | 38 | 45 | 635 |
| NbUGT74N6 | Group L | | 458 | yes | yes | yes | yes | 2.582 | 0.024 | 1.104 | 2.956 | 11 | 134 | 131 | 335 | 1527 |
| NbUGT94E7 | Group A | | 327 | yes | yes | yes | no | 4.769 | -0.119 | 0.880 | 2.262 | 38 | 1902 | 2090 | 4188 | 12300 |
| NbUGT74N4 | Group L | | 461 | yes | yes | yes | yes | 3.328 | -0.146 | 0.691 | 2.085 | 403 | 4980 | 5578 | 9515 | 27698 |
| NbUGT74T6 | Group L | | 457 | yes | yes | yes | yes | 1.421 | -0.299 | -0.113 | 0.986 | 8910 | 19815 | 24513 | 22619 | 49467 |
| NbUGT85A73 | Group G | 100 | 486 | yes | yes | yes | yes | -1.293 | 0.012 | 0.149 | 0.605 | 7224 | 2934 | 2910 | 3231 | 4453 |
| NbUGT85A104 | Group G | | 496 | yes | yes | yes | yes | -3.836 | 1.865 | 0.271 | 0.661 | 341 | 83 | 21 | 26 | 35 |
| NbUGT85A74 | Group G | | 462 | yes | yes | yes | no | -1.946 | 0.398 | 0.747 | 0.616 | 1525 | 514 | 388 | 657 | 599 |
| NbUGT73A24 | Group D | 81 | 477 | yes | yes | yes | yes | 0.671 | 0.053 | 1.120 | 1.217 | 3752 | 6326 | 6087 | 13633 | 14616 |
| NbUGT73A25 | Group D | 47 | 477 | yes | yes | yes | yes | 2.688 | 0.114 | 1.114 | 1.614 | 239 | 1929 | 1773 | 4038 | 5836 |
| NbUGT73A32 | Group D | | 412 | yes | yes | yes | no | 1.309 | -0.167 | 0.117 | -0.181 | 271 | 602 | 677 | 735 | 596 |
| NbUGT73Q2 | Group D | | 486 | yes | no | yes | yes | 5.389 | 0.037 | 0.138 | 1.671 | 55 | 3321 | 3231 | 3578 | 11082 |
| NbUGT72AX1 | Group E | 2 | 489 | yes | yes | yes | yes | -0.502 | -0.177 | -0.427 | -0.740 | 11298 | 6959 | 7898 | 5824 | 4658 |
| NbUGT72AY1 | Group E | 23 | 478 | yes | yes | yes | yes | -0.549 | -0.647 | -0.379 | -0.768 | 226 | 92 | 150 | 113 | 84 |
| NbUGT72B34 | Group E | 11 | 479 | yes | yes | yes | yes | 1.979 | -0.097 | -0.070 | 0.356 | 1028 | 3831 | 4099 | 3902 | 5258 |
| NbUGT72B35 | Group E | 1 | 479 | yes | yes | yes | yes | -0.120 | 0.042 | 0.575 | 0.334 | 4025 | 3811 | 3701 | 5530 | 4673 |
| NbUGT72B58 | Group E | | 479 | yes | yes | yes | yes | -1.049 | -0.084 | 0.244 | -0.311 | 1158 | 525 | 557 | 660 | 448 |

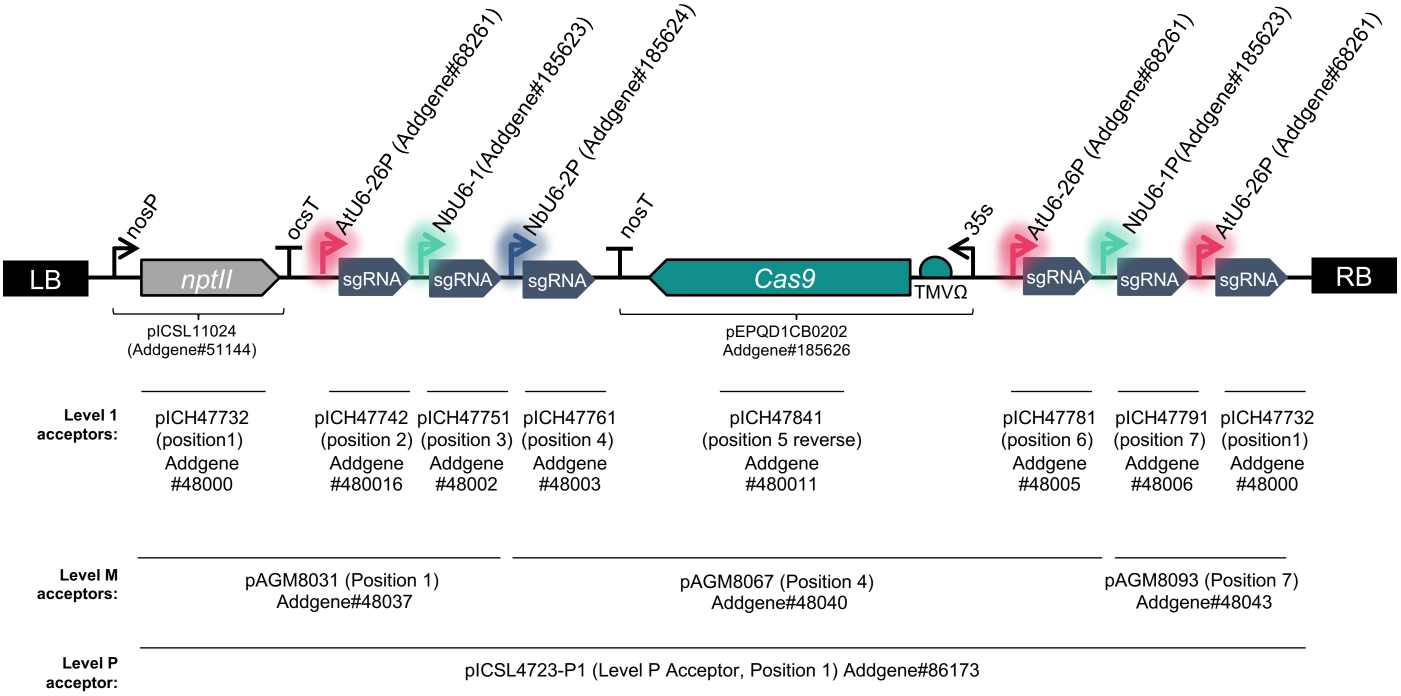


**Supplementary Figure S1*.*** Design and hierarchical assembly of binary constructs used for Cas9-mediated mutagenesis of *Nicotiana benthamiana.*

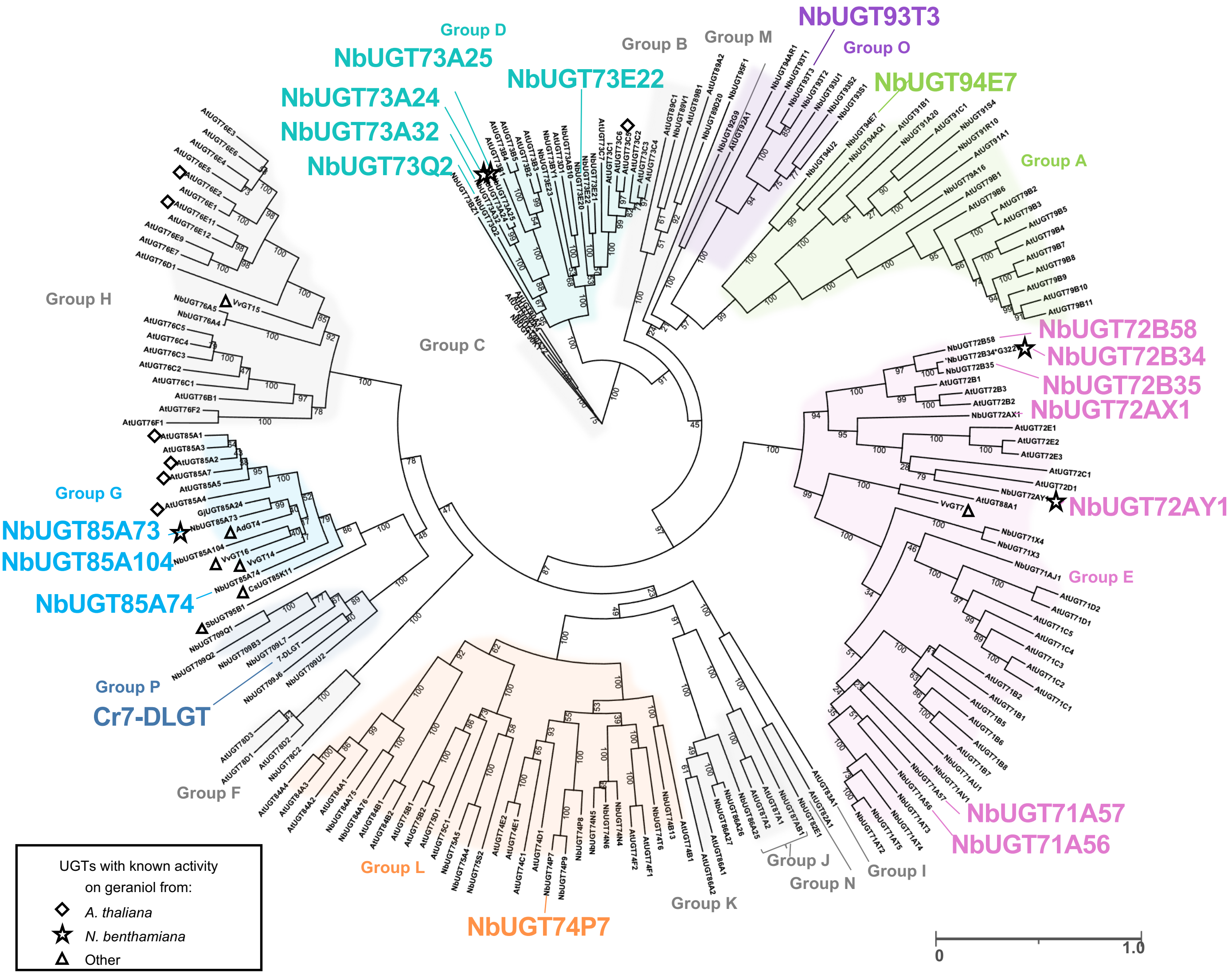


**Supplementary Figure S2. Maximum likelihood (RAxML) phylogenetic comparison of 193 Family 1 UDP-glycosyltransferases (UGTs) from *N. benthamiana* and *A. thaliana* and nine UGT sequences previously shown to be active on geraniol or iridoid substrates.** Groups A-P are annotated according to nomenclature used by Caputi et al 2011. Labeled taxa indicate enzymes in which Cas9-mediated targeted mutations were subsequently introduced. Scale bar indicates the number of substitutions per site.

A

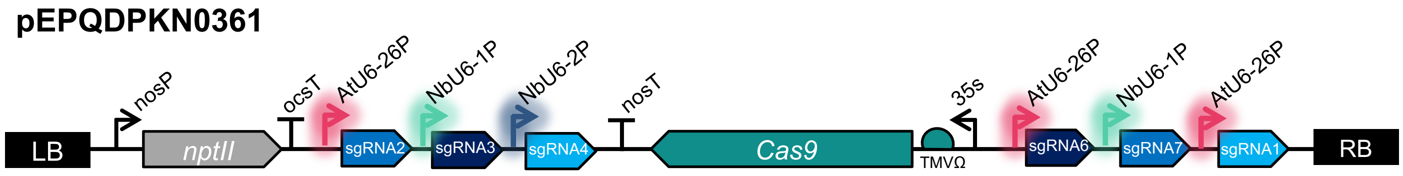


B

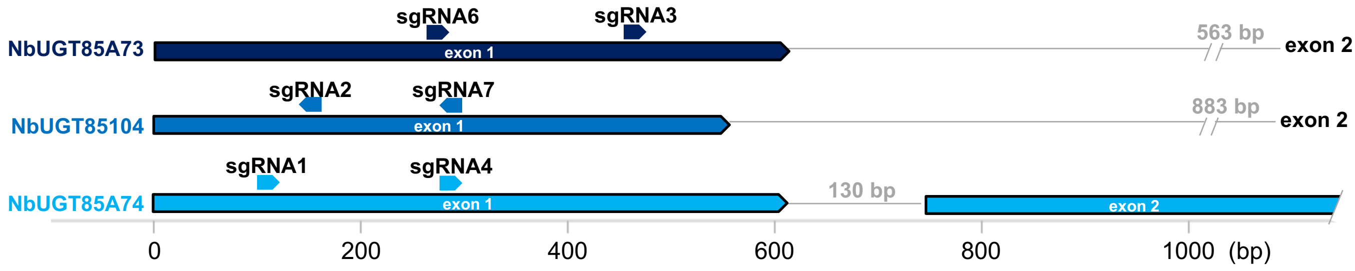


C

| NbUGT85A73 | | | |
|---|---|---|---|
| **Plant line** | **sgRNA6**<br>GAGAAATGGGTTCTGT-TGAAGGG | **sgRNA3**<br>ACCCGTCTCGTGCATCG-TGTCGG | **T-DNA** |
| **0361-34-05** | **bi-allelic**<br>CGTGGCCCTGATTCTCTTCAAGGG<br>CGTGGCCCTGAT-----TCAAGGG<br>frameshift at aa63 and stop codon at aa71 | **homozygous**<br>ACCCGTCTCGTGCATCGTTGTCGG<br>sequence remains out of frame | - |
| **0361-34-08** | **homozygous**<br>CGTGGCCCTGAT-----TCAAGGG<br>frameshift resulting in incorrect sequence from aa62 -125 | **homozygous**<br>ACCCGTCTCGTGCATCGTTGTCGG<br>seqeunce back in frame from S126 | + |

| NbUGT85A104 | | | |
|---|---|---|---|
| **Plant line** | **sgRNA2**<br>CCTCCATAT-ACGAGGCTTTCATA | **sgRNA7**<br>CCCTC-TAACGATGACGCGACACA | **T-DNA** |
| **0361-34-05** | **homozygous**<br>CCTCCATATAACGAGGCTTTCATA<br>frameshift at aa40 and stop codon at aa50 | **homozygous**<br>CCCTCTTAACGATGACGCGACACA<br>sequence remains out of frame | - |
| **0361-34-08** | **homozygous**<br>CCTCCATATAACGAGGCTTTCATA<br>frameshift at aa40 and stop codon at aa50 | **homozygous**<br>CCCTCTTAACGATGACGCGACACA<br>sequence remains out of frame | + |

| NbUGT85A74 | | | |
|---|---|---|---|
| **Plant line** | **sgRNA1**<br>GAGAAATGGGTTCTGTT-GAAGGG | **sgRNA4**<br>AGAGGACCTTATTCTCTTAAAGG | **T-DNA** |
| **0361-34-05** | **bi-allelic**<br>GAGAAATGGGTTCTGTTCGAAGGG<br>GAGAAATGGGTTCTGTTAGAAGGG<br>frameshift at aa5 and stop codon at aa47 | **wild type**<br>AGAGGACCTTATTCTCTTAAAGG<br>n/a | - |
| **0361-34-08** | **homozygous**<br>GAGAAATGGGTTCTGTTAGAAGGG<br>frameshift at aa5 and stop codon at aa47 | **wild type**<br>AGAGGACCTTATTCTCTTAAAGG<br>n/a | + |

**Supplementary Figure S3.** Cas9-mediated mutagenesis of three Group G UGTs (A) Schematic showing construct design (B) locations of sgRNAs (C) sequences of targets and genotypes of T1 plants

A

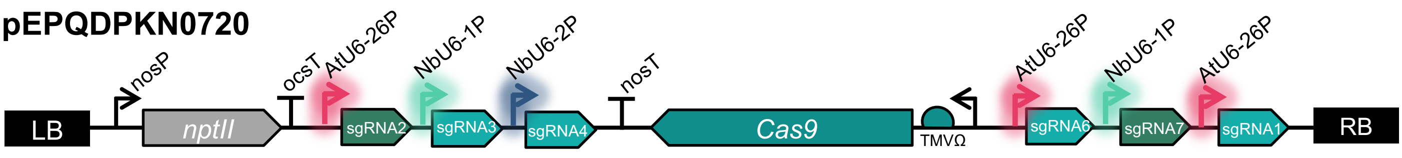


B

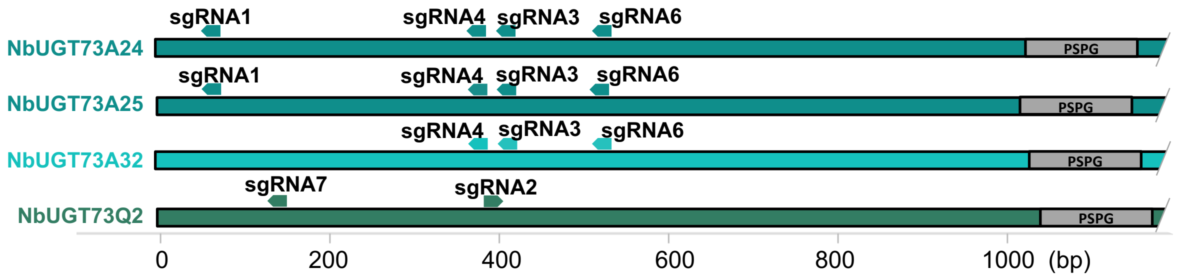


C

**NbUGT73A24**

| Plant line | sgRNA1 | sgRNA4 | sgRNA3 | sgRNA6 | T-DNA |
|---|---|---|---|---|---|
| | CCTACACTAGACATGGCGAAGCT | CCTTGGACTACTGATTCTGCAGC | CCGAGAATAGTTTTCCATGGCAC | CCGAA-TTTGCCTCACGAAATTAA | |
| 0720-06-01 | homozygous<br>CCTACA-----------------<br>319 bp deletion from sgRNA1 to sgRNA4 deleting aa21-126 and introducing frameshift | homozygous<br>-------------ATTCTGCAGC | homozygous<br>CCGAG---AGTTTTCCATGGCAC<br>sequence remains out of frame | homozygous<br>CCGAATTTTGCCTCACGAAATTAA<br>sequence remains out of frame | - |
| 0720-06-03 | homozygous<br>CCTACA-----------------<br>319 bp deletion from sgRNA1 to sgRNA4 deleting aa21-126 and introducing frameshift | homozygous<br>-------------ATTCTGCAGC | homozygous<br>CCGAG---AGTTTTCCATGGCAC<br>sequence remains out of frame | homozygous<br>CCGAATTTTGCCTCACGAAATTAA<br>sequence remains out of frame | + |

**NbUGT73A25**

| Plant line | sgRNA1 | sgRNA4 | sgRNA3 | sgRNA6 | T-DNA |
|---|---|---|---|---|---|
| | CCTACA-CTAGACATGGCGAAGCT | CCTTGGACTACTGATTCTGCAGC<br>Mismatch in sgRNA | CCGAGAATAGTTTTCCATGGCACAAGCTTC | CCGAA-TTTGCCTCACGAAATTAA | |
| 0720-06-01 | bi-allelic<br>CCTACACCTAGACATGGCGAAGCT<br>CCTACA---AGACATGGCGAAGCT<br>frameshift from aa21 and stop codon at aa 32 | wild type<br>CCTTGGACTACTGATAGTGCAGC<br>sequence remains out of frame | bi-allelic<br>CCGAGAA------TCCATGGCACAAGCTTC<br>CCGAGA---------------------TTC<br>sequence remains out of frame | homozygous<br>CCGAATTTTGCCTCACGAAATTAA<br>sequence remains out of frame | - |
| 0720-06-03 | homozygous<br>CCTACACCTAGACATGGCGAAGCT<br>frameshift from aa21 and stop codon at aa 33 | wild type<br>CCTTGGACTACTGATAGTGCAGC<br>sequence remains out of frame | homozygous<br>CCGAGAA------TCCATGGCACAAGCTTC<br>sequence remains out of frame | homozygous<br>CCGAATTTTGCCTCACGAAATTAA<br>sequence remains out of frame | + |

**NbUGT73A32**

| Plant line | sgRNA4 | sgRNA3 | sgRNA6 | T-DNA |
|---|---|---|---|---|
| | CCTTGGACTACTGATTCTGCAGC | CCGAGAATAGTTTTTCATGGTAC<br>Mismatch in sgRNA | CCGAATTTGCCTCACGAAATTAA<br>Mismatch in sgRNA | |
| 0720-06-01 | homozygous<br>CCTTG-------GATTCTGCAGC<br>frameshift from aa125 and stop codon at aa135 | wild type<br>CCAAGAATAGTTTTTCATGGTAC<br>sequence remains out of frame | wild type<br>CCTAATTTGCCTCACGAAATCAA<br>sequence remains out of frame | - |
| 0720-06-03 | homozygous<br>CCTTG-------GATTCTGCAGC<br>frameshift from aa125 and stop codon at aa135 | wild type<br>CCAAGAATAGTTTTTCATGGTAC<br>sequence remains out of frame | wild type<br>CCTAATTTGCCTCACGAAATCAA<br>sequence remains out of frame | + |

**NbUGT73Q2**

| Plant line | sgRNA7 | sgRNA2 | T-DNA |
|---|---|---|---|
| | CCTCTC-AACGCACCTAAATTCTC | GTTGACGTTGCAGCC-AAGCTGGG | |
| 0720-06-01 | homozygous<br>CCTCTCAAACGCACCTAAATTCTC<br>frameshift from aa42 and stop codon at aa49 | homozygous<br>GTTGACGTTGCAGCCAAAGCTGGG<br>sequence remains out of frame | - |
| 0720-06-03 | homozygous<br>CCTCTCAAACGCACCTAAATTCTC<br>frameshift from aa42 and stop codon at aa49 | homozygous<br>GTTGACGTTGCAGCCAAAGCTGGG<br>sequence remains out of frame | + |

**Supplementary Figure S4.** Cas9-mediated mutagenesis of four Group D UGTs (A) Schematic showing construct design (B) locations of sgRNAs (C) sequences of targets and genotypes of T1 plants

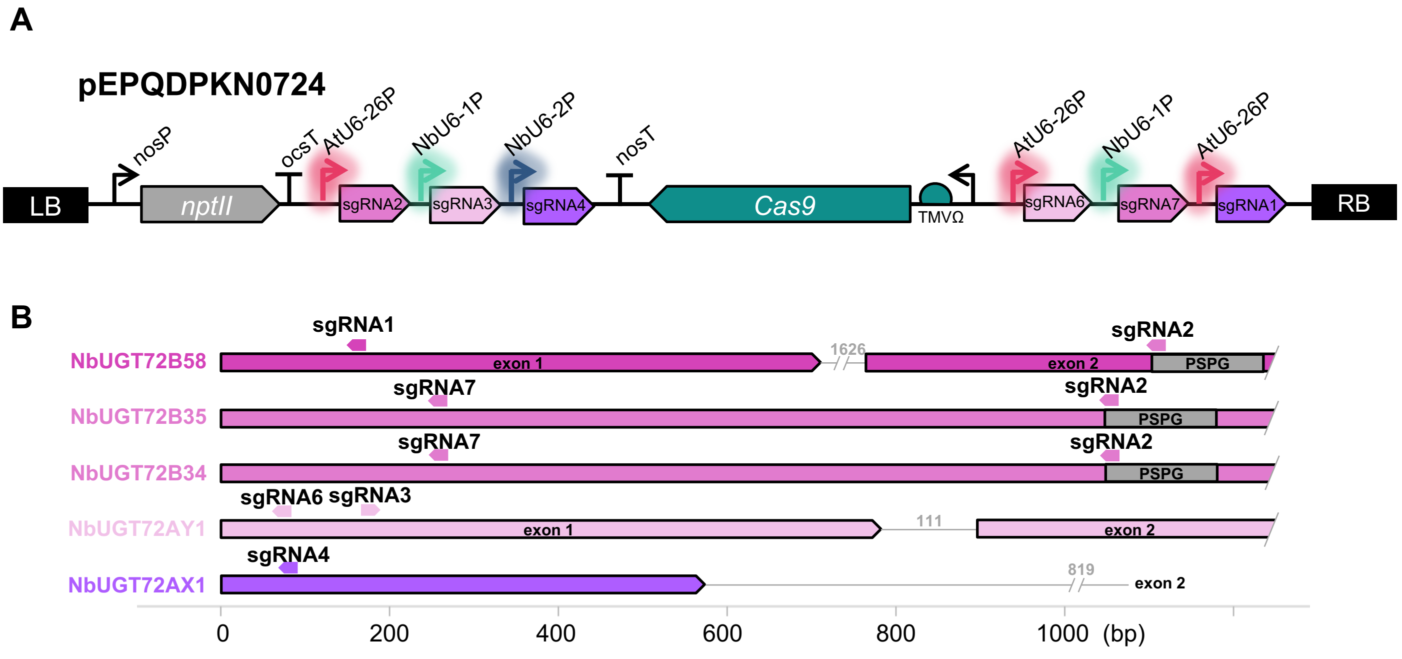


C

| NbUGT72B58 | | | |
|---|---|---|---|
| Plant line | sgRNA1<br>CCTACT-AACGGTCCCATCTCTAA | sgRNA2<br>CCTAATTGGGCCCCACAAGCCCAAATC | T-DNA |
| 0724-37-06 | **homozygous**<br>CCTACTAAACGGTCCCATCTCTAA<br>frameshift at aa52 and stop codon at aa57 | **wild type**<br>CCTAATTGGGCCCCACAAGCCCAAATC<br>n/a | - |
| 0724-22a-06 | **homozygous**<br>CCTACT----GGTCCCATCTCTAA<br>3 bp in-frame deletion of aa52 (not considered a loss-of-function mutation) | **wild type**<br>CCTAATTGGGCCCCACAAGCCCAAATC<br>n/a | + |

| NbUGT72B35 | | | |
|---|---|---|---|
| Plant line | sgRNA7<br>CCTCTTGATGTTAAAATTGAAAC | sgRNA2<br>CCTAATTGGGCCCCACAAGCTCAAATC | T-DNA |
| 0724-37-06 | **wild type**<br>CCTCTTGATGTTAAAATTGAAAC<br>n/a | **wild type**<br>CCTAATTGGGCCCCACAAGCTCAAATT<br>n/a | - |
| 0724-22a-06 | **wild type**<br>CCTCTTGATGTTAAAATTGAAAC<br>n/a | **homozygous**<br>CCTAAT--GGCCCCACAAGCTCAAATT<br>frameshift at aa350 and stop codon at aa357 | + |

| NbUGT72B34 | | | |
|---|---|---|---|
| Plant line | sgRNA7<br>CCTCTTGATGTTAAAATTGAAAC | sgRNA2<br>CCTAA-TTGGGCCCCACAAGCTCAAATT | T-DNA |
| 0724-37-06 | **wild type**<br>CCTCTTGATGTTAAAATTGAAAC<br>n/a | **bi-allelic**<br>CCTAATTTGGGCCCCACAAGCTCAAATT<br>CCTAA--TGGGCCCCACAAGCTCAAATT<br>framshift from aa350 and stop codon aa396 | - |
| 0724-22a-06 | **homozygous**<br>CCTCTTG----------------<br>812 bp deletion and frameshift introducing a stop codon | **homozygous**<br>-------------------------ATT | + |

| NbUGT72AY1 | | | |
|---|---|---|---|
| Plant line | sgRNA6<br>CCCAGT-TCTAGTCTTAGGCAACC | sgRNA3<br>GAAAACCACTCTCACCAATGAGG | T-DNA |
| 0724-37-06 | **homozygous**<br>CCCAGTCTCTAGTCTTAGGCAACC<br>frameshift from aa23, stop codon at aa 34 | **homozygous**<br>GAAAACCACTCTCACC-ATGAGG<br>sequence back in frame, but after stop codon | - |
| 0724-22a-06 | **homozygous**<br>CCCAGTTTCTAGTCTTAGGCAACC<br>frameshift from aa23, stop codon at aa 34 | **homozygous**<br>GAAAACCACTCTCACC-ATGAGG<br>sequence back in frame, but after stop codon | + |

| NbUGT72AX1 | | |
|---|---|---|
| Plant line | sgRNA4<br>CCCTGG-CATGGGACACATCATCC | T-DNA |
| 0724-37-06 | **homozygous**<br>CCCTGG--ATGGGACACATCATCC<br>frameshift from aa26, stop codon at aa 40 | - |
| 0724-22a-06 | **homozygous**<br>CCCTGGACATGGGACACATCATCC<br>frameshift from aa26, stop codon at aa 34 | + |

**Supplementary Figure S5.** Cas9-mediated mutagenesis of five Group E UGTs (A) Schematic showing construct design (B) locations of sgRNAs (C) sequences of targets and genotypes of T1 plants

**A**

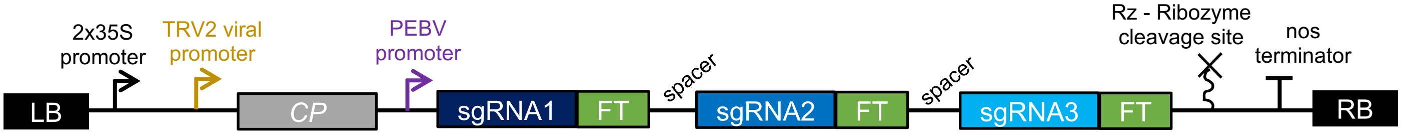


**B**

### pEPQDKN0777

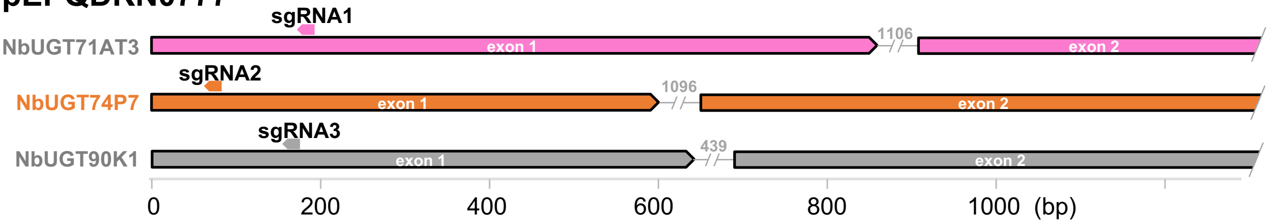


| | NbUGT71AT3 | NbUGT74P7 | NbUGT90K1 |
|---|---|---|---|
| | **sgRNA1** | **sgRNA2** | **sgRNA3** |
| | CCCTTAGACTATGGTGTTCAAAA | CCCCGCACAAGGCCACATAAATC | CCTCGCGGCCCTTTTGCGCCACC |
| **NbCas9-0777B-01** | **wild type**<br>CCCTTAGACTATGGTGTTCAAAA<br>n/a | **homozygous**<br>CCCCGC-----GCCACATAAATC<br>frameshift from aa18, stop codon at aa35 | **wild type**<br>CCTCGCGGCCCTTTTGCGCCACC<br>n/a |

### pEPQDKN0778

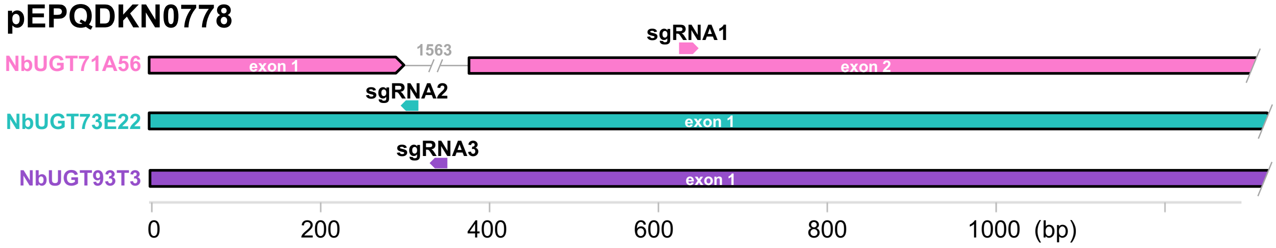


| | NbUGT71A56 | NbUGT73E22 | NbUGT93T3 |
|---|---|---|---|
| | **sgRNA1** | **sgRNA2** | **sgRNA3** |
| | TCACCTTGTACCCACT-GTGGAGG | CCCTA-TGATTGACACTGCGCGAT | CCTCTT-ATGGTCTTCCTGTTTAC |
| **NbCas9-0778B-05** | **homozygous**<br>TCACCTTGTACCCACTGGTGGAGG<br>frameshift from aa21, stop codon at aa54 | **homozygous**<br>CCCTATTGATTGACACTGCGCGAT<br>frameshift from aa24, stop codon at aa26 | **homozygous**<br>CCTCTTAATGGTCTTCCTGTTTAC<br>frameshift from aa47, stop codon at aa47 |

### pEPQDKN0779

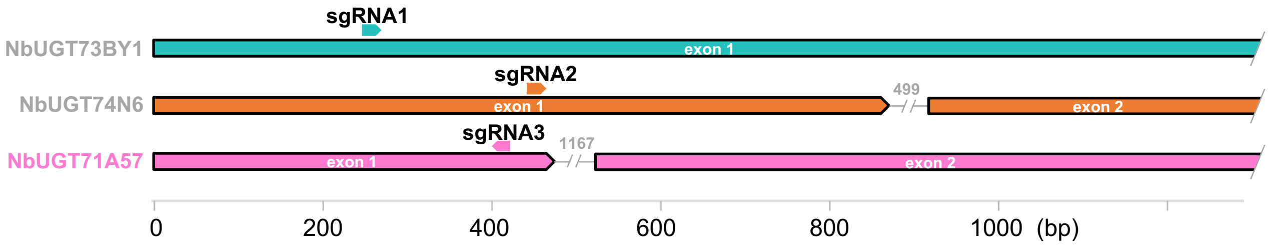


| | NbUGT73BY1 | NbUGT74N6 | NbUGT71A57 |
|---|---|---|---|
| | **sgRNA1** | **sgRNA2** | **sgRNA3** |
| | CCACACCTCTCAATGCCCAAAGG | CTACCAAGCGGAAAGCTTCGCGG | CCGCGTTTATTGCCGGTCATAAG |
| **NbCas9-0779A-01** | **wild type**<br>CCACACCTCTCAATGCCCAAAGG<br>n/a | **wild type**<br>CTACCAAGCGGAAAGCTTCGCGG<br>n/a | **homozygous**<br>CCGCG-TTATTGCCGGTCATAAG<br>frameshift from aa89, stop codon at aa113 |

**Supplementary Figure S6.** Cas9-mediated mutagenesis of 16 UGTs using mobile sgRNAs (A) Schematic showing construct design (B) locations of mobile sgRNAs on target genes and genotypes of target genes in E1 plants

### pEPQDKN0780

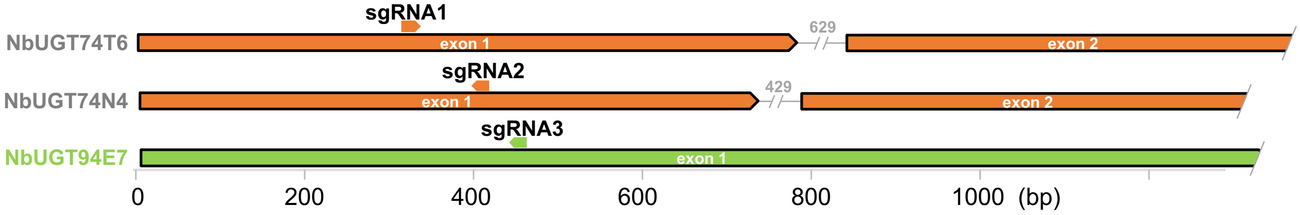


| | NbUGT74T6 | NbUGT74N4 | NbUGT94E7 |
|---|---|---|---|
| | sgRNA1 | sgRNA2 | sgRNA3 |
| | GTATCGATACCATTTCCGATGGG | CCCTGTGAATTGCATAGTTTATG | CCCCAT-CTCAATCACGCCCTTAG |
| **NbCas9-0780B-01** | **wild type**<br>GTATCGATACCATTTCCGATGGG<br>n/a | **wild type**<br>CCCTGTGAATTGCATAGTTTATG<br>n/a | **homozygous**<br>CCCCATCCTCAATCACGCCCTTAG<br>frameshift from aa95, stop codon at aa100 |

### pEPQDKN0781

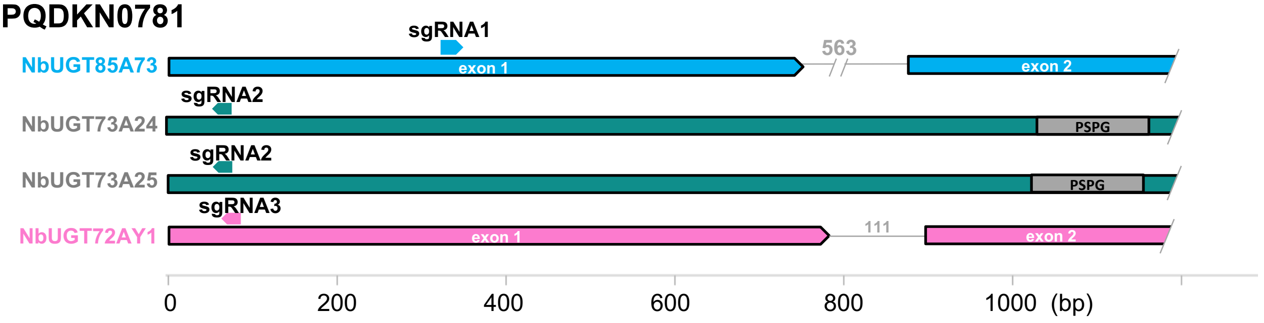


| | NbUGT85A73 | NbUGT73A24 and NbUGT73A25 | NbUGT72AY1 |
|---|---|---|---|
| | sgRNA1 | sgRNA2 | sgRNA3 |
| | CGTGGCCCTGATTCTCTCAAGGG | CCTACACTAGACATGGCGAAGCT | CCCAGTTCTAGTCTTAGGCAACC |
| **NbCas9-0781B-03** | **bi-allelic**<br>CGTGGCCCTGATTCTCT-AAGGG<br>CGTGGCCCTGAT----TCAAGGG<br>frameshift at aa62/63 and stop codon at aa129 | **wild type**<br>CCTACACTAGACATGGCGAAGCT<br>n/a | **homozygous**<br>CCCAG-TCTAGTCTTAGGCAACC<br>frameshift from aa23, stop codon at aa23 |

**Supplementary Figure S6 continued.** Cas9-mediated mutagenesis of 16 UGTs using mobile sgRNAs (A) Schematic showing construct design (B) locations of mobile sgRNAs on target genes and genotypes of target genes in E1 plants

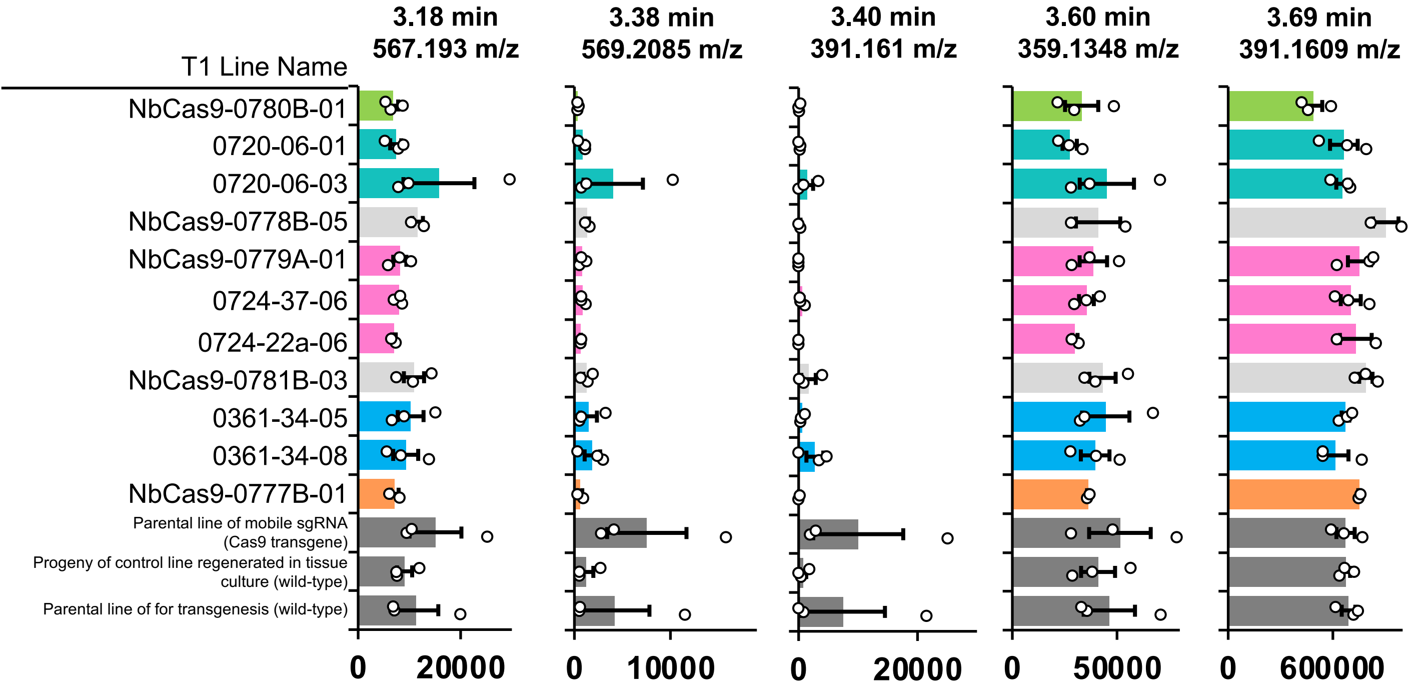


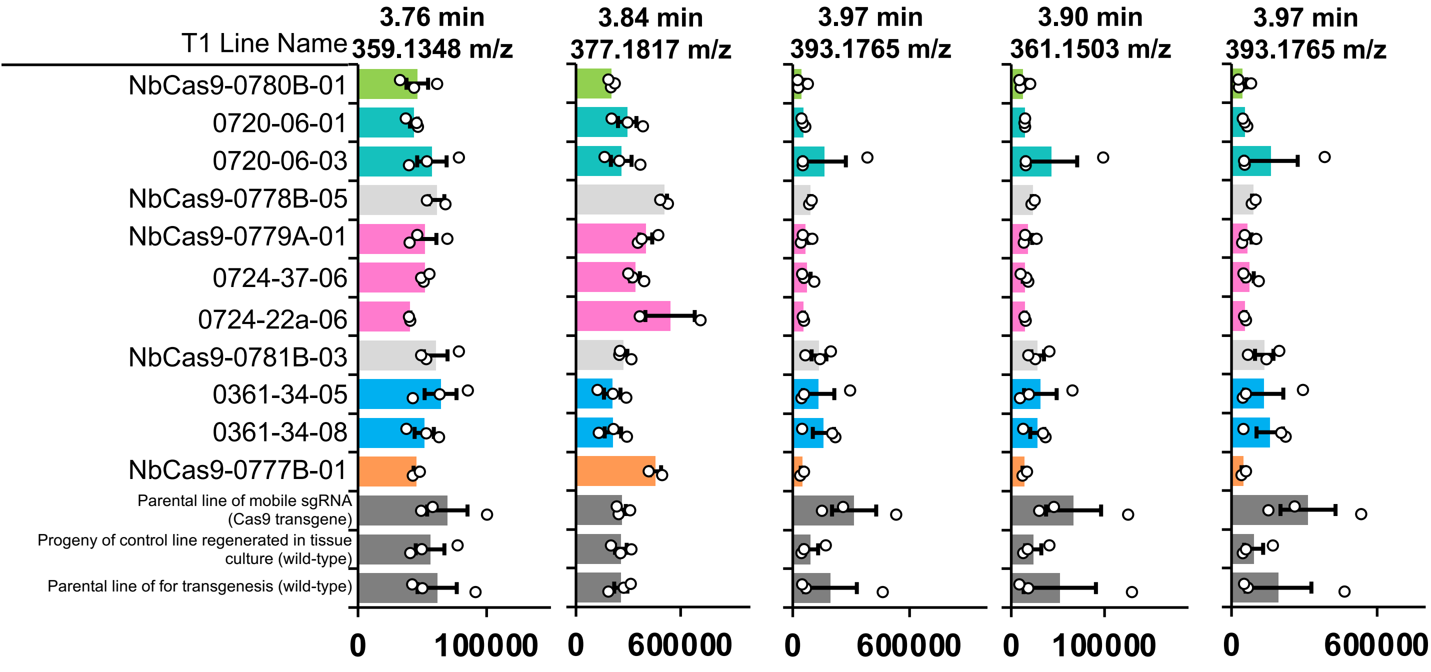


**Supplementary Figure S7.** Abundance of derivative peaks in lines of *N. benthamiana* infiltrated with the geraniol biosynthesis pathway (P19 + **DXS** + **GPPS** + **GES**)

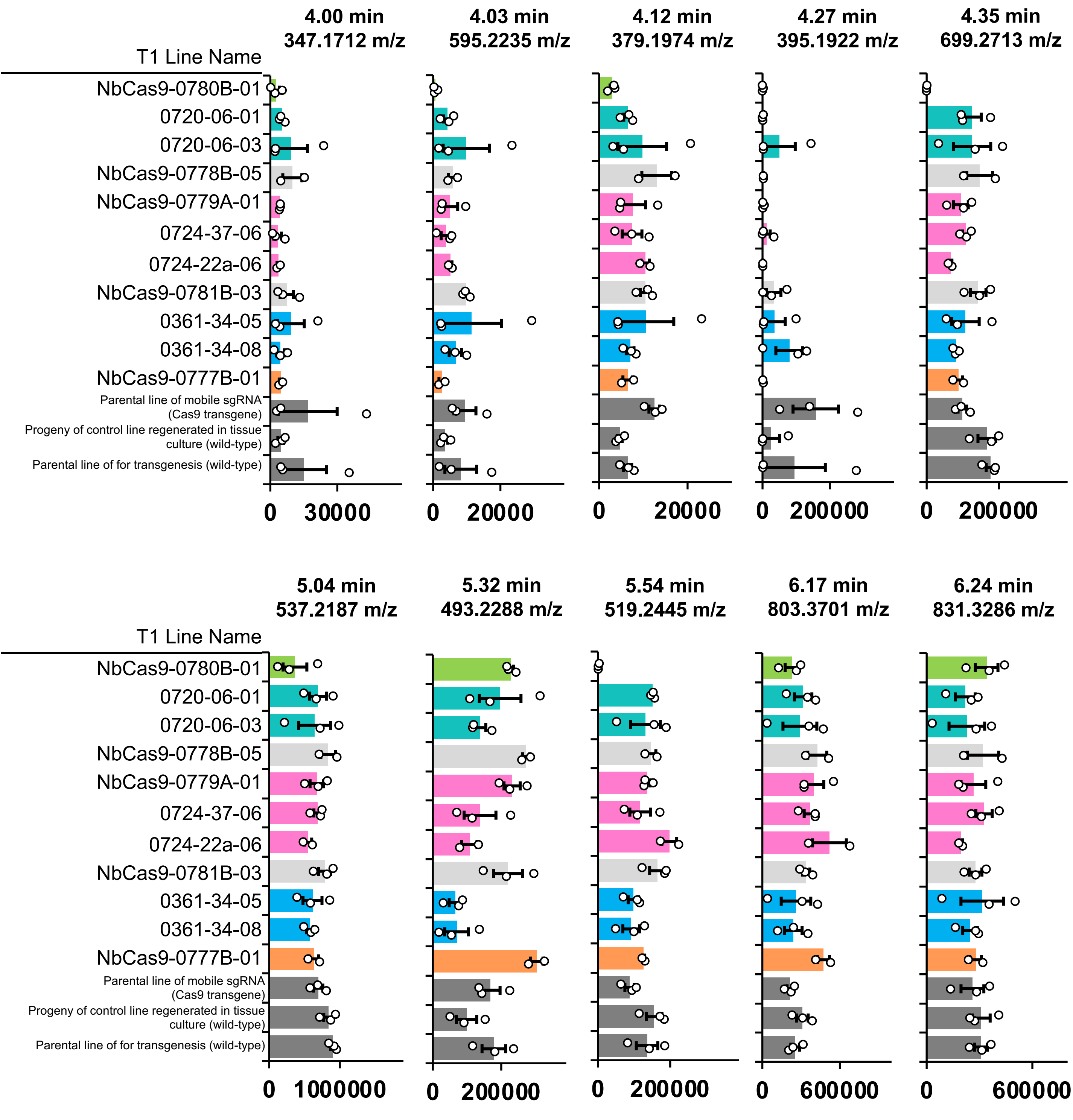


**Supplementary Figure S7 continued** Abundance of derivative peaks in lines of *N. benthamiana* infiltrated with the geraniol biosynthesis pathway (P19 + **DXS** + **GPPS** + **GES**)

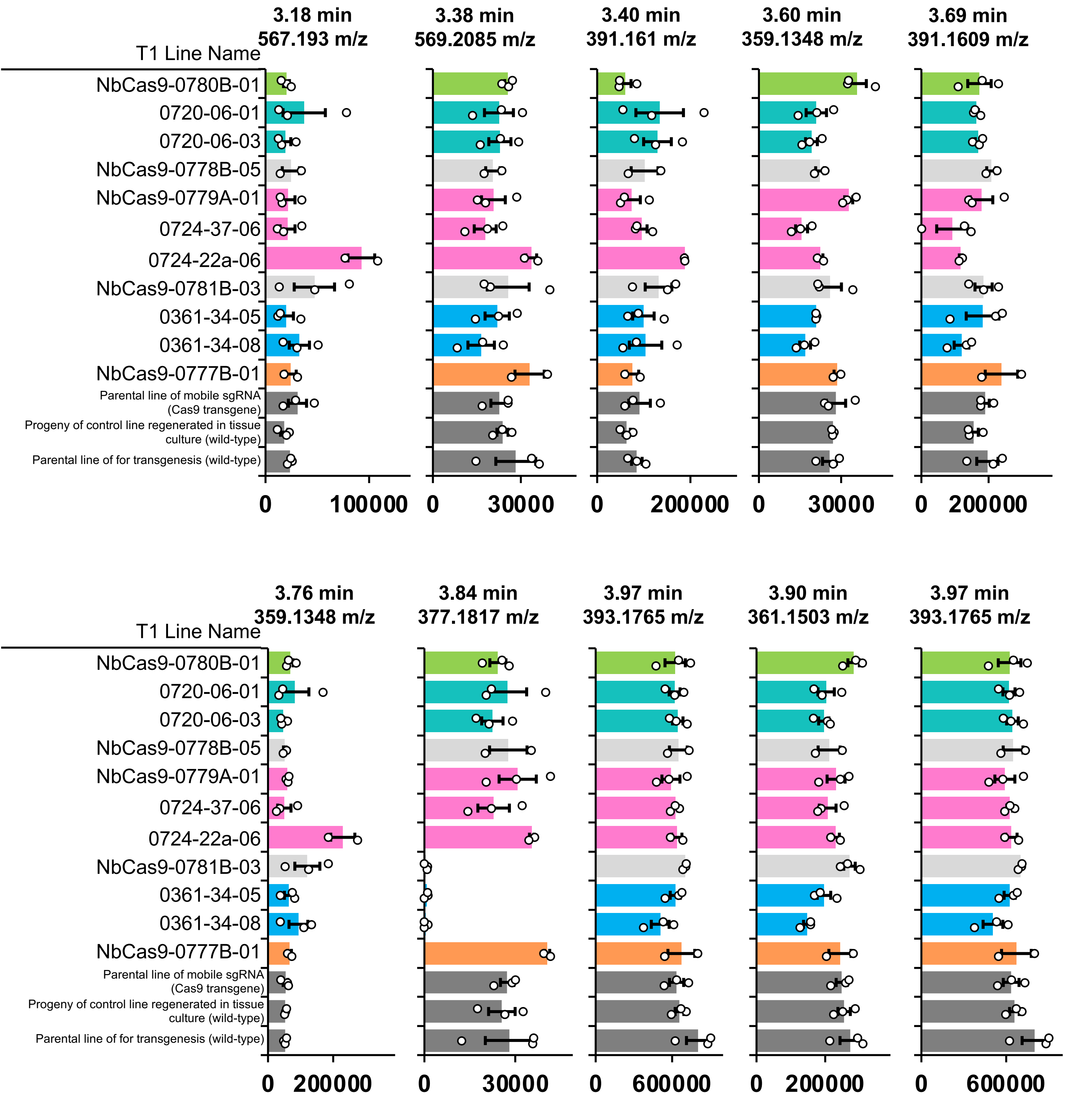


**Supplementary Figure S8** Abundance of derivative peaks in lines of *N. benthamiana* infiltrated with the nepetalactol biosynthesis pathway (P19 + **DXS** + **GPPS** + **GES** + **G8H** + **GOR** + **ISY + MLPL**)

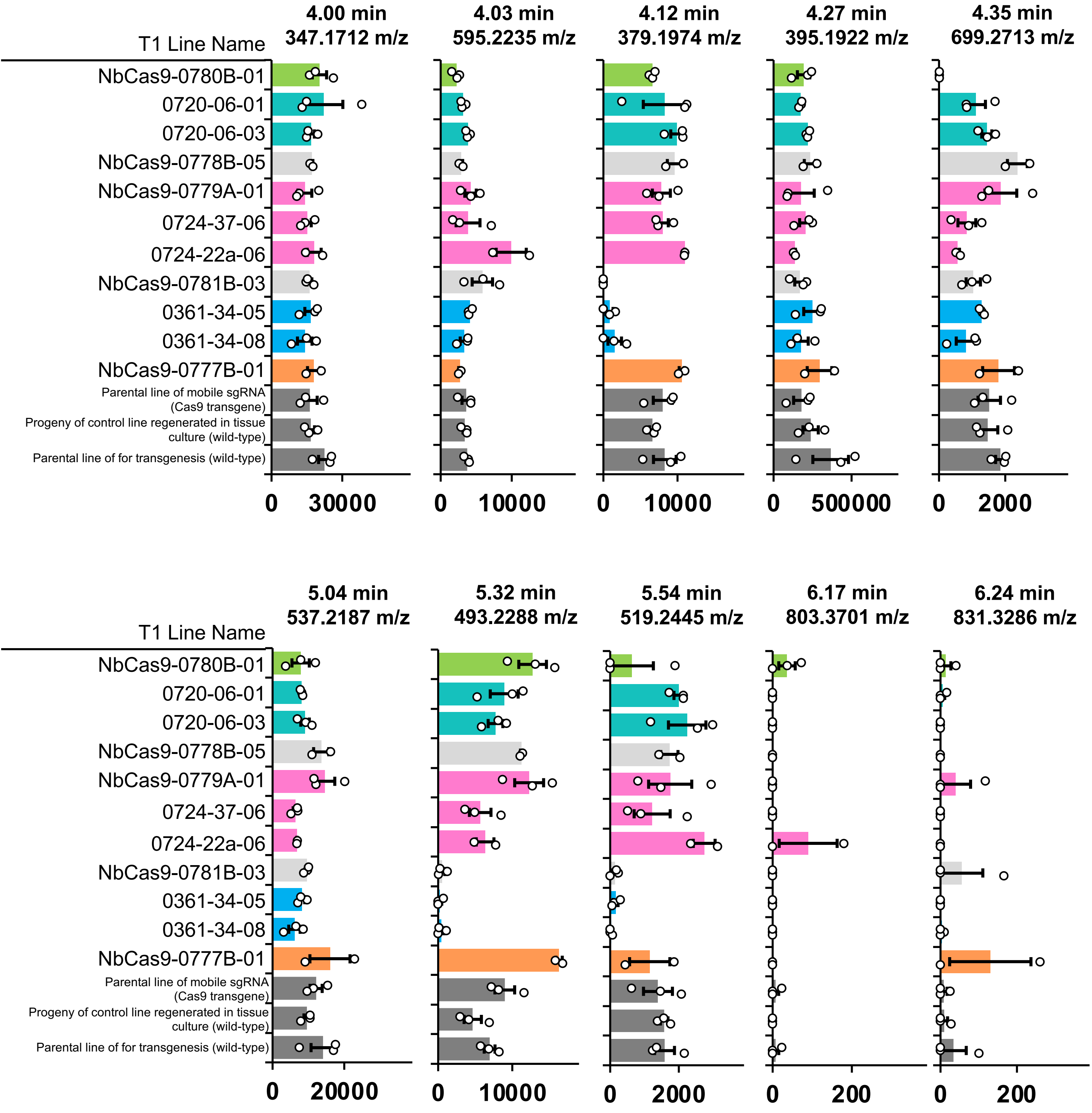


**Supplementary Figure S8 continued** Abundance of derivative peaks in lines of *N. benthamiana* infiltrated with the nepetalactol biosynthesis pathway (P19 + **DXS** + **GPPS** + **GES** + **G8H** + **GOR** + **ISY + MLPL**)

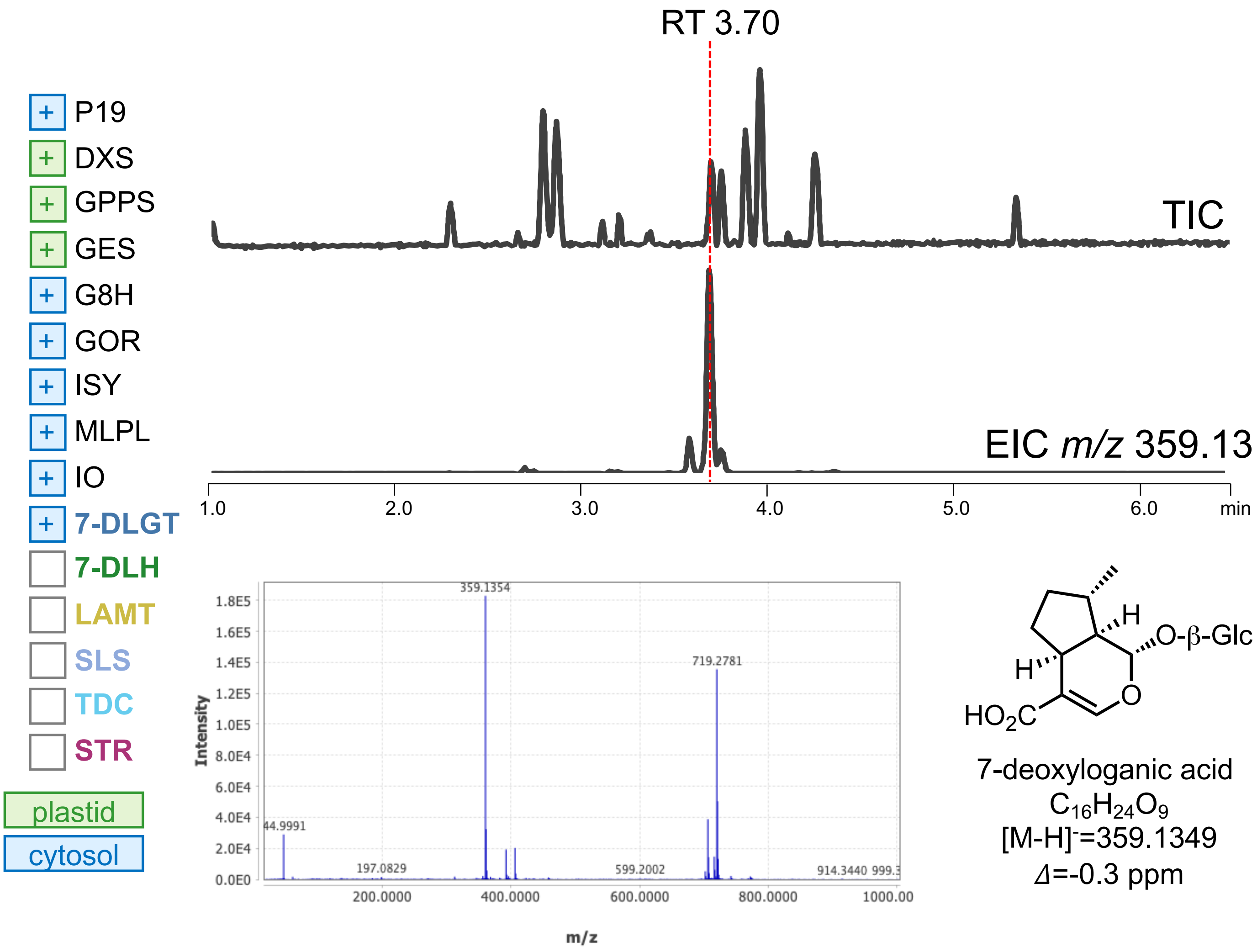


**Supplementary Figure S9**. Detection of 7-deoxyloganic acid following transient expression of pathway genes in *N. benthamiana*

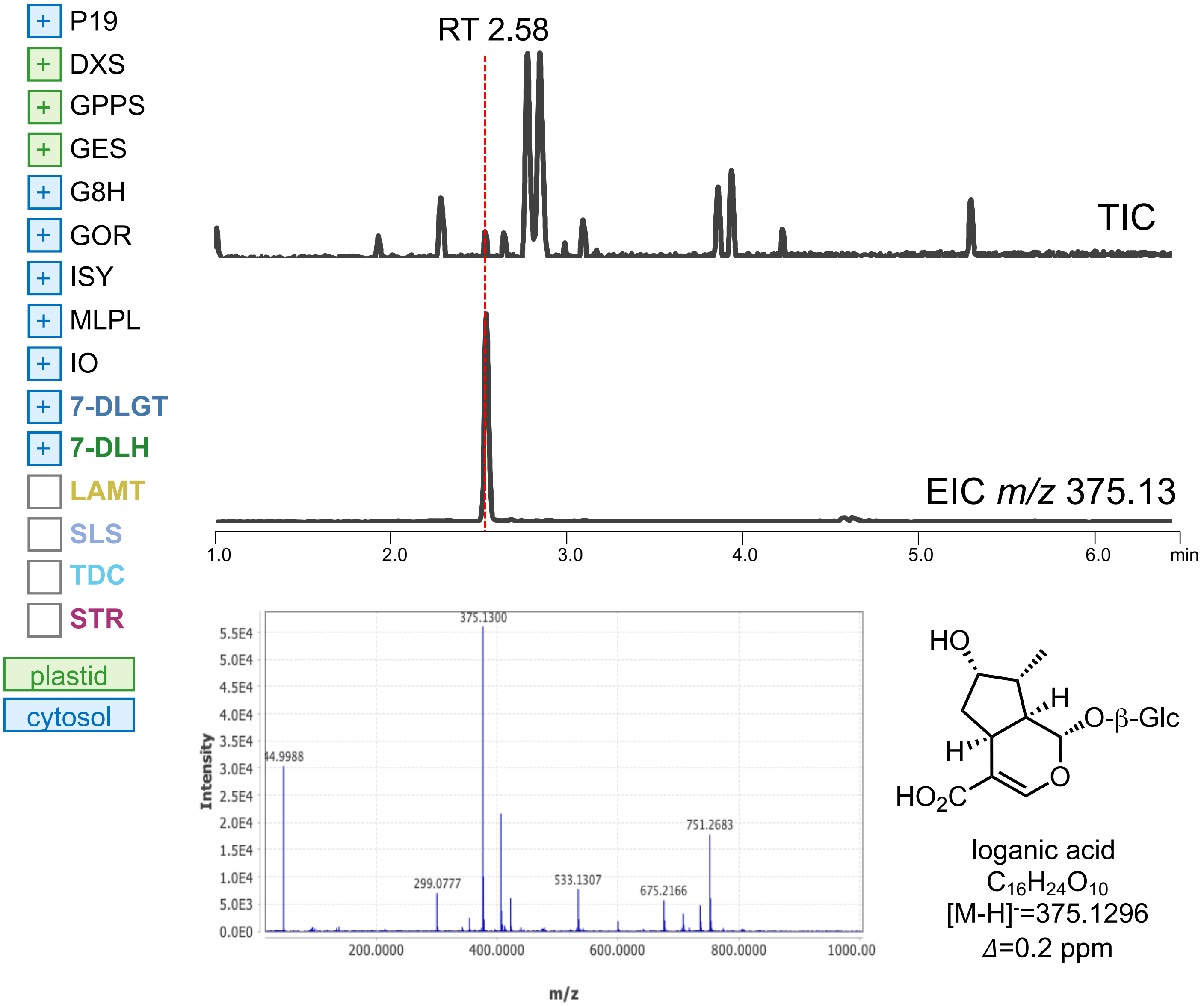


**Supplementary Figure S10**. Detection of loganic acid following transient expression of pathway genes in *N. benthamiana*

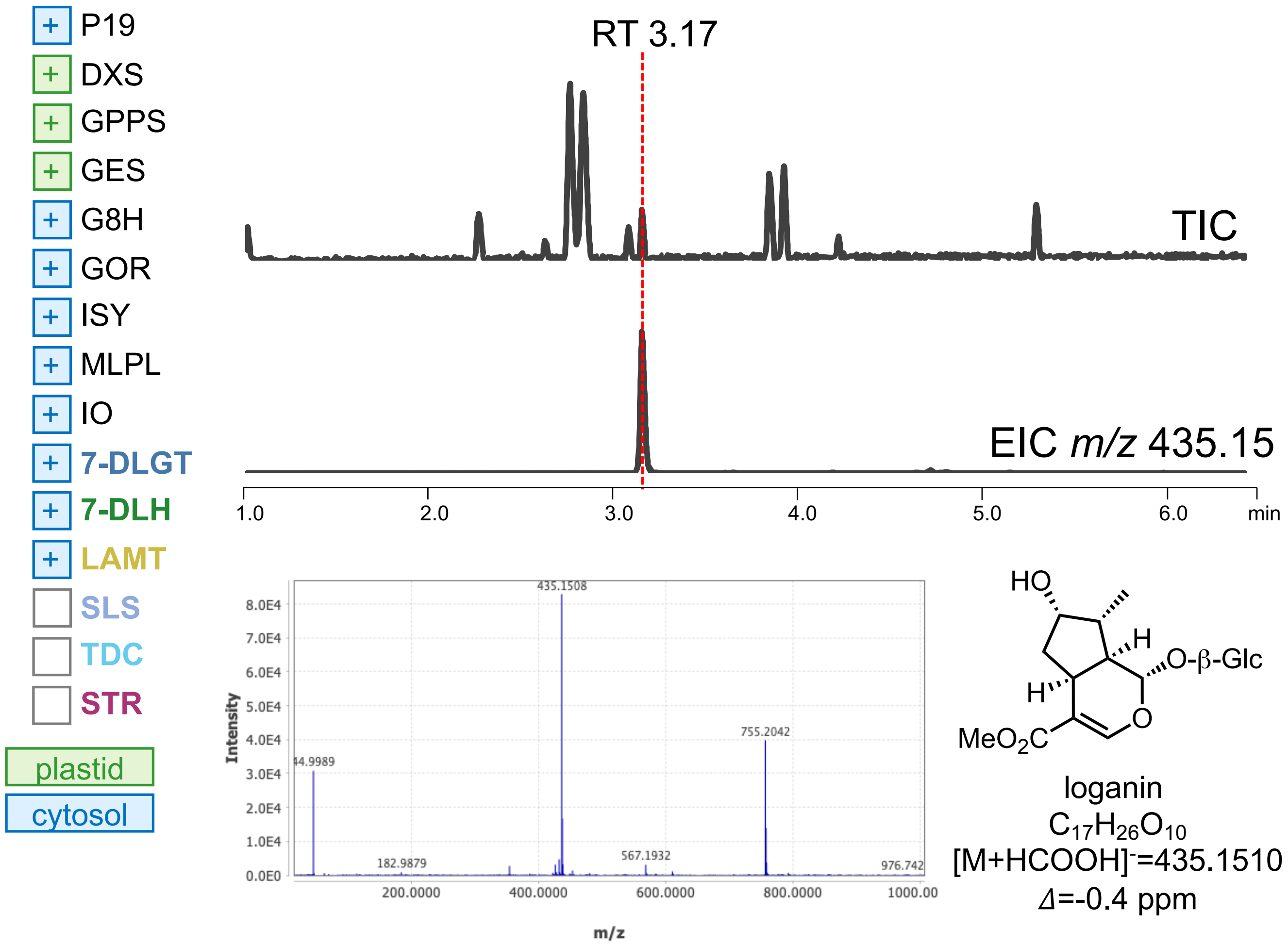


**Supplementary Figure S11**. Detection of loganin following transient expression of pathway genes in *N. benthamiana*

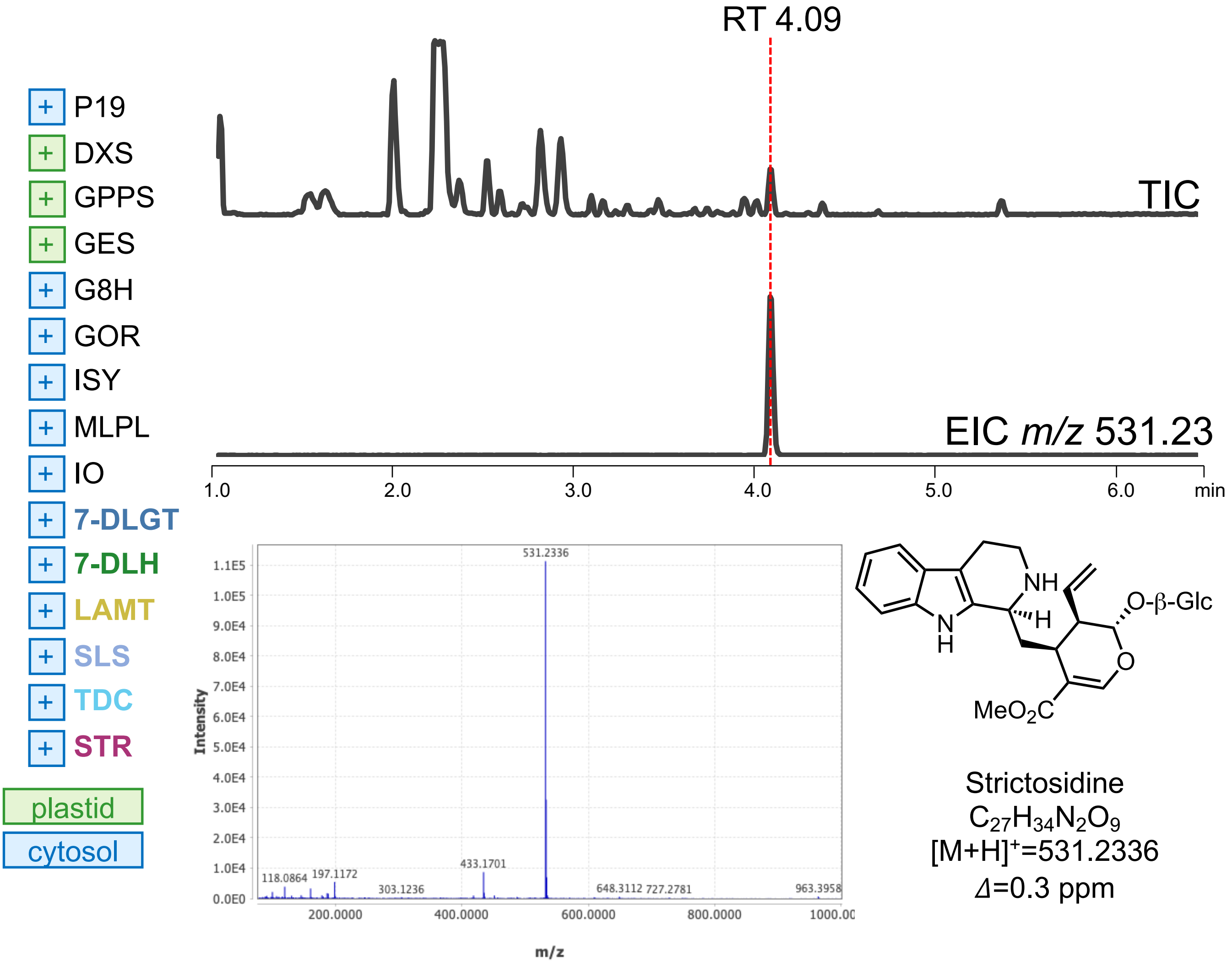


**Figure S12.** Detection of strictosidine following transient following transient expression of pathway genes in *N. benthamiana*

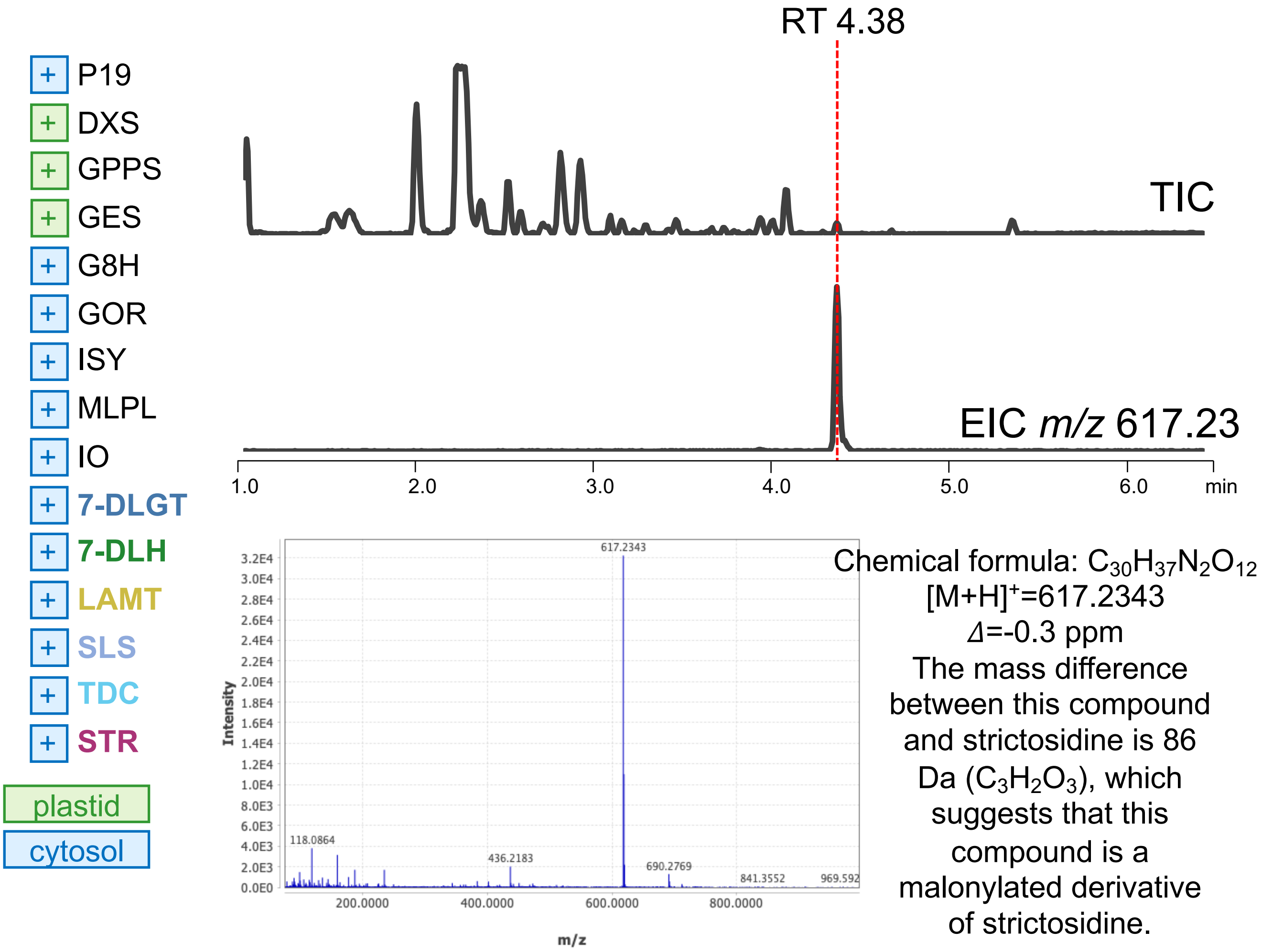


**Figure S13.** Detection of putative malonyl-strictosidine following transient expression of pathway genes in *N. benthamiana*

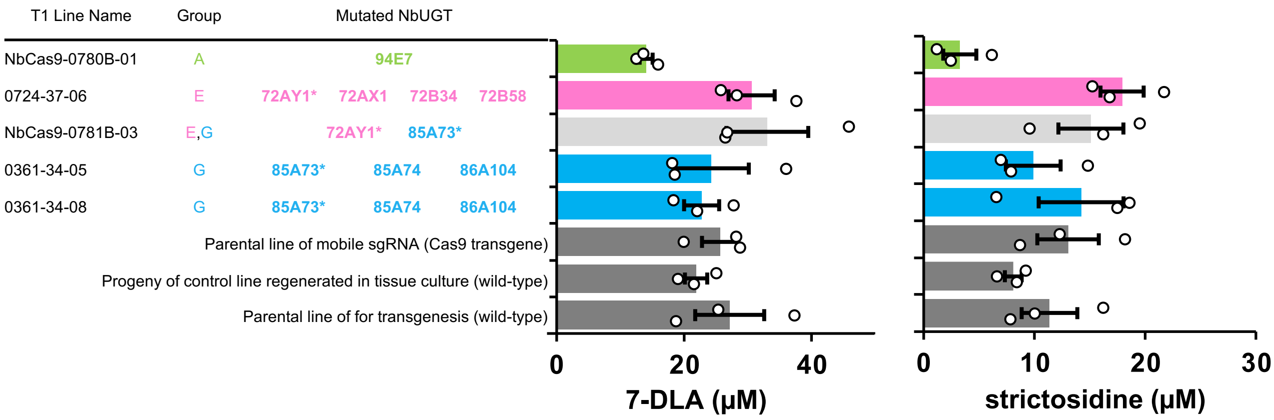


**Figure S14.** Quantification of strictosidine following transient expression of pathway genes in mutated and wild-type *N. benthamiana* by UPLC/MS analysis

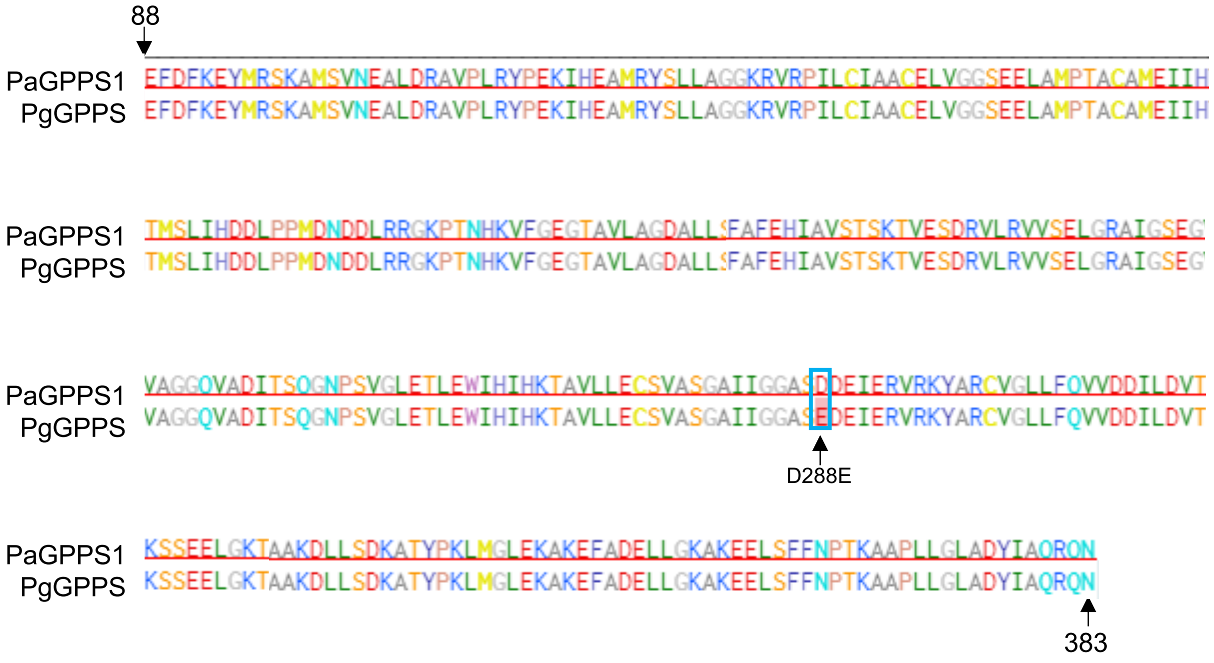


**Supplementary Figure S15.** Amino acid sequence alignment of GPPS enzymes. PaGPPS1 (GQ369788.1) - Used in this study and in Miettinen *et al. Nat. Comm.* 5:3606 (2014), and Dong *et al. New Phytologist* 209:2 679-690 (2016). PgGPPS (AHE15048.1) shown by Dudley *et al. Metab. Eng.* 61 251-260 (2020) to produce higher levels of the monoterpene limonene relative to six other GPPS sequences commonly used in terpene metabolic engineering.
